# BioSkills Guide: Development and National Validation of a Tool for Interpreting the Vision and Change Core Competencies

**DOI:** 10.1101/2020.01.11.902882

**Authors:** Alexa W Clemmons, Jerry Timbrook, Jon C Herron, Alison J Crowe

## Abstract

To excel in modern STEM careers, biology majors need a range of transferrable skills, yet competency development is often a relatively underdeveloped facet of the undergraduate curriculum. Here, we have elaborated the Vision and Change core competency framework into a resource called the BioSkills Guide, a set of measurable learning outcomes that can be more readily interpreted and implemented by faculty. College biology educators representing over 250 institutions, including 73 community colleges, contributed to the development and validation of the guide. Our grassroots approach during the development phase engaged over 200 educators over the course of five iterative rounds of review and revision. We then gathered evidence of the BioSkills Guide’s content validity using a national survey of over 400 educators. Across the 77 outcomes in the final draft, rates of respondent support for outcomes were high (74.3% - 99.6%). Our national sample included college biology educators across a range of course levels, subdisciplines of biology, and institution types. We envision the BioSkills Guide supporting a variety of applications in undergraduate biology, including backward design of individual lessons and courses, competency assessment development, curriculum mapping and planning, and resource development for less well-defined competencies.

## INTRODUCTION

Undergraduate biology students pursue a wide variety of career paths. Approximately 46% of undergraduates majoring in life sciences-related fields go on to STEM or STEM-related occupations, including research, engineering, management, and healthcare (Landivar, 2013). The over half of life science majors employed outside of STEM can be found in non-STEM related management, business, and K-12 education, among many other positions. Considering that the majority of college students and the general public indicate career success as the primary motivation for attending college (Pew Research Center, 2016; Strada Education Network, 2018; Twenge & Donnelly, 2016), it follows that undergraduate biology curricula should include competencies that will help students thrive in their post-college pursuits, in or out of STEM.

Employers across fields routinely rank competencies such as collaboration, communication, and problem-solving at the top of the list of desirable employee traits (NACE, 2018; Strauss, 2017), and also report that new hires are not adequately trained in these areas (Bayer Corporation, 2014; Hart Research Associates, 2018). While ‘skills gap’ rhetoric and the associated vocational framing of higher education has been criticized (Camilli & Hira, 2019; Cappelli, 2015), college courses are nonetheless a natural environment for competency development because of the opportunities to practice skills in the context of relevant knowledge and receive formative feedback from disciplinary experts (Hora, 2018).

### Competencies and STEM Curriculum Reform

Numerous national reports have pushed educators to reexamine how competencies are integrated into undergraduate STEM coursework (NASEM, 2016; NRC, 2003, 2012b). In undergraduate biology, these recommendations are presented in the report “Vision and Change in Undergraduate Biology Education: A Call to Action” (AAAS, 2011). The recommendations of Vision and Change emerged from discussions among over 500 stakeholders in undergraduate biology education, including educators, administrators, students, scientists, and education researchers. To prepare students for modern careers, the report urges biology educators to frame discussions of curricula around five core concepts and six core competencies (listed in Table 1).

**Table 1.** Comparison of Vision and Change in Undergraduate Biology Education core competencies (AAAS, 2011) and Framework for K-12 Science Education scientific practices (NRC, 2012a). ^a^This scientific practice aligns with two of the core competencies. ^b^Conceptions of what models are and how they are used are not well defined in Vision and Change and thus may differ from the scientific practice presented in the K-12 Framework. ^c^Crosscutting concepts is a separate dimension of the 3D K-12 Framework, not a scientific practice.

| Vision and Change Core Competencies | K-12 Framework Scientific Practices |
| --- | --- |
| Ability to apply the process of science | Asking questions<br>Analyzing and interpreting data<br>Planning and carrying out investigations<br>Engaging in argument from evidence<br>Obtaining, evaluating, and communicating information<sup>a</sup> |
| Ability to use quantitative reasoning | Using mathematics and computational thinking |
| Ability to use modeling and simulation<sup>b</sup> | Developing and using models |
| Ability to tap into the interdisciplinary nature of science | Crosscutting concepts<sup>c</sup> |
| Ability to communicate and collaborate with other disciplines | Obtaining, evaluating, and communicating information<sup>a</sup> |
| Ability to understand the relationship between science and society |  |
|  | Constructing explanations |

The publication of Vision and Change in 2011 coincided temporally with several similar efforts to guide STEM curriculum reform. The updated AP Biology Curriculum Framework emphasized science practices (College Board, 2015). Foundations for Future Physicians advised premedical and medical educators away from curriculum based on lists of courses and toward the measurement of scientific competencies (AAMC & HHMI, 2009). The NRC’s Framework for K-12 Science Education advocated for the “three-dimensional” integration of disciplinary core ideas, crosscutting concepts, and scientific practices (NRC, 2012a). The K-12 Framework’s approach to elementary and secondary science education aimed to improve science literacy in the population as a whole by better engaging students in authentic scientific experiences. Since its publication, the K-12 Framework has emerged as the consensus framework for developing K-12 science curricula, and has since been enumerated into the Next Generation Science Standards (NGSS) (*Next Gener. Sci. Stand. States, By States*, 2013).

In comparing the Vision and Change core competencies with the K-12 Framework scientific practices, we find a few notable differences (Table 1). Whereas Vision and Change explicitly includes the ability to collaborate and to understand the relationship between science and society, these practices are not directly called out in the K-12 Framework. Similarly, while the K-12 Framework specifically highlights the ability of students to construct explanations, this practice is only implicitly included in Vision and Change within the core competency of Process of Science. However, taken as a whole, the overlap between the core competencies and scientific practices is substantial (Table 1). The parallel evolution of K-12 and undergraduate curricular goals represents an opportunity to cohesively improve educational outcomes and is an area that deserves continued attention to ensure a smooth transition from high school to college.

The development of the Vision and Change curricular recommendations was an important milestone in undergraduate biology education. By bringing together biologists and biology education experts to reimagine the curriculum, the resulting recommendations were specifically tailored to undergraduate biology but with substantial overlap with related educational efforts. Furthermore, the resulting concepts and competencies provided a common goal, written in the language of biology educators, promoting buy-in. As such, the Vision and Change curricular framework has been widely embraced by the undergraduate biology community (AAAS, 2015, 2018, 2019; “About CourseSource,” n.d.; Brancaccio-Taras et al., 2016; Dirks & Knight, 2016). However, because the report’s descriptions of the core concepts and competencies were left intentionally brief to encourage ongoing conversations among educators, they require elaboration in order to be implemented. Since then, two groups have unpacked the core concepts into more detailed frameworks (Brownell, Freeman, Wenderoth, & Crowe, 2014; Cary & Branchaw, 2017).

For competencies, biology education researchers have enumerated a variety of specific scientific practices, including: science process skills (Coil, Wenderoth, Cunningham, & Dirks, 2010), experimentation (Pelaez et al., 2017), scientific literacy (Gormally, Brickman, & Lutz, 2012), responsible conduct of research (Diaz-Martinez et al., 2019), quantitative reasoning (Durán & Marshall, 2018; Stanhope et al., 2017), bioinformatics (Wilson Sayres et al., 2018), data science (Kjelvik & Schultheis, 2019), data communication (Angra & Gardner, 2016), modeling (Diaz Eaton et al., 2019; Quillin & Thomas, 2015), the interdisciplinary nature of science (Tripp & Shortlidge, 2019), and scientific writing (Timmerman, Strickland, Johnson, & Payne, 2011). Efforts to define general or STEM-wide education goals for college graduates can also inform how we teach competencies in biology, such as the Association of American College & University VALUE rubrics (Rhodes, 2010) and more targeted work on information literacy (Association of College and Research Libraries, 2015), communication (Mercer-Mapstone & Kuchel, 2017), and process skills (Cole, Lantz, Ruder, Reynders, & Stanford, 2018; Understanding Science, 2016). However, no resource has yet been developed that *holistically* considers competencies across college biology programs or that is intentionally aligned with the recommendations of Vision and Change.

### Project Goals and Context

With the overarching goal of improving biology undergraduates’ achievement of competencies relevant to their careers and life as scientifically literate citizens, we set out to expand the six Vision and Change core competencies into measurable learning outcomes that describe what a general biology major should be able to *do* by the time they graduate. The intention of this work is to establish competency learning outcomes that:

1. define what each of the broadly stated competencies means for an undergraduate biology major, especially for less commonly discussed competencies such as Modeling and Interdisciplinary Nature of Science,
2. draw on instructor expertise to calibrate an appropriate level of competency that can be achieved over the course of a 4-year biology program,
3. serve as a starting point for backward design of individual courses or departmental programs, and
4. ease interpretation and, therefore, adoption of the Vision and Change core competencies in undergraduate college curricula.

The term “competency” describes a “blend of content knowledge and related skills” (NRC, 2012b) and is thus appropriate for describing complex tasks like modeling biological systems or understanding the interrelatedness of science and society. The term “scientific practice” is employed similarly in the Framework for K-12 Science Education (NRC, 2012a). However, throughout the development of this resource through workshops, round tables, and informal conversations we found that the term “skill” was more immediately recognizable (to biology educators not engaging in discipline-based education research) and less frequently unintentionally confused with the term “concept” (especially when talking about “concepts and competencies”). While it should be noted that use of the term “skill” can connote a simplified behaviorist framing of science education (e.g., teacher-centered practice and rote memorization via repetitive drills) (Agarkar & Brock, 2017), we did not find this implied definition to be held among our sample of biology educators. Instead, we found that the term “skills” was understood to refer to a broad set of competencies performed within a biological context. For the purpose of this study, we have therefore used the term “skills” interchangeably with competencies and have named the resource we developed the “BioSkills Guide”.

We describe here the iterative, mixed-methods approach we used to develop and establish content validity of the BioSkills Guide. Evidence of content validity was collected via a national survey of college biology educators, a population we selected based on their combined expertise in biology and biology education. This population includes many researchers engaged in discipline-based education research, and thus brings that expertise as well. We chose to focus on this population as they are the intended users of the guide. Institutional change has been shown to be most effective when the work is envisioned and led by those directly impacted by the change (Henderson, 2010). A similar grassroots approach was used to develop Vision and Change itself, as well as related frameworks elaborating the core concepts (Brownell, Freeman, et al., 2014; and Cary & Branchaw, 2017), which have been widely utilized in our field (Branchaw et al., 2020; Smith et al., 2019). We believe this approach is another reason why Vision and Change has been so impactful in biology education.

Specifically, we asked the following research questions (RQs):

RQ1a: Can we identify an essential set of learning outcomes aligned with the Vision and Change core competencies?

RQ1b: How much do biology educators agree on this essential set of competency learning outcomes?

RQ2a: Does biology educators’ support of learning outcomes differ across competencies?

RQ2b: Do biology educators with different professional backgrounds differ in their support of learning outcomes across competencies?

The final draft of the BioSkills Guide contains 77 measurable learning outcomes (20 program-level and 57 course-level) that elaborate the six Vision and Change core competencies. Both the BioSkills Guide and an “expanded BioSkills Guide”, which contains illustrative examples of activities intended to support student mastery of the learning outcomes, are available in Supplemental Materials. The BioSkills Guide is also available at https://qubeshub.org/qubesresources/publications/1305.

## METHODS

This work can be divided into two phases: a constructive development phase (RQ1a) and an evaluative validation phase (RQ1b; the phases are summarized in Figure 1). During the development phase we used a range of methods to gather biology education community feedback on sequential drafts of the BioSkills Guide: web surveys, unstructured and semi-structured interviews, workshops, and round tables (Table 2). During the validation phase we used a web survey to measure support for the final draft among the broader biology education community. We then applied the validation phase survey data to answer RQ2a and 2b. This study was approved by the University of Washington, Human Subjects Division as exempt (STUDY00001746).

**Table 2.**
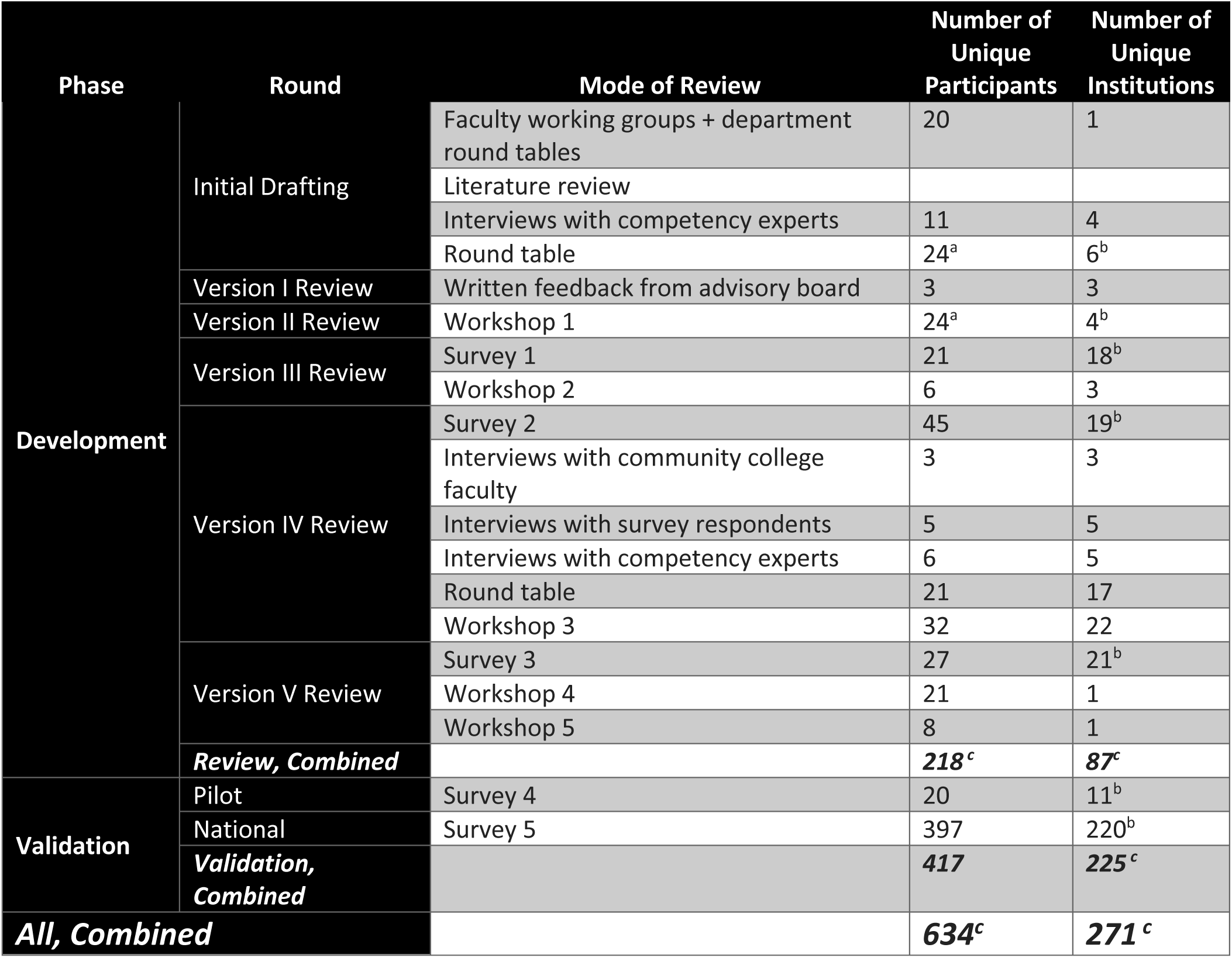
Unique participants and institutions during BioSkills Guide development and validation. ^**a**^ Number of participants is an underestimation because not all participants completed sign-in sheet. ^b^ Number of institutions is an underestimation because institution is unknown for some participants. ^c^ Number of total participants is a conservative estimation due to missing information as described in ^a^ and ^b^. Number is lower than the sum of above rows because a small percent of people participated at multiple stages, which has been accounted for where possible (e.g., known participants were only counted once; anonymous survey respondents indicating they had previously reviewed the BioSkills Guide were deducted from the total).

**Figure 1.**
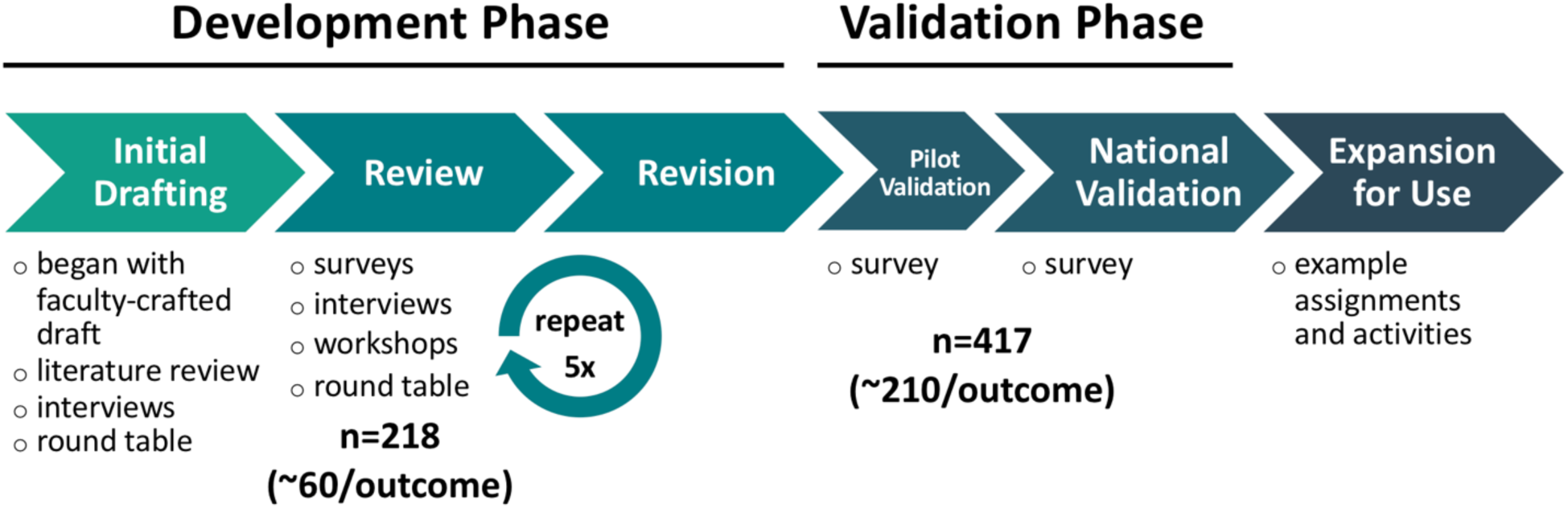
BioSkills Guide methods overview. Initial drafting included all work to generate BioSkills Guide Version I. Five rounds of review and revision were carried out on Versions I-V (RQ1a). Pilot validation evaluated Version VI (RQ1b). National validation evaluated final version of BioSkills Guide (RQ1b).

### Development Phase

To address RQ1a, we developed the initial draft of the BioSkills Guide by building on a set of programmatic learning outcomes crafted by biology faculty at a large, public research university in the Northwest as part of routine departmental curriculum review. We supplemented the initial draft by cross-checking its content with the literature, leading unstructured interviews with competency experts, and gathering feedback on a portion of the draft at a round table at a national biology education conference (additional details in Supplemental Methods).

We next began the first of five rounds of review and revision of iterative drafts of the learning outcomes (Table 2). First, we collected feedback on Version I of the outcomes in writing and via a virtual meeting with our advisory board (three biology faculty with expertise in institutional change, programmatic assessment, and/or curricular framework development). To review Version II of the guide, we collected written feedback on outcome importance, clarity, and completeness at a workshop of biology faculty, postdocs, and graduate students. The final three rounds were larger in scale, and each included a survey to gather feedback on outcome importance, ease of understanding, completeness, and categorization from at least 21 college biology educators (5-19 per learning outcome per round) (Table 2, Supplemental Table 4). We recruited respondents at regional and national biology education meetings and through regional biology education networks. In order to participate in any of the surveys, respondents must have served as instructor-of-record of a college-level biology course. We chose this inclusion criterion because college biology instructors have expertise in both biology and biology education. Many respondents also had discipline-based education research experience (48.4% during development phase). We gathered additional input on Version III-V drafts using four workshops, one round table, and 14 one-on-one interviews. Additional details on BioSkills Guide development are in Supplemental Methods.

At the end of each round of review, we compiled and summarized all relevant data (i.e. data from workshops, interviews, round tables, or surveys) from that round into a single document to inform revisions. This document was then reviewed by committee (two authors, AWC and AJC, for Versions I-III revisions; three authors, AWC, AJC and JCH, for Versions IV-V revisions) and used to collectively decide on revisions. The committee discussed all revisions and their justifications over the course of several meetings per round, revisiting relevant feedback from previous rounds as necessary.

During revisions, we reworded outcomes based on feedback to ensure they were easy to understand, calibrated to the right level of challenge for an undergraduate program, and widely relevant to a variety of biology subdisciplines, institution types, and course levels (Supplemental Table 1). New outcomes were considered for addition if they were suggested by more than one participant. We removed outcomes only after multiple rounds of negative feedback despite revisions to improve ease of reading or possible concerns about challenge level or relevance. We did not have an *a priori* quantitative threshold for survey ratings to determine whether to retain outcomes, however we critically evaluated any outcomes that had lower than 90% ratings of ‘Important’ or ‘Very Important’ by reviewing qualitative feedback from survey comments, interviews, and workshops. This process resulted in the removal of 21 outcomes total (ranging from 50-88% survey ratings of ‘Important’ or ‘Very Important’, with an average of 73.5%) over the course of 5 rounds of review (Supplemental Table 1). Occasionally outcomes were removed despite having higher quantitative support than other outcomes that were retained, due to qualitative feedback such as that the outcome had substantial overlap with other outcomes, was too specialized or at too high of a challenge level for an undergraduate general biology major, or could not be readily assessed. In general, we identified problems in the drafts by looking at outcomes that had low ratings or low consensus (e.g., a mixture of low and high ratings). We then used qualitative feedback from survey comments, workshops, round tables and interviews to inform revisions.

### Validation Phase

To address RQ1b, we next sought to gather evidence of content validity of the final draft via a survey of college biology educators. Before proceeding with a national survey, however, we first conducted a “pilot” validation on a smaller pool of educators (n=20). After reviewing the results, we revised one outcome: “Identify methodological problems and suggest alternative approaches or solutions”. The previous revision of this outcome had reworded it to use language that was appropriate for a wide range of study types (not just experiments) and had removed the word “troubleshooting”. We speculated that this term had resounded with respondents and thus led to previously observed greater levels of support, so we revised the outcome to reintroduce it. This was the only revision to the guide before moving on to the large-scale national validation (Supplemental Table 1). Additional details on the pilot validation can be found in Supplemental Methods.

For national validation, we invited participation through direct emails and numerous listservs: Society for Advancement of Biology Education Research (SABER), Partnership for Undergraduate Life Sciences Education (PULSE) regional networks, HHMI Summer Institutes, authors of CourseSource articles tagged with “Science Process Skills”, Community College BioInsites, Northwest Biology Instructors Organization, the Science Education Partnership and Assessment Laboratory network, Human Anatomy and Physiology Society, SABER Physiology Special Interest Group, several other regional biology education-related networks, and 38 participants suggested by previous survey participants. We additionally encouraged advisory board members, other collaborators, and survey respondents to share the survey invitation widely. Because of the snowball sampling approach and the expected overlap of many of the email lists, it is not possible to estimate the total number of people who were invited to participate. To participate in the survey, respondents had to meet the same survey inclusion criterion (i.e. having taught a college biology course) as during the development phase.

For RQ1b analysis, we combined data from the pilot validation and national validation surveys. 572 people initiated the validation phase surveys (21 for pilot validation, 551 for national validation). Of those, 22 people did not meet our survey inclusion criterion and 133 people did not respond to any questions after the initial screening question (i.e. did not rate any learning outcomes) and so could not be included in our analysis. It is possible that some of these 133 individuals started the survey on one device (e.g., home computer, mobile phone) and later restarted and completed the survey using a different device (e.g., work computer), thus some of these 133 instances may include individuals that ultimately responded to the survey. We do not have demographic data (e.g., institution type, familiarity with Vision & Change) for these 133 instances and therefore cannot assess whether these individuals differed on demographic characteristics compared to those who did rate at least one learning outcome. Ultimately, responses from 417 people were retained for the analysis for RQ1b (572-22-133=417; total responses per outcome ranged from 211 to 237).

One minor modification was made in the BioSkills Guide after national validation. The Modeling learning outcome “Build and revise conceptual models (e.g., diagrams, concept maps, flow charts) to propose how a biological system or process works.” was revised to remove the parenthetical list of examples. We made this revision based on post-validation feedback from modeling experts, among whom there was disagreement as to whether visual representations such as diagrams and concept maps constitute conceptual models. To avoid confusion, we removed the examples. No other revisions were made to the learning outcomes after the national validation survey (Supplemental Table 1).

### Survey Design

As mentioned above, we employed five surveys over the course of this project (three in the development phase and two in the validation phase; Table 2). Surveys were designed and administered following best practices in survey design and the principles of social exchange theory (Dillman, Smyth, & Christian, 2014). For development phase surveys, respondents rated each learning outcome on bipolar 5-point Likert scales for: (1) how important or unimportant it is for a graduating general biology major to achieve (‘Very Important’, ‘Important’, ‘Neither Important nor Unimportant’, ‘Unimportant’, and ‘Very Unimportant’), and (2) how easy or difficult it is for them to understand (‘Very Easy’, ‘Easy’, ‘Neither Easy nor Difficult’, ‘Difficult’, ‘Very Difficult’). We also asked respondents to comment on their responses, suggest missing outcomes, and evaluate (yes/no) whether each learning outcome was accurately categorized within its program-level outcome (when evaluating course-level outcomes) or competency (when evaluating program-level outcomes). For validation phase surveys, we shortened the questionnaire by removing the items on ease of understanding and categorization and by reducing the frequency of questions that asked respondents to comment on their responses. To minimize time commitments and thus maximize survey responses, we asked respondents to review outcomes associated with only two (during development phase) or three (during validation phase) randomly assigned competencies, with the option to review up to all six competencies. We collected respondent demographic information for all surveys. See Supplemental Tables 2 and 6 for a summary of demographic information collected. The entire questionnaires for Version V review and national validation can be found in Supplemental Materials.

### Descriptive Analysis of Survey Responses

To address RQ1a and 1b, we calculated and visualized descriptive statistics of survey responses and respondent demographics in R version 3.5.1 (R Core Team, 2018) using the tidyverse, ggmap, maps, ggthemes, ggpubr, and wesanderson packages (Arnold, 2019; Kahle & Wickham, 2013; Kassambara, 2018; Ram & Wickham, 2018; Wickham, 2016). For importance and ease of understanding responses, we calculated the mean, minimum, and maximum ratings (where 5 = ‘Very Important’ or ‘Very Easy’ and 1 = ‘Very Unimportant’ or ‘Very Difficult’). We binned responses of ‘Very Important’ or ‘Important’ as ‘Support’, and calculated ‘Percent Support’ as the percent of respondents who ‘Supported’ the outcome out of all respondents who reviewed that outcome. We calculated the percent of respondents who selected ‘Very Easy’ or ‘Easy’ out of all respondents who reviewed that outcome (development phase only). We calculated the percent of respondents who indicated that the outcome was accurately categorized within its competency or program-level learning outcome (development phase only, not shown). We read and summarized the open-ended comments to inform revisions (development phase) or to summarize suggestions of missing outcomes (validation phase). We summarized responses to demographic questions by calculating the frequency and percent of respondents who selected different responses for each question. We determined the Carnegie classification of their institution type, minority-serving institution status, and geographic location by matching their institution name with the Carnegie dataset (Indiana University Center for Postsecondary Research, 2016). We then mapped participant locations using their institution’s city and state GPS coordinates, obtained via the Google API (Kahle & Wickham, 2013).

### Treatment of Missing Data for Statistical Modeling

To address RQ2a and RQ2b, we fit models of respondents’ support of learning outcomes using the competency of each outcome and respondents’ answers to end-of-survey demographic questions as predictors. Of our 417 initial respondents (i.e. respondents that rated at least one outcome) included in the RQ1b analysis, 71 did not provide all five demographic characteristics investigated in RQ2, and therefore were not included in these analyses. After removing these 71 individuals, our analytic dataset for RQ2 contained responses from 346 respondents, comprising 15,321 importance ratings across 77 learning outcomes. To ensure that these omissions did not bias our inference, we compared rates of outcome support (i.e. the dependent variable in our models) from the 71 individuals that were removed from the RQ2 analyses to the 346 individuals that were retained and found that rates of outcome support did not differ overall or by competency across the two groups (Supplemental Methods and Supplemental Table 9). As we did not have all demographic data on the 71 individuals removed from our RQ2 analyses, we cannot assess if demographic characteristics of the individuals we removed differed from the individuals that we retained.

As we randomly assigned respondents to rate outcomes for particular subset of competencies, all respondents did not rate all outcomes. Thus, the number of ratings per outcome in the RQ2 analytic dataset ranged from 183-206. When respondents were not assigned to rate outcomes from a particular competency, these data are missing completely at random (MCAR). The multilevel models we use in this study (described further below) allow for an unequal number of measurements across respondents in such cases (West, Welch, & Galecki, 2014). There were a few instances where respondents saw an outcome within their assigned competency but did not rate it (i.e. item nonresponse), but this behavior was rare (an average of 0.4% for each outcome). Our analyses do not include ratings on these missing outcomes, and this small amount of missing data is unlikely to bias our results (Graham, 2009).

### Statistical Models of Learning Outcome Ratings

In estimating models for RQ2a and 2b, we accounted for three key aspects of our data structure. First, each respondent rated multiple competencies and each competency contained multiple outcomes (refer to Supplemental Figure 1). We accounted for the non-independence in respondents and learning outcomes by fitting multilevel models with respondent and learning outcome as random effects (random intercepts) (Theobald, 2018). Second, by design each respondent rated learning outcomes corresponding to a random subset of competencies, so not all learning outcomes were evaluated by all respondents. To account for the imperfect nesting of responses within respondents and learning outcomes in our analyses, we used cross-classified multilevel models (Olson & Smyth, 2015; Yan & Tourangeau, 2008). Third, respondents rated importance on a 5-point Likert scale (from ‘Very Important’ to ‘Very Unimportant’), but the ratings for learning outcomes were generally very high (i.e. not normally distributed, see Figure 2 and Supplemental Table 4). We accounted for this skewed distribution by using the binary variable ‘Support’ (i.e. Support = 1 if the learning outcome was rated ‘Important’ or ‘Very Important’, otherwise Support = 0) as our dependent variable. Thus, we fit cross-classified multilevel binary logistic regression models (Raudenbush & Bryk, 2002) to address RQ2a and 2b. We estimated these models using the meqrlogit command in Stata (version 14.2).

**Figure 2.**
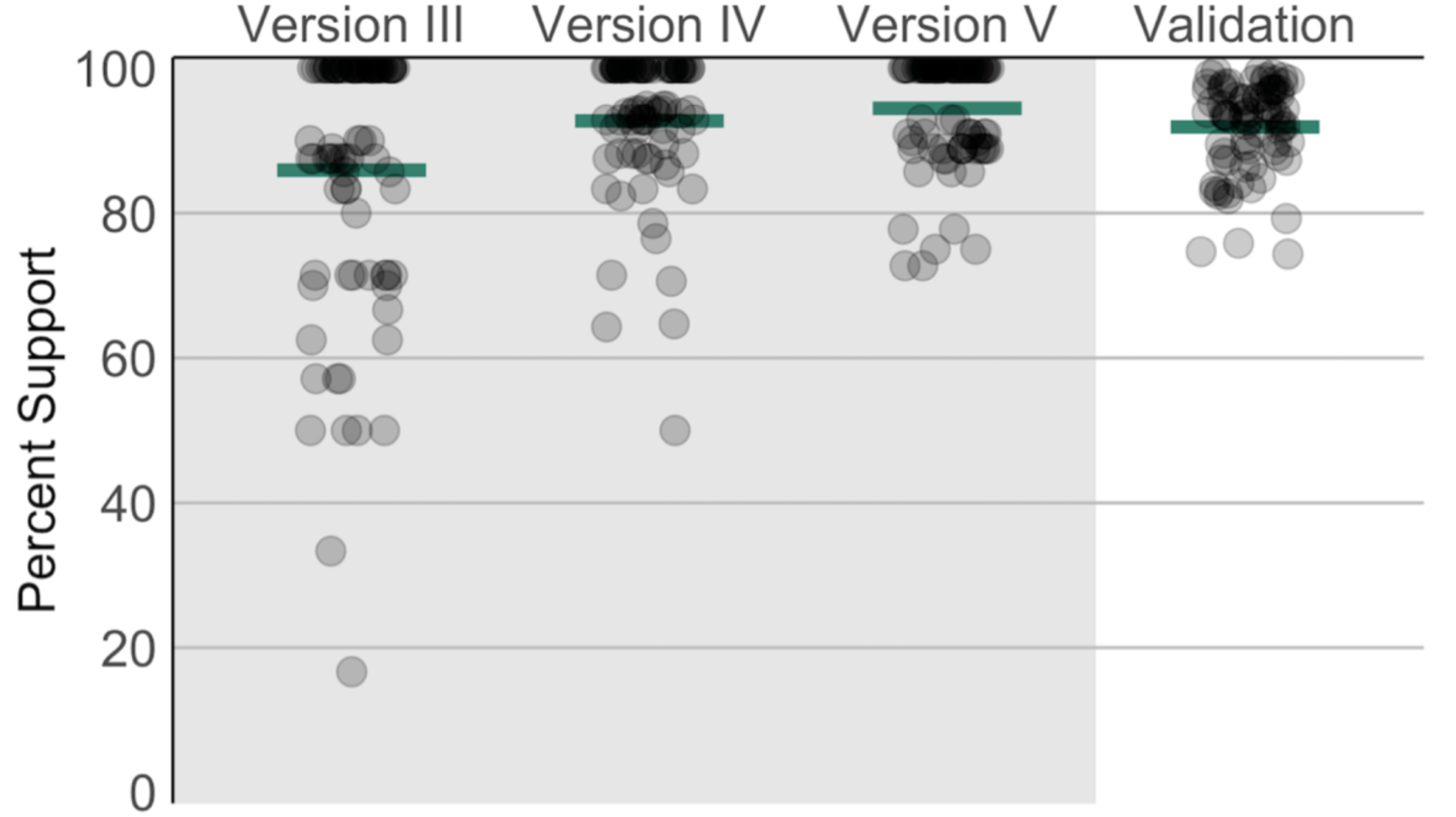
Learning outcome ratings show increasing consensus over iterative rounds of revision. Survey ratings were summarized by calculating the percent of respondents who selected ‘Important’ or ‘Very Important’ for each outcome (i.e. Percent Support). Ratings from pilot and national validation surveys were combined as “Validation” (RQ1b). Each circle represents a single learning outcome. Horizontal lines indicate means across all outcomes from that survey. Points are jittered to reveal distribution. These data are represented in tabular form in Table 3.

We investigated six categorical independent variables as fixed effects: (1) the competency associated with the learning outcome (see six core competencies in Table 1) and five respondent demographics. The demographic variables were: (2) institution type (Associate’s, Bachelor’s, Master’s, or Doctoral Granting) and whether or not the respondent (3) has experience in discipline-based education research (DBER), (4) is currently engaged in disciplinary biology research, (5) has experience in ecology/evolutionary biology research, or (6) has familiarity with Vision and Change. These respondent characteristics were coded using answers to the survey’s demographic questions (e.g., DBER experience and ecology/evolution experience variables were inferred from jointly considering responses to field of current research and graduate training questions).

**Table 3.**
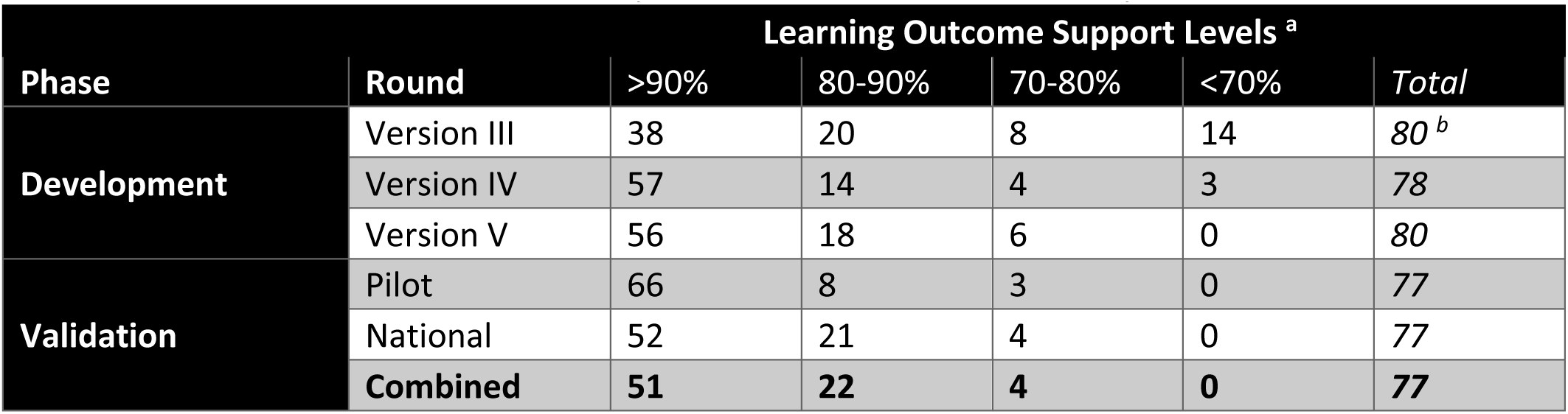
Learning outcome ratings show increasing support over iterative rounds of revision. ^a^ Survey ratings were summarized by calculating the percent of respondents who selected ‘Important’ or ‘Very Important’ for each outcome (i.e. Percent Support). Outcomes were then binned into the indicated ranges. These data are visually represented in Figure 2. ^b^ One outcome (out of 81 total) was mistakenly omitted from the Version III survey.

We used backward model selection to test our hypotheses that the competency of learning outcomes (RQ2a) and the demographics of respondents (RQ2b) affect respondents’ rating of learning outcomes. For each research question, we began with a complex model and removed fixed effects one-by-one that did not improve model fit in order to find the best fitting and most parsimonious models. Specifically, for RQ2a, the initial complex model used ‘Support’ as the dependent variable and included a random effect for learning outcome, a random effect for respondent, and a fixed effect for learning outcome competency. For RQ2b, the initial complex model used ‘Support’ as the dependent variable and included a random effect for learning outcome, a random effect for respondent, and five interactions as fixed effects: competency X institution type, competency X experience in DBER, competency X engagement in disciplinary biology research, competency X experience in ecology/evolution, and competency X Vision and Change familiarity.

During model selection, we determined model fit by comparing the Akaike’s Information Criterion (AIC) value of each model to the previous model. We interpreted two models with ΔAIC ≤2 to have equivalent fit, and in those cases chose the more parsimonious model. Otherwise, the model with the lower AIC value was interpreted to have a better fit. We used likelihood ratio tests to investigate the fit of random effects. Inclusion of random effects for learning outcome and respondent was supported for all models.

As there are numerous problems with interpreting individual coefficients from logistic regression models (Long & Freese, 2014; Mustillo, Lizardo, & McVeigh, 2018), we used predicted probabilities to interpret the best fitting models. For RQ2a, we used the estimated regression equation from the best fitting model to calculate the predicted probability that a respondent would support an outcome within each of the six competencies. For RQ2b, we used the estimated regression equation from the best fitting model to calculate the predicted probability of outcome support for each combination of competency and respondent demographics of interest, holding all other variables at their means (Long & Freese, 2014). When comparing two predicted probabilities, we considered non-overlapping 95% confidence intervals as statistically significant differences.

Additional details on data processing, analysis of missing data, and descriptive statistics of our six independent variables can be found in Supplemental Methods and Supplemental Tables 10-11.

### Aligning Examples with Learning Outcomes

During initial drafting, several faculty included a list of examples of in-class activities and assignments associated with each learning outcome. After national validation, we updated this list by revising, adding, or re-aligning examples in keeping with outcome revisions. Example additions drew from conversations with biology educators throughout the development phase. Two authors (AWC and AJC), who have experience teaching undergraduate biology courses and expertise in molecular and cell biology, carried out the drafting and revising portion of this work. To confirm alignment of the examples with corresponding course-level learning outcomes, three additional college biology instructors (including author JCH) independently reviewed the examples and assessed alignment (yes/no). We selected these additional example reviewers based on their complementary expertise in ecology, evolutionary biology, and physiology. We removed or revised examples until unanimous agreement on alignment was reached.

## RESULTS

### Development of the BioSkills Guide

#### RQ1a: Can we identify an essential set of learning outcomes aligned with the Vision and Change core competencies?

Soliciting and incorporating feedback from participants with diverse professional expertise in undergraduate biology education was essential to ensure we identified core competency learning outcomes that were useful on a broad scale. The initial draft of the BioSkills Guide was crafted by faculty and expanded to include input from 51 unique participants from at least 8 institutions. We then carried out five increasingly larger rounds of review and revision, engaging approximately 218 unique participants from at least 87 institutions (Table 2). Throughout the development phase, we monitored demographics of participant pools and took steps to gather feedback from traditionally under-sampled groups (Figure 3C, Supplemental Tables 2-3).

**Figure 3.**
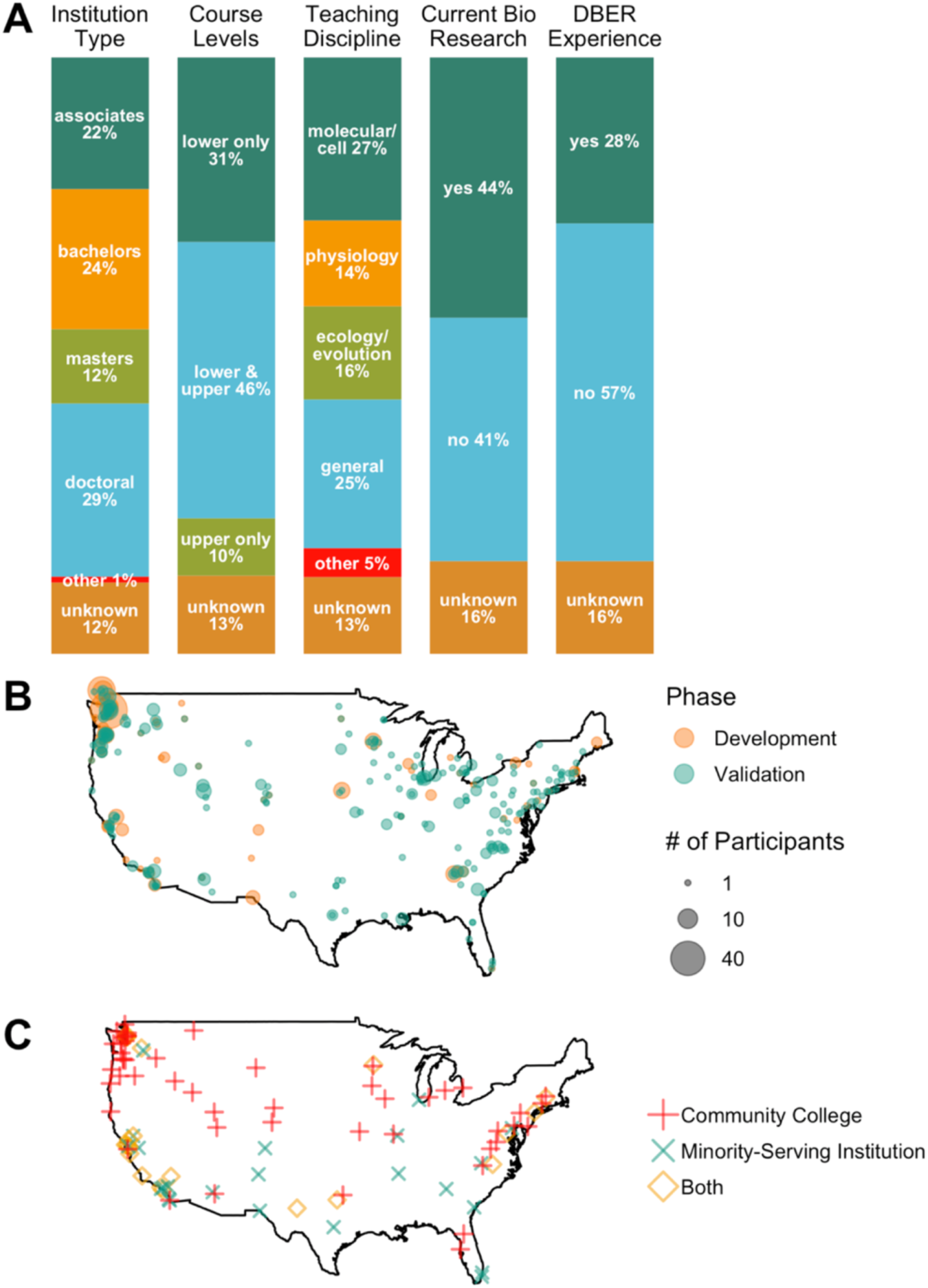
BioSkills Guide development and validation participants spanned a range of institution types, expertise, and geographic locations. **(A) Self-reported demographics of validation phase survey respondents** (n=417). Current engagement in disciplinary biology research was inferred from field of current research. Experience in Discipline-Based Education Research (DBER) was inferred from fields of current research and graduate training. **(B) Geographic distribution of participants**. 263 unique institutions, representing 556 participants with known institutions. Size is proportional to the number of participants from that institution. Only institutions in the continental US and British Columbia are shown. Additional participants came from Alaska, Alberta, Hawaii, India, Puerto Rico, and Scotland (8 institutions). **(C) Geographic distribution of participants from community colleges and minority-serving institutions (MSIs).** 73 unique community colleges and 49 unique MSIs (46 shown; not shown are MSIs in Alaska and Puerto Rico). 23 institutions were classified as both community colleges and MSIs.

To triangulate faculty perceptions of competency outcomes, we collected and applied quantitative and qualitative feedback on drafts of the BioSkills Guide (Figure 1). In general, we observed that interview, workshop, and round table data corroborated many of the trends observed from the surveys, with the same outcomes being least supported (e.g., rated ‘Unimportant’) or arousing confusion (e.g., rated ‘Difficult’ to understand). This provided evidence that the survey was as effective as the other qualitative methods at gauging faculty perceptions of competencies. The survey therefore enabled us to quantitatively assess areas of strength and weakness within drafts more quickly and across a broader population. Using both quantitative and qualitative feedback, every outcome was revised for substance and/or style at least once over the course of the development phase, with most outcomes being revised several times (Supplemental Table 1).

There are four key structural features of the BioSkills Guide that were introduced by faculty early in the development phase. First, the initial draft was written as *learning outcomes* (i.e. descriptions of what students will be able to know and do) rather than statements (i.e. descriptions of the competency itself). We kept this structure to better support backward design (Wiggins & McTighe, 1998). Second, the guide has a two-tiered structure: each core competency contains 2-6 *program-level* learning outcomes, and each program-level learning outcome contains 2-6 *course-level* learning outcomes (illustrated in Supplemental Figure 1). Faculty who participated in the initial drafting spontaneously generated this nested organization, likely reflecting their intended use(s) of the guide for a range of curricular tasks at the program and course levels. Third, the initial draft was written at the level of *a graduating general biology major* (four-year program). We decided to keep this focus to align with the goals of Vision and Change which presented the core concepts and competencies as an overarching framework for the entire undergraduate biology curricula (AAAS, 2011). A similar approach was taken during development of the BioCore Guide for the core concepts, based on their alignment with Vision and Change and their finding that the vast majority of colleges offer a general biology degree (Brownell, Freeman, et al., 2014). Finally, we decided, via conversations with our advisory board, to include only *measurable* learning outcomes so as to directly support assessment use and development. This led us to reframe outcomes related to student attitudes and affect (e.g., an outcome on appreciating the role of science in everyday life was revised to “use examples to describe the relevance of science in everyday experiences”).

### National Validation of the BioSkills Guide

#### RQ1b: How much do biology educators agree on this essential set of competency learning outcomes?

We gathered evidence of content validity of the final draft of the BioSkills Guide using a national survey. We decided to move to validation based on the results of the fifth round of review (Version V). Specifically, the lowest rated outcome from the Version V survey had 72.7% support (Figure 2, Supplemental Table 4). The previous minimums were 16.7% and 50% for Versions III and IV surveys, respectively. Furthermore, all outcomes were rated ‘Easy’ or ‘Very Easy’ to understand by the majority of respondents (Supplemental Figure 2, Supplemental Table 5), and no new substantial suggestions for changes were raised in survey comments or workshop feedback on Version V.

The validation survey included 417 college biology educators, from at least 225 institutions, who evaluated the learning outcomes for their importance for a graduating general biology major (Table 2). Respondents had representation from a range of geographic regions, biology subdisciplines taught, course levels taught, research focuses, and institution types (Figure 3, Supplemental Table 6), including respondents representing a range of community colleges and minority-serving institutions (Figure 3C, Supplemental Table 3).

Each respondent was asked to review a subset of outcomes, resulting in each outcome being reviewed by 211-237 college biology educators. The lowest mean importance rating for any outcome was 4 (equivalent to a rating of ‘Important’), and the average mean importance rating across all outcomes was 4.5 (Supplemental Tables 4, 7). We additionally inferred ‘Percent support’ for each outcome by calculating the percent of respondents who reviewed it who rated it as ‘Important’ or ‘Very Important’. Percent support ranged from 74.3-99.6%, with a mean of 91.9% (Figure 2, Supplemental Table 4). Nearly two-thirds (or 51) of the 77 outcomes had greater than 90% support (Table 3). Four outcomes had less than 80% support, with the lowest rated outcome being supported by 74% of respondents who reviewed it (Table 4). In addition to having them rate the outcomes, we asked respondents to describe any essential learning outcomes that were missing from the guide (summarized in Supplemental Table 8).

**Table 4.** Top five and bottom five supported learning outcomes from validation phase. ^a^ All outcomes shown except “Modeling: Build and evaluate models of biological systems” are course-level learning outcomes. ^b^ Percent support was calculated as the percent of respondents who rated the outcome as ‘Important’ or ‘Very important’. Five highest and lowest rated outcomes by percent support are shown. ^c^ Mean, maximum, and minimum of survey respondents’ importance ratings, where 5 = ‘Very Important’ and 1 = ‘Very Unimportant’.

| Competency | Outcome <sup>a</sup> | Percent Support <sup>b</sup> | Mean <sup>c</sup> | Max <sup>c</sup> | Min <sup>c</sup> |
| --- | --- | --- | --- | --- | --- |
| Quantitative Reasoning | Perform basic calculations (e.g., percentages, frequencies, rates, means). | 99.6 | 4.9 | 5 | 3 |
| Quantitative Reasoning | Create and interpret informative graphs and other data visualizations. | 99.6 | 4.9 | 5 | 3 |
| Process of Science | Analyze data, summarize resulting patterns, and draw appropriate conclusions. | 99.1 | 4.8 | 5 | 1 |
| Quantitative Reasoning | Interpret the biological meaning of quantitative results. | 99.1 | 4.7 | 5 | 3 |
| Quantitative Reasoning | Record, organize, and annotate simple data sets. | 98.7 | 4.8 | 5 | 3 |
| Process of Science | Evaluate and suggest best practices for responsible research conduct (e.g., lab safety, record keeping, proper citation of sources). | 82 | 4.2 | 5 | 2 |
| Science & Society | Identify and describe how systemic factors (e.g., socioeconomic, political) affect how and by whom science is conducted. | 78.9 | 4.1 | 5 | 1 |
| Modeling | Modeling: Build and evaluate models of biological systems. <sup>a</sup> | 75.5 | 4 | 5 | 1 |
| Interdisc. Nature of Science | Suggest how collaborators in STEM and non-STEM disciplines could contribute to solutions of real-world problems. | 74.3 | 4 | 5 | 1 |
| Interdisc. Nature of Science | Describe examples of real-world problems that are too complex to be solved by applying biological approaches alone. | 74 | 4 | 5 | 1 |

### Interpreting Statistical Models of Learning Outcome Support

#### RQ2a: Does biology educators’ support of learning outcomes differ across competencies?

For RQ2a, we hypothesized that differences in learning outcome ratings (as observed in RQ1b) could, in part, be explained by the learning outcome’s competency, with certain competencies being more supported than others. Indeed, a model that included competency had a better fit than one that did not (ΔAIC = -22.21, Supplemental Table 12). It is worth noting that despite the fact that inclusion of competency improved model fit, predicted probabilities of support were high across all six competencies (ranging from 94.2% to 99.1% support, Figure 4A).

**Figure 4.**
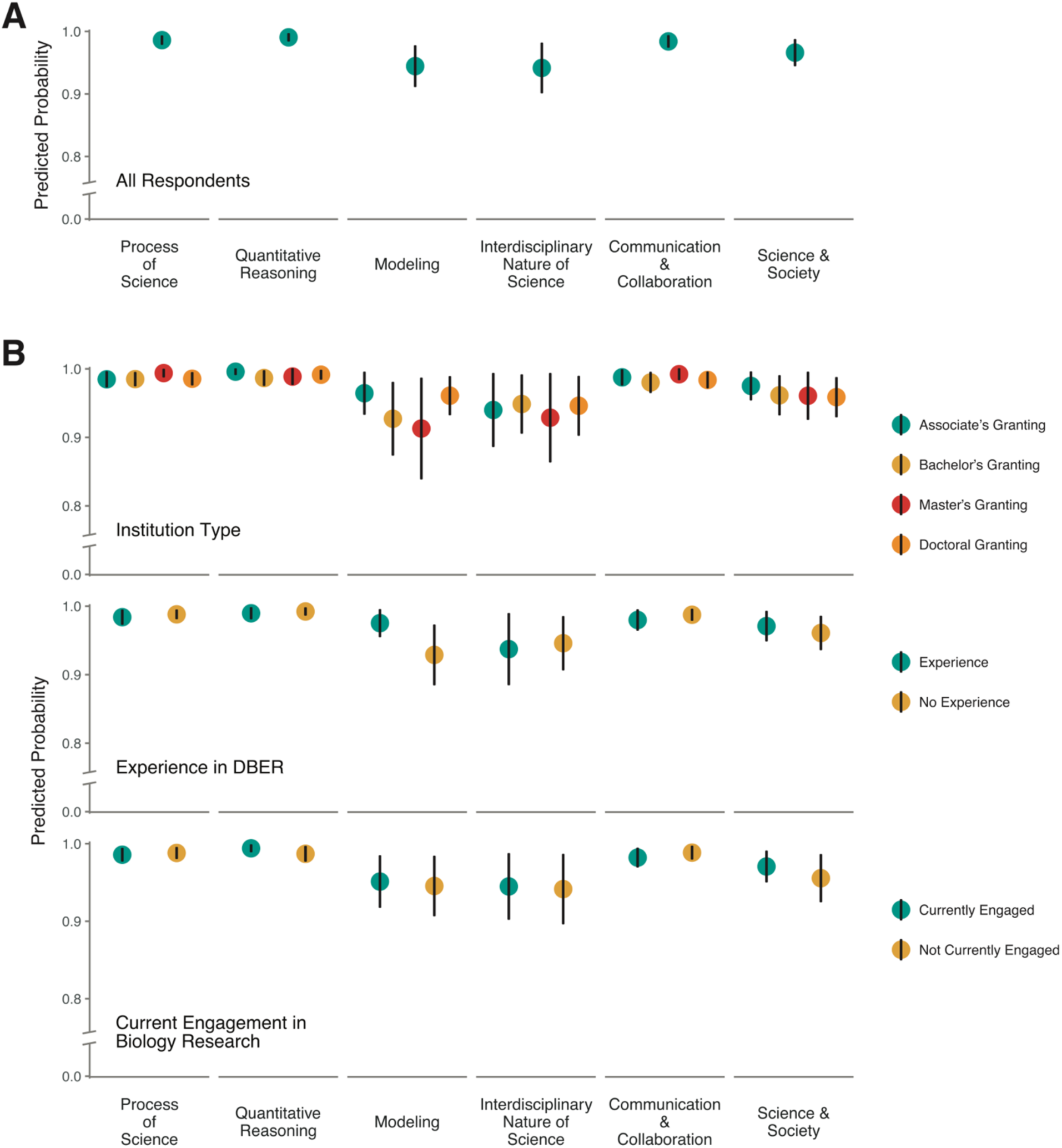
Competency and respondent demographics have significant but small effects on learning outcome support. Predicted probabilities of a respondent supporting (i.e. rating ‘Important’ or ‘Very Important’) a learning outcome in the indicated competency for **(A)** all respondents (RQ2a) or **(B)** respondents in various demographic groups (RQ2b). Predicted probabilities were calculated using best fitting models for each research question. Vertical lines represent 95% confidence intervals. Note that y-axis has been truncated.

#### RQ2b: Do biology educators with different professional backgrounds differ in their support of learning outcomes across competencies?

For RQ2b, we hypothesized that differences in respondent demographics like expertise (i.e. experience in DBER, experience with ecology/evolutionary biology research, familiarity with Vision and Change) or professional culture (i.e. institution type, current engagement in disciplinary biology research) would affect respondents’ support of learning outcomes in different competencies, likely through differences in perceptions of their usefulness or feasibility. For example, respondents who have spent time conducting ecology and/or evolutionary biology research might rate Modeling and Quantitative Reasoning learning outcomes more highly because of the important role quantitative modeling has historically played in these fields. We tested this hypothesis using backward model selection, fitting models including the interaction of competency and our five respondent demographics. We found that the best fitting model was one that included three competency by demographic interactions and one respondent demographic main effect. Specifically, respondents’ support of outcomes within each competency differed based on their institution type, experience in DBER, and current engagement in biology research (Supplemental Table 12). Respondents’ support of outcomes within each competency did not differ based on their familiarity with Vision and Change nor their experience with ecology/evolutionary biology research, however experience with ecology/evolutionary research was retained in the best fitting model as a main effect (Supplemental Figure 3).

The magnitudes of the observed differences were again small (Figure 4B). For example, respondents who have experience with DBER exhibited similarly high support for Modeling (97.5%), Quantitative Reasoning (99.0%), Process of Science (98.4%), and Communication and Collaboration (98.0%) outcomes. In contrast, respondents who do not have experience with DBER were statistically significantly less likely to support Modeling outcomes (92.9%) than Quantitative Reasoning (99.2%), Process of Science (98.8%), or Communication and Collaboration (98.8%) outcomes (i.e. the confidence intervals did not overlap; Figure 4B). However, predicted probabilities for learning outcome support were uniformly above 90% for all respondent groups and competencies, and the greatest difference observed was 6.3%.

### Summary of the Core Competencies

Below we provide descriptions of the core competencies that summarize our understandings of college biology educator priorities, as represented by the learning outcomes in the final draft of the BioSkills Guide (Supplemental Materials).

#### Process of Science

The Process of Science outcomes are presented in a particular order; however, in practice, they are applied in a non-linear fashion. For example, scientific thinking and information literacy include foundational scientific competencies such as critical thinking and understanding the nature of science, and thus are integral to all parts of the process of science. Question formulation, study design, and data interpretation and evaluation are iteratively applied when carrying out a scientific study, and also must be mastered to achieve competence in evaluating scientific information. The final program-level outcome, “Doing Research”, emerged from conversations with biology educators who emphasized that the experience of applying and integrating the other Process of Science outcomes while engaging in authentic research leads to outcomes that are likely greater than the sum of their parts. Course-based or independent research experiences in the lab or field are generally thought to be particularly well-suited for teaching Process of Science; however, many of these outcomes can also be practiced by engaging with scientific literature and existing datasets. Competence in Process of Science outcomes will help students become not only proficient scientists, but also critical thinkers and scientifically literate citizens.

#### Quantitative Reasoning

This comprehensive interpretation of Quantitative Reasoning includes math, logic, data management and presentation, and an introduction to computation. Beyond being essential for many data analysis tasks, this competency is integral to work in all biological subdisciplines and an important component of several other core competencies. Indeed, the universality of math and logic provide a “common language” that can facilitate interdisciplinary conversations. Furthermore, the outcomes emphasize the application of quantitative reasoning *in the context of* understanding and studying biology, mirroring national recommendations to rethink how math is integrated into undergraduate biology coursework. In summary, the outcomes presented here can be included in nearly any biology course to support the development of strong quantitative competency.

#### Modeling

Models are tools that scientists use to develop new insights into complex and dynamic biological structures, mechanisms, and systems. Biologists routinely use models informally to develop their ideas and communicate them with others. Models can also be built and manipulated to refine hypotheses, predict future outcomes, and investigate relationships among parts of a system. It is important to note that there are many different types of models, each with their own applications, strengths, and limitations which must be evaluated by the user. The Modeling outcomes can be practiced using an array of different model types: mathematical (e.g., equations, charts), computational (e.g., simulations), visual (e.g., diagrams, concept maps), and physical (e.g., 3D models).

#### Interdisciplinary Nature of Science

Scientific phenomena are not constrained by traditional disciplinary silos. To have a full understanding of biological systems, students need practice integrating scientific concepts across disciplines, including multiple fields of biology and disciplines of STEM. Furthermore, today’s most pressing societal problems are ill-defined and multi-faceted and therefore require interdisciplinary solutions. Efforts to solve these complex problems benefit from considering perspectives of those working at multiple biological scales (i.e. molecules to ecosystems), in multiple STEM fields (e.g., math, engineering), in non-STEM fields (e.g., humanities, social sciences), as well as input from those outside of academia (e.g., city planners, medical practitioners, community leaders). Productive interdisciplinary biologists therefore recognize the value in collaborating with experts across disciplines and have the competency needed to communicate with diverse groups.

#### Communication & Collaboration

Communication and collaboration are essential components of the scientific process. These outcomes include competencies for interacting with biologists, other non-biology experts, and the general public for a variety of purposes. In the context of undergraduate biology, metacognition involves the ability to accurately sense and regulate one’s behavior both as an individual and as part of a team. Regardless of their specific career trajectories, all biology students require this competency to thoughtfully and effectively work and communicate with others.

#### Science & Society

Science does not exist in a vacuum. Scientific knowledge is constructed by the people engaged in science. It builds on past findings and changes in light of new interpretations, new data, and changing societal influences. Furthermore, advances in science affect lives and environments worldwide. For these reasons, students should learn to reflexively question not only how scientific findings were made, but by whom and for what purpose. A more integrated view of science as a socially situated way of understanding the world will help students be better scientists, advocates for science, and scientifically literate citizens.

### Examples of Activities that Support Competency Development

The faculty who wrote the initial draft of the BioSkills Guide included classroom examples in addition to learning outcomes. A number of early development phase participants expressed that they appreciated having these examples for use in brainstorming ways competencies might be adapted for different courses. Based on this positive feedback, we decided to retain and supplement the examples so that they could be used by others (Supplemental Materials). These examples are *not* exhaustive and have not undergone the same rigorous process of review as the learning outcomes, but we have confirmed alignment of the examples with five college biology educators with complementary subdisciplinary teaching expertise. We envision the examples aiding with interpretation of the learning outcomes in a variety of class settings (i.e. course levels, subdisciplines of biology, class sizes).

## DISCUSSION

### The BioSkills Guide Is a Nationally Validated Resource for the Core Competencies

Employing feedback from over 600 college biology educators, we have developed and gathered evidence of content validity for a set of 77 essential learning outcomes for the six Vision and Change core competencies. During national validation, all learning outcomes had support from ≥74% of survey respondents, with an average of 92% support. This high support suggests that we successfully recruited and applied input from a range of educators during the development phase. As the broadest competency-focused learning outcome framework for undergraduate biology education to date, the BioSkills Guide provides insight on the array of competencies that biology educators consider essential for all biology majors to master during college. We propose that this guide be used to support a variety of curricular tasks including course design, assessment development, and curriculum mapping (Figure 5).

**Figure 5.**
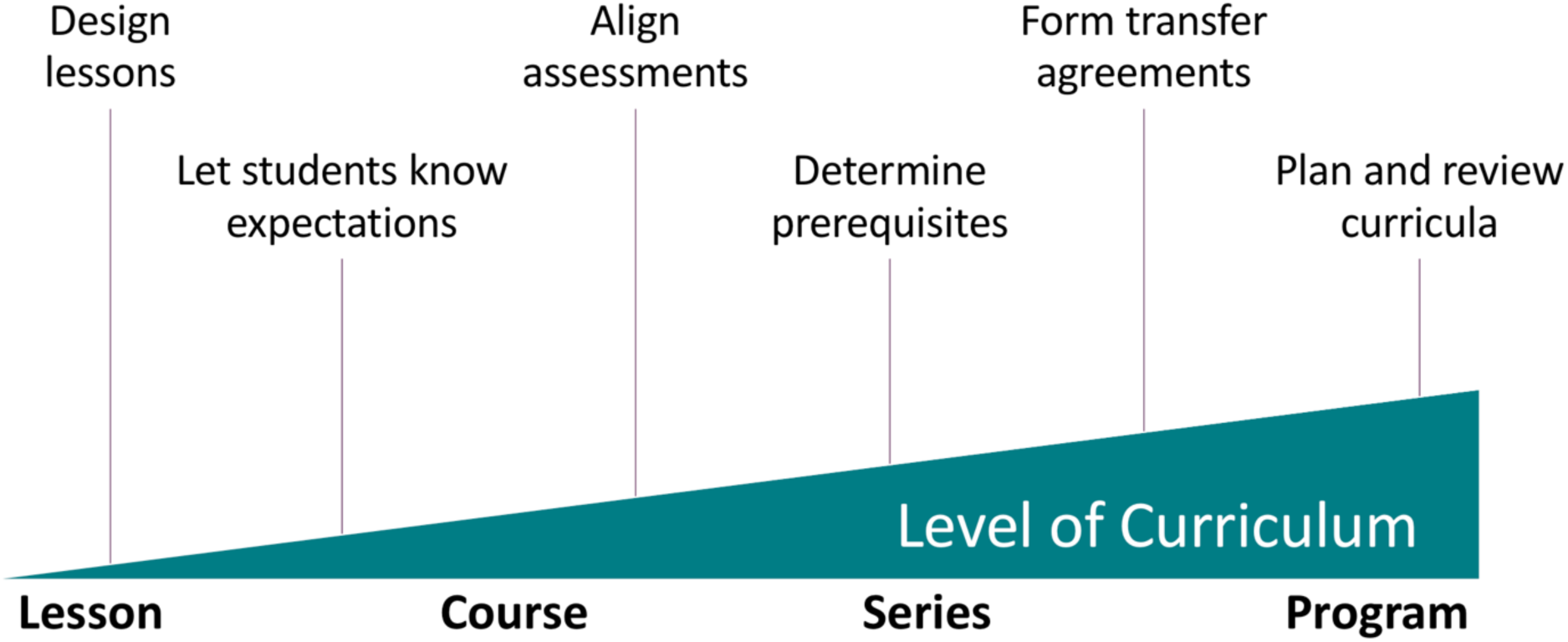
The BioSkills Guide can support a range of curricular scales.

### Examining Variation in Educator Survey Responses

We used statistical modeling to investigate whether respondents’ professional backgrounds could explain their likelihood of supporting outcomes in different competencies. We detected several respondent demographics that were associated with differences in support of learning outcomes within different competencies, however observed differences may not have been large enough to be meaningful on a practical level. In other words, it is unclear whether differences in the perceived importance of particular outcomes by less than 10% of individuals among various educator populations is sufficient to sway curricular decisions.

The results of our RQ2 analyses suggest that (a) there was not sufficient variation in our dataset to detect substantial differences, (b) educators from different backgrounds (at least those investigated in this study) think similarly about competencies, or (c) a combination of these two. In support of (a), 51 out of 77 outcomes had greater than 90% support, likely due to our intentional study design of iteratively revising outcomes to reach consensus during the development phase. In support of (b), it is reasonable that college biology educators in the US are more culturally alike than different given broad similarities in their graduate education experiences (Grunspan, Kline, & Brownell, 2018). Thus, we believe the most likely explanation for the small size of the observed differences is a combination of study design and similarities in educator training.

We could not help but note that where demographic by competency interactions existed, trends, albeit small, consistently pointed toward differences in support for the competency Modeling (Figure 4B). Further work is needed to determine if this trend is supported, but we offer a hypothesis based on observations made over the course of this project: Although we strove to write learning outcomes that are clear and concrete, it is possible that respondents interpreted the difficulty level or focus of modeling-related learning outcomes differently depending on their interpretation of the term “model”. Varying definitions of models were a common theme in survey comments and interviews. Recently a group of mathematicians and biologists (NIMBioS) joined forces to address this issue (Diaz Eaton et al., 2019). They argue that differences in conceptions of modeling among scientists within and across fields have stood in the way of progress in integrating modeling into undergraduate courses. In an effort to improve biology modeling education, they propose a framework, including a definition of model (“a simplified, abstract or concrete representation of relationships and/or processes in the real world, constructed for some purpose”). It is important to note that this definition is not fully consistent with other work on models in science education in its relative emphasis of the role of models for generating new insights vs the role of models as representations (Gouvea & Passmore, 2017). Furthermore, whether a particular representation is considered to be a model depends on how a given user interacts with that representation. For example, an undergraduate student’s drawing illustrating how genes are upregulated by changes in the environment would not bring new insights for a molecular biologist but would be considered a conceptual model for the student, since the student is using the drawing to develop a more sophisticated understanding of how gene expression phenotypes are impacted by environmental conditions (Dauer et al., 2019). While additional work is needed to build a shared understanding of modeling in the undergraduate STEM education community, we believe the NIMBioS definition of model is a valuable starting point for future discussions around the value, relevance, and possible implementations of modeling in college biology. Since the BioSkills Guide elaborates learning outcomes for undergraduate biology majors, we chose a similarly broad definition of models as representations of biological phenomena that can be used for a variety of purposes, as elaborated below in ‘Expanding Modeling’.

### Limitations of the BioSkills Guide

When developing the guide, we made two early design choices that constrained its content. First, we chose to align the outcomes with the Vision and Change core competency framework. We chose this approach in order to build on the momentum Vision and Change has already gained in the undergraduate biology community (AAAS, 2015, 2018, 2019; “About CourseSource,” n.d.; Brancaccio-Taras et al., 2016; Brownell, Freeman, et al., 2014; Cary & Branchaw, 2017; Dirks & Knight, 2016) and thus maximize the chances that we would build a resource that undergraduate biology educators would find useful and adopt. However, due to this choice there are areas in which the guide does not align with other science curriculum frameworks. For example, while Vision and Change core competencies and the Framework for K-12 Science Education scientific practices overlap substantially (Table 1), the latter includes the practice of constructing explanations, where explanations are defined as “accounts that link scientific theory with specific observations and phenomena”. Constructing explanations is not explicitly represented in either Vision and Change or the BioSkills Guide.

The second design choice was that we sought evidence of content validity via a survey of undergraduate biology educators and researchers in biology education, rather than learning sciences researchers who focus on science practices, nature of science, science communication, scientific modeling, etc. We chose this population for our sample because they bring expertise in both biology and biology education and they are the intended users of the guide. To achieve transformation in undergraduate science education, those undergoing the change must be a part of the change process (Henderson, 2010). Furthermore, by developing the guide hand-in-hand with a broad sample of educators, we aimed to create a tool written in the language used and understood by those who would be implementing these practices in their classrooms. In many cases throughout the development phase, we found that small changes in wording affected reviewers’ ratings of the learning outcomes, and thus precise use of language was essential. Indeed, developing a common language around scientific practices (e.g., the distinction between argumentation and explanation) has been shown to be a key step in adoption of NGSS by K-12 teachers (Friedrichsen & Barnett, 2018).

While sampling from this population has advantages, there are also limitations. Although a substantial share of our survey respondents indicated experience in discipline-based education research as well (48.4% during development phase, 27.8% during validation phase), the BioSkills Guide outcomes primarily represent biology educators’ and discipline-based education researchers’ understandings of competencies. Thus, some outcomes represent beliefs held by undergraduate biology educators and researchers that do not fully reflect current understandings in the learning sciences community. One example relates to the definition of “model”, as described above. Another example is the learning outcome “Design controlled experiments, including plans for analyzing the data”, which could be interpreted to overlook the fact that many scientific studies are not experimental (McComas, 1998). In this case, this interpretation would only partially be true. Feedback we received during the development phase indicated that reviewers of the BioSkills Guide in fact recognized the importance of including non-experimental studies when teaching the Process of Science. In response to this feedback, we replaced the word “experiment” in the initial draft with the word “study” in several outcomes to be inclusive of experimental and non-experimental studies. However, workshop and interview data indicated that, on the whole, biology educators also supported explicitly teaching experimental design as a way to introduce students to the rigors of scientific thinking. This led to our retaining the term “experiment” in this particular learning outcome, which received 91.5% support during the validation phase.

Limitations such as these should be kept in mind when interpreting the guide, and we encourage educators to consult multiple frameworks when designing and revising curricula. We suggest that the Framework for K-12 Science Education (NRC, 2012a), and the associated standards (*Next Gener. Sci. Stand. States, By States*, 2013), is an especially important resource for undergraduate biology educators to be familiar with, given its impact in K-12 science education and the importance of scaffolding the transition from secondary to post-secondary science courses. The K-12 Framework has transformed the K-12 education community’s conversations about curriculum by providing a common language with a strong theoretical grounding. Since the framework’s introduction in 2012, understandings of it have naturally deepened through the work of applying it in curriculum and research (Brown & Sadler, 2018). Ongoing implementation work with the scientific practices, especially as they integrate with the framework’s other dimensions (i.e. crosscutting concepts and disciplinary core ideas) has yielded numerous productive insights, including the importance of phenomena as an anchor for 3D curricula (Reiser, Novak, & Mcgill, 2017). In a similar vein, we hope that efforts to implement the BioSkills Guide will help facilitate growth in undergraduate biology education.

Points of discrepancy between the BioSkills Guide and other science education frameworks may reflect areas where understandings of science competencies or practices are still evolving. Future work should consider where and why biology educators’ priorities and conceptions of competencies differ from experts in other fields, including the cognitive and learning sciences and other DBER fields. Such research will undoubtedly be made stronger by working cross-disciplinarily with those experts (Dolan, 2017).

### Defining the Scope of Core Competencies

During the development phase, input from participants led us to expand or revise the focus of certain core competencies relative to their original descriptions in the Vision and Change report (AAAS, 2011). We believe that these evolutions in understanding are in keeping with the spirit of Vision and Change, which encouraged educators to engage in ongoing conversations about elaboration and implementation.

#### Defining the Role of Research in Process of Science

Vision and Change and other leaders in STEM education have emphasized the importance of incorporating research experiences into the undergraduate curriculum (AAAS, 2011; Auchincloss et al., 2014; NASEM, 2017). We therefore drafted a program-level learning outcome related to “doing authentic research” for Process of Science. However, it was initially unclear how this outcome should be worded and what course-level learning outcomes, if any, should be embedded within it. This outcome generally had strong support (>80% rating ‘Important’ or ‘Very Important’) throughout the development phase, but a survey question asking for suggestions of appropriate course-level outcomes yielded only outcomes found elsewhere in the guide (e.g., collaboration, data analysis, information literacy) or affect-related outcomes (e.g., persistence, belonging) which we had previously decided were beyond the scope of this resource. We gained additional insight into this question through qualitative approaches. Roundtable and interview participants reiterated that the learning outcomes associated with research experiences, whether in a course-based or independent setting, were distinct from and “greater than the sum of the parts” of those gained during other activities aimed at practicing individual, related outcomes. Furthermore, numerous participants indicated the outcome was important for supporting continued efforts to systematically include research in undergraduate curricula (also see, Cooper, Soneral, & Brownell, 2017). This feedback prompted us to retain this program-level outcome even though it lacks accompanying course-level learning outcomes.

#### Expanding Modeling

The Vision and Change description of the “Ability to Use Modeling and Simulation” provides examples that emphasize the use of computational and mathematical models, such as “computational modeling of dynamic systems” and “incorporating stochasticity into biological models” (AAAS, 2011). From interviews and survey comments, we found that many participants likewise valued these skill sets, likely because they help prepare students for jobs (also see Durán & Marshall, 2018). However, many participants felt the definition of “modeling” should be expanded to include the use of conceptual models. This sentiment is supported by the K-12 STEM education literature, which establishes conceptual modeling as a foundational scientific practice (NRC, 2012a; Passmore, Stewart, & Cartier, 2009; Svoboda & Passmore, 2013). Such literature defines models and promotes their use based on their ability to help students (and scientists) develop new insights (Gouvea & Passmore, 2017). Indeed, building and interpreting conceptual models supports learning of other competencies and concepts, including data interpretation (Zagallo, Meddleton, & Bolger, 2016), study design (Hester et al., 2018), systems thinking (Bergan-Roller, Galt, Chizinski, Helikar, & Dauer, 2018; Dauer et al., 2019; Dauer, Momsen, Speth, Makohon-Moore, & Long, 2013), and evolution (Speth et al., 2014). Proponents of incorporating drawing into the undergraduate biology curriculum have made similar arguments to increase the scope of Modeling as a competency (Quillin & Thomas, 2015). Given this expansion of the competency, we decided to revise the competency “title” accordingly. Throughout the project, we found that the phrase “Modeling and Simulation” triggered thoughts of computational and mathematical models and their applications, to the exclusion of conceptual modeling. We have therefore revised the shorthand title of this competency to the simpler “Modeling” to emphasize the range of models (e.g., conceptual, physical, mathematical, computational; also see Diaz Eaton et al., 2019) that students may create and work with in college biology courses.

#### Defining the Interdisciplinary Nature of Science

Like Modeling, the “Ability to Tap into the Interdisciplinary Nature of Science” is a forward-looking competency. It represents the forefront of biological research, but not necessarily current practices in the majority of undergraduate biology classrooms. Elaborating it into learning outcomes therefore required additional work, including interviews with interdisciplinary biologists, examination of the literature (e.g., Gouvea, Sawtelle, Geller, & Turpen, 2013; NAE & NRC, 2014; Project Kaleidoscope, 2011), and discussions at two round tables at national biology education research conferences. Since initiating this work, a framework has been presented for implementing this competency in undergraduate biology education, including a working definition: “Interdisciplinary science is the collaborative process of integrating knowledge/expertise from trained individuals of two or more disciplines—leveraging various perspectives, approaches, and research methods/methodologies—to provide advancement beyond the scope of one discipline’s ability” (Tripp & Shortlidge, 2019). We believe this definition aligns well with the content of the Interdisciplinary Nature of Science learning outcomes in the final draft of the BioSkills Guide, especially in its emphasis on collaboration.

#### Expanding Communication & Collaboration

The faculty team who composed the initial draft of the BioSkills Guide expanded the Communication & Collaboration competency significantly. First, they loosened the constraints implied by the title assigned by Vision and Change (“Ability to Communicate and Collaborate *with Other Disciplines*”) to encompass communication and collaboration with many types of people: other biologists, scientists in other disciplines, and non-scientists. This expansion was unanimously supported by participant feedback throughout the development phase and has been promoted in the literature (Brownell, Price, & Steinman, 2013; Mercer-Mapstone & Kuchel, 2017). Second, the drafting faculty included a program-level outcome relating to metacognition. Metacognition and other self-regulated learning skills were not included in the Vision and Change core competencies, but the majority of survey respondents nonetheless supported these outcomes. Some respondents raised concerns about the appropriateness of categorizing metacognition in this competency. However, since its inclusion was well-supported by qualitative and quantitative feedback and it was most directly connected with this competency, we have retained it here.

### Next Steps for the Core Competencies

The BioSkills Guide defines course- and program-level learning outcomes for the core competencies, but there is more work to be done to support backward design of competency teaching. Instructors will need to create lesson-level learning objectives that describe how competencies will be taught and assessed in the context of day-to-day class sessions. It is likely that a similar national-level effort to define lesson-level objectives would be particularly challenging because of the number of possible combinations. First of all, most authentic scientific tasks (e.g., presenting data for peer review, using models and interdisciplinary understandings to make hypotheses about observed phenomena, proposing solutions to real-world problems) require simultaneous use of multiple competencies. Second, instructors will need to define how core competencies interface with biology content and concepts. To this end, existing tools for interpreting the Vision and Change core concepts (Brownell, Freeman, et al., 2014; Cary & Branchaw, 2017) will be valuable companions to the BioSkills Guide, together providing a holistic view of national recommendations for the undergraduate biology curriculum.

We view the complexities of combining concepts and competencies in daily learning objectives as a feature of the course planning process, allowing instructors to retain flexibility and creative freedom. Furthermore, one well-designed lesson can provide the opportunity to practice multiple concepts and competencies. For example, to model the process of cell respiration, students apply not only the competency of modeling but also conceptual understandings of systems and the transformation of matter and energy (Bergan-Roller et al., 2018; Dauer et al., 2013). The 3D Learning Assessment Protocol (Laverty et al., 2016), informed by the multidimensional design of the Framework for K-12 Science Education (NRC, 2012a), may be a valuable resource for considering these sorts of combinations. Several groups have already begun proposing solutions to this work in the context of Vision and Change (Cary & Branchaw, 2017; Dirks & Knight, 2016).

Another complexity to consider when planning core competency teaching is *at what point in the curriculum* competencies should be taught and *in what order*. Scaffolding competencies across course series or whole programs will require thoughtful reflection on the component parts of each learning outcome and how students develop these outcomes over time. To assist in this work, there are a number of resources focusing on particular competencies (for example, see Angra & Gardner, 2016; Diaz-Martinez et al., 2019; Diaz Eaton et al., 2019; Pelaez et al., 2017; Quillin & Thomas, 2015; Tripp & Shortlidge, 2019; Wilson Sayres et al., 2018), all of which describe specific competencies in further detail than is contained in the BioSkills Guide. Additionally, work in K-12 education, and more recently higher education, developing learning progressions could guide future investigations of competency scaffolding (Schwarz et al., 2009; Scott, Wenderoth, & Doherty, 2019). We encourage educators to be thoughtful not only about how individual competencies can build over the course of a college education, but how all of the competencies will work together to form complex, authentic expertise that is greater than the sum of its parts.

Given that over 50% of STEM majors attend a community college during their undergraduate career (NSF NCSES, 2010), yet less than 5% of biology education research studies include community college participation (Schinske et al., 2017), we were intentional about including community college faculty throughout the development and validation of the BioSkills Guide (Figure 3C, Supplemental Table 3). So, while the learning outcomes are calibrated to what a general biology major should be able to do by the end of a *four-year* degree, we were able to develop widely relevant outcomes by identifying connections between each competency and current teaching practices of two-year faculty. Nonetheless, it remains an open question whether certain competencies should be emphasized at the introductory level, either because they are necessary prerequisites to upper-level work or because introductory biology may be a key opportunity to develop biological literacy for the many people who begin but do not end up completing a life sciences major. Discussions of how and when to teach competencies in introductory biology are ongoing (Kruchten et al., 2018). It will be essential that priorities, needs, and barriers for faculty from a range of institutional contexts, particularly community colleges, are considered in those discussions (for example, Corwin, Kiser, LoRe, Miller, & Aikens, 2019).

### Applications of the BioSkills Guide

#### The BioSkills Guide is intended to be a resource, not a prescription

We encourage educators to adapt the outcomes to align with their students’ interests, needs, and current abilities. Reviewing the suggestions for additional learning outcomes made by national validation survey respondents (Supplemental Table 8) provides some preliminary insight into how educators may choose to revise the guide. For example, some respondents wished to increase the challenge level of particular outcomes (e.g., “use computational tools to analyze large datasets” rather than “describe how biologists answer research questions using… large datasets…”) or to create more focused outcomes (e.g., “describe the ways scientific research has mistreated people from minority groups” rather than “…describe the broader societal impacts of biological research on different stakeholders”). Moreover, the content of the guide as a whole should be revisited and updated over time, as college educator perceptions will evolve in response to the changing nature of biology, the scientific job market, and increased adoption of NGSS at the K-12 level.

We envision many applications of the BioSkills Guide across curricular scales (Figure 5). The guide intentionally contains a two-tiered structure, with program-level learning outcomes that are intended to be completed by the end of a four-year degree and course-level learning outcomes that are smaller in scale and more closely resemble outcomes listed on a course syllabus. The program-level learning outcomes could serve as a framework for curriculum mapping, allowing departments to document which courses teach which competencies and subsequently identify program strengths, redundancies, and gaps. These data can then inform a variety of departmental tasks, including allocating funds for development of new courses, re-evaluating degree requirements, assembling evidence for accreditation, and selecting and implementing programmatic assessments. Course-level learning outcomes can spark more informed discussions about particular program-level outcomes, and will likely be valuable in discussions of articulation and transfer across course levels.

Course-level learning outcomes can additionally be used for backward design of individual courses. It can be immensely clarifying to move from broader learning goals such as “Students will be able to communicate science effectively” to concrete learning outcomes such as “Students will be able to use a variety of modes to communicate science (e.g., oral, written, visual).” Furthermore, the outcomes and their aligned example activities included in the Expanded BioSkills Guide (Supplemental Materials) can be used for planning new lessons and for recognizing competencies that are already included in a particular class. Examples such as “write blogs, essays, papers, or pamphlets to communicate findings”, “present data as infographics”, and “give mini-lectures in the classroom” help emphasize the range of ways communication may occur in the classroom. Once clear learning outcomes have been defined, they can be shared with students to explain the purpose of various activities and assignments and increase transparency in instructor expectations. This may help students develop expert-like values for competency development (Marbach-Ad, Hunt, & Thompson, 2019) and encourage them to align their time and effort with faculty priorities.

The BioSkills learning outcomes may be especially relevant for the design of high-impact practices, such as course-based undergraduate research experiences, service learning, and internships (Auchincloss et al., 2014; Brownell & Kloser, 2015; Kuh, 2008), which already emphasize competencies, but often are not developed using backward design (Cooper et al., 2017). In these cases, there is a risk of misalignment between instructor intentions, in-class activities, and assessments (Wiggins & McTighe, 1998). One possible reason for the lack of backward design in these cases is that writing clear, measurable learning outcomes can be challenging and time-consuming. We hope the BioSkills Guide will allow instructors to more quickly formulate learning outcomes, freeing up time for the subsequent steps of backward design (i.e. designing summative and formative assessments and planning instruction).

Assessment is an essential part of evidence-based curriculum review. For some competencies, such as Process of Science, a number of high-quality assessments have been developed (for example, (Brownell, Wenderoth, et al., 2014; Dasgupta, Anderson, & Pelaez, 2014; Deane, Nomme, Jeffery, Pollock, & Birol, 2016; Gormally et al., 2012; Sirum & Humburg, 2011; Timmerman et al., 2011); for a general discussion of CURE assessment see (Shortlidge & Brownell, 2016)). However, substantial gaps remain in the availability of assessments for most other competencies. The BioSkills Guide could be used as a framework for assessment development, similar to how the BioCore Guide was used to develop a suite of programmatic conceptual assessments intentionally aligned with Vision and Change core concepts (Smith et al., 2019). Given the difficulty of assessing particular competencies (e.g., collaboration) with fixed choice or even written response questions, it is unlikely that a single assessment could be designed to cover all six competencies. However, by aligning currently available competency assessments with the BioSkills Guide, outcomes lacking aligned assessments will become apparent and point to areas in need of future work.

While motivations and paths for implementing the BioSkills Guide will vary by department or instructor, the end goal remains the same: better integration of competency teaching in undergraduate biology education. With more intentional and effective competency teaching, biology graduates will be more fully prepared for their next steps, whether those steps are in biology, STEM more generally, or outside of STEM completely. The six core competencies encompass essential skills, embedded in scientific knowledge, needed in competitive careers and also in the daily life of a scientifically literate citizen. We have developed and gathered content validity evidence for the BioSkills Guide with input from a diverse group of biology educators to ensure value for courses in a variety of subdisciplines and levels, and biology departments at a variety of institution types. Thus, we hope the BioSkills Guide will help facilitate progress in meeting the recommendations of Vision and Change with the long-term goal of preparing students for modern careers.

## ACKNOWLEDGMENTS

This project was funded by the National Science Foundation (DUE 1710772). We thank the UW Department of Biology Undergraduate Program Committee for providing the initial draft of learning outcomes that were used to develop the BioSkills Guide. Thank you to Sara Brownell, Jenny McFarland, Erika Offerdahl, Pamela Pape-Lindstrom, and the UW Biology Education Research Group for their continued feedback and assistance throughout this project. We additionally thank Jess Blum, Jeremy Bradford, Lisa Corwin, Alex Doetsch, Deb Donovan, Jenny Loertscher, Kelly McDonald, Jeff Schinske, and Kimberly Tanner for help recruiting survey participants. We thank Jennifer Doherty and Mary Pat Wenderoth for evaluating the aligned examples, Emily Scott and Sara Brownell for constructive feedback on an early version of this manuscript, and Sarah Eddy and Elli Theobald for consultations on statistical methods. We thank manuscript reviewers for providing valuable input that led to significant changes in the manuscript. Finally, we deeply appreciate the time and expertise of the many biologists and biology educators who provided feedback on the BioSkills Guide.

## Notes

https://qubeshub.org/qubesresources/publications/1305

### Competing Interest Statement

The authors have declared no competing interest.

### Summary of Updates

Additional context has been added comparing the Vision and Change core competencies to the Framework for K-12 Science Education scientific practice. Additional explanation for our choice of approach has been added. Statistical analysis approach was changed to an approach more familiar to our intended readers, but findings and conclusion remain unchanged.

