## Supplemental Materials for "BioSkills Guide: Development and National Validation of a Tool for Interpreting the Vision and Change Core Competencies"

**Supplemental Material 1. BioSkills Guide.**
Nationally validated program- and course-level learning outcomes for the Vision and Change core competencies.

**Supplemental Material 2. Expanded BioSkills Guide with aligned examples.**
Educator-aligned (n=5) examples for each BioSkills Guide course-level learning outcome.

**Supplemental Material 3. Supplemental Figures.**

**Supplemental Material 4. Supplemental Tables.**

**Supplemental Material 5. Supplemental Methods.**
Additional details on the methods used in this project.

**Supplemental Material 6. BioSkills development phase questionnaire.**
Questionnaire used during development phase survey for review of Version V. Questionnaires for versions III and IV were very similar, except for revision of learning outcomes.

**Supplemental Material 7. BioSkills validation phase questionnaire.**
Questionnaire used during national validation survey. Questionnaire for pilot validation was identical, except for wording of one learning outcome.

**Supplemental Material 1. BioSkills Guide.**

Nationally validated program- and course-level learning outcomes for the Vision and Change core competencies.

#### PROCESS OF SCIENCE

| Program-Level Learning Outcomes | Course-Level Learning Outcomes |
|---|---|
| **SCIENTIFIC THINKING** Explain how science generates knowledge of the natural world. | Explain how scientists use inference and evidence-based reasoning to generate knowledge. |
| | Describe the iterative nature of science and how new evidence can lead to the revision of scientific knowledge. |
| **INFORMATION LITERACY** Locate, interpret, and evaluate scientific information. | Find and evaluate the credibility of a variety of sources of scientific information, including popular science media and scientific journals. |
| | Interpret, summarize, and evaluate evidence in primary literature. |
| | Evaluate claims in scientific papers, popular science media, and other sources using evidence-based reasoning. |
| **QUESTION FORMULATION** Pose testable questions and hypotheses to address gaps in knowledge. | Recognize gaps in our current understanding of a biological system or process and identify what specific information is missing. |
| | Develop research questions based on your own or others' observations. |
| | Formulate testable hypotheses and state their predictions. |
| **STUDY DESIGN** Plan, evaluate, and implement scientific investigations. | Compare the strengths and limitations of various study designs. |
| | Design controlled experiments, including plans for analyzing the data. |
| | Execute protocols and accurately record measurements and observations. |
| | Identify methodological problems and suggest how to troubleshoot them. |
| | Evaluate and suggest best practices for responsible research conduct (e.g., lab safety, record keeping, proper citation of sources). |
| **DATA INTERPRETATION & EVALUATION** Interpret, evaluate, and draw conclusions from data in order to make evidence-based arguments about the natural world. | Analyze data, summarize resulting patterns, and draw appropriate conclusions. |
| | Describe sources of error and uncertainty in data. |
| | Make evidence-based arguments using your own and others' findings. |
| | Relate conclusions to original hypothesis, consider alternative hypotheses, and suggest future research directions based on findings. |
| **DOING RESEARCH** Apply science process skills to address a research question in a course-based or independent research experience. | |

#### QUANTITATIVE REASONING

| Program-Level Learning Outcomes | Course-Level Learning Outcomes |
|---|---|
| **NUMERACY** Use basic mathematics (e.g., algebra, probability, unit conversion) in biological contexts. | Perform basic calculations (e.g., percentages, frequencies, rates, means). |
| | Select and apply appropriate equations (e.g., Hardy-Weinberg, Nernst, Gibbs free energy) to solve problems. |
| | Interpret and manipulate mathematical relationships (e.g., scale, ratios, units) to make quantitative comparisons. |
| | Use probability and understanding of biological variability to reason about biological processes and statistical analyses. |
| | Use rough estimates informed by biological knowledge to check quantitative work. |
| | Describe how quantitative reasoning helps biologists understand the natural world. |
| **QUANTITATIVE & COMPUTATIONAL DATA ANALYSIS** Apply the tools of graphing, statistics, and data science to analyze biological data. | Record, organize, and annotate simple data sets. |
| | Create and interpret informative graphs and other data visualizations. |
| | Select, carry out, and interpret statistical analyses. |
| | Describe how biologists answer research questions using databases, large data sets, and data science tools. |
| | Interpret the biological meaning of quantitative results. |

#### MODELING

| Program-Level Learning Outcomes | Course-Level Learning Outcomes |
|---|---|
| **PURPOSE OF MODELS** Recognize the important roles that scientific models, of many different types (conceptual, mathematical, physical, etc.), play in predicting and communicating biological phenomena. | Describe why biologists use simplified representations (models) when solving problems and communicating ideas. |
| | Given two models of the same biological process or system, compare their strengths, limitations, and assumptions. |
| **MODEL APPLICATION** Make inferences and solve problems using models and simulations. | Summarize relationships and trends that can be inferred from a given model or simulation. |
| | Use models and simulations to make predictions and refine hypotheses. |
| **MODELING** Build and evaluate models of biological systems. | Build and revise conceptual models to propose how a biological system or process works. |
| | Identify important components of a system and describe how they influence each other (e.g., positively or negatively). |
| | Evaluate conceptual, mathematical, or computational models by comparing their predictions with empirical data. |

#### INTERDISCIPLINARY NATURE OF SCIENCE

| Program-Level Learning Outcomes | Course-Level Learning Outcomes |
|---|---|
| **CONNECTING SCIENTIFIC KNOWLEDGE** Integrate concepts across other STEM disciplines (e.g., chemistry, physics) and multiple fields of biology (e.g., cell biology, ecology). | Given a biological problem, identify relevant concepts from other STEM disciplines or fields of biology. |
| | Build models or explanations of simple biological processes that include concepts from other STEM disciplines or multiple fields of biology. |
| **INTERDISCIPLINARY PROBLEM SOLVING** Consider interdisciplinary solutions to real-world problems. | Describe examples of real-world problems that are too complex to be solved by applying biological approaches alone. |
| | Suggest how collaborators in STEM & non-STEM disciplines could contribute to solutions of real-world problems. |
| | Be able to explain biological concepts, data, and methods, including their limitations, using language understandable by collaborators in other disciplines. |

#### COMMUNICATION & COLLABORATION

| Program-Level Learning Outcomes | Course-Level Learning Outcomes |
|---|---|
| **COMMUNICATION** Share ideas, data, and findings with others clearly and accurately. | Use appropriate language and style to communicate science effectively to targeted audiences (e.g., general public, biology experts, collaborators in other disciplines). |
| | Use a variety of modes to communicate science (e.g., oral, written, visual). |
| **COLLABORATION** Work productively in teams with people who have diverse backgrounds, skill sets, and perspectives. | Work with teammates to establish and periodically update group plans and expectations (e.g., team goals, project timeline, rules for group interactions, individual and collaborative tasks). |
| | Elicit, listen to, and incorporate ideas from teammates with different perspectives and backgrounds. |
| | Work effectively with teammates to complete projects. |
| **COLLEGIAL REVIEW** Provide and respond to constructive feedback in order to improve individual and team work. | Evaluate feedback from others and revise work or behavior appropriately. |
| | Critique others' work and ideas constructively and respectfully. |
| **METACOGNITION** Reflect on your own learning, performance, and achievements. | Evaluate your own understanding and skill level. |
| | Assess personal progress and contributions to your team and generate a plan to change your behavior as needed. |

| SCIENCE & SOCIETY | |
|---|---|
| **Program-Level Learning Outcomes** | **Course-Level Learning Outcomes** |
| **ETHICS**<br>Demonstrate the ability to critically analyze ethical issues in the conduct of science. | Identify and evaluate ethical considerations (e.g., use of animal or human subjects, conflicts of interest, confirmation bias) in a given research study. |
| | Critique how ethical controversies in biological research have been and can continue to be addressed by the scientific community. |
| **SOCIETAL INFLUENCES**<br>Consider the potential impacts of outside influences (historical, cultural, political, technological) on how science is practiced. | Describe examples of how scientists' backgrounds and biases can influence science and how science is enhanced through diversity. |
| | Identify and describe how systemic factors (e.g., socioeconomic, political) affect how and by whom science is conducted. |
| **SCIENCE'S IMPACT ON SOCIETY**<br>Apply scientific reasoning in daily life and recognize the impacts of science on a local and global scale. | Apply evidence-based reasoning and biological knowledge in daily life (e.g., consuming popular media, deciding how to vote). |
| | Use examples to describe the relevance of science in everyday experiences. |
| | Identify and describe the broader societal impacts of biological research on different stakeholders. |
| | Describe the roles scientists have in facilitating public understanding of science. |


**Alexa Clemmons, Jerry Timbrook, Jon Herron & Alison Crowe**

DEPARTMENT OF BIOLOGY
UNIVERSITY of WASHINGTON



This work was funded by the University of Washington Department of Biology and NSF DUE 1710772 research grant: UW: PI A. Crowe.


**To download or share the BioSkills Guide:**
**https://qubeshub.org/qubesresources/publications/1305**

#### HOW WAS THE BIOSKILLS GUIDE DEVELOPED?

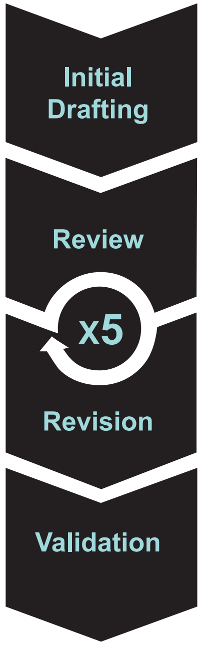


- Began with faculty-crafted draft
- Literature review
- Interviews
- Workshop

- Surveys
- Interviews
- Workshops

n=218 (~60/outcome)

- Survey

n=417 (~210/outcome)

#### HOW CAN THE BIOSKILLS GUIDE HELP YOU?

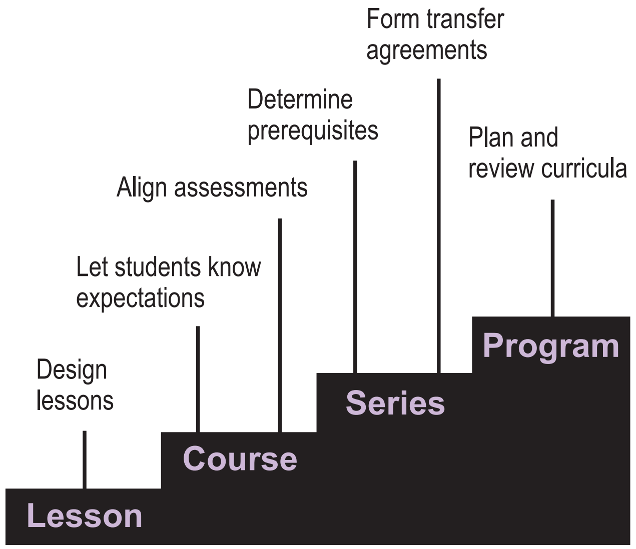


### A Tool for Interpreting the Vision and Change CORE COMPETENCIES

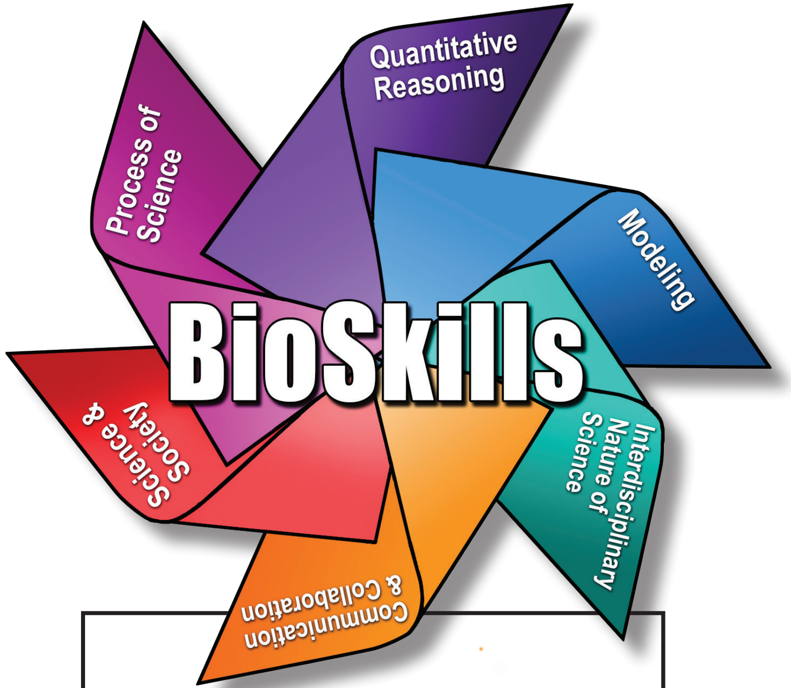


**The BioSkills Guide** comprises program- and course-level learning outcomes that elaborate what general biology majors should be able to do by the time they graduate. Building on the six core competencies of Vision and Change, the learning outcomes were developed and then nationally validated using input from over 600 college biology educators from a range of biology subdisciplines and institution types.

**Supplemental Material 2. Expanded BioSkills Guide with aligned examples.**
Educator-aligned (n=5) examples for each BioSkills Guide course-level learning outcome.

### Expanded BioSkills Guide: Process of Science

| Program-Level Learning Outcomes | Course-Level Learning Outcomes | Examples |
|---|---|---|
| **SCIENTIFIC THINKING Explain how science generates knowledge of the natural world.** | Explain how scientists use inference and evidence-based reasoning to generate knowledge. | Differentiate between evidence and claims in various types of scientific media. |
| | | Explain to a non-scientist what scientists mean when they say "the data suggest…" |
| | | Describe ways that some conclusions could be more certain than others. |
| | | Evaluate popular arguments against vaccines and climate change. |
| | Describe the iterative nature of science and how new evidence can lead to the revision of scientific knowledge. | Provide an example illustrating how a new finding led to the revision of scientific understanding. |
| | | Explain the merits of repeating a study in different contexts (e.g., different model systems, ecological systems, or experimental parameters) for generating increasingly refined hypotheses. |
| **INFORMATION LITERACY Locate, interpret, and evaluate scientific information.** | Find and evaluate the credibility of a variety of sources of scientific information, including popular science media and scientific journals. | Carry out literature searches using databases, Google Scholar, and library resources. |
| | | Identify authors and potential conflicts of interest in web sources. |
| | | Identify reliable online and print sources for use when making decisions. |
| | Interpret, summarize, and evaluate evidence in primary literature. | Summarize conclusions from data figures or Results section of a journal article before reading Discussion section. |
| | | Interpret and teach data figures to peers in jigsaw or small group settings. |
| | | Lead or participate in journal clubs. |
| | Evaluate claims in scientific papers, popular science media, and other sources using evidence-based reasoning. | Differentiate between objective evidence and subjective opinions in popular media or discussion sections of primary literature. |
| | | Critique scientific information in daily life (e.g., nutritional and medical guidelines) by reviewing primary literature. |
| | | Critique the evidence supporting two conflicting hypotheses. |
| | | Compare treatments of similar science topics in primary literature, popular science media, and online discussions. |

### Expanded BioSkills Guide: Process of Science

| Program-Level Learning Outcomes | Course-Level Learning Outcomes | Examples |
|---|---|---|
| **QUESTION FORMULATION Pose testable questions and hypotheses to address gaps in knowledge.** | Recognize gaps in our current understanding of a biological system or process and identify what specific information is missing. | When explaining or diagramming a biological process, recognize areas of uncertainty or missing steps. |
| | | Read the introduction section of a journal article and identify what was known at the time the study began and what unanswered question the study was designed to address. |
| | | Read multiple sources (e.g. primary literature, review articles, popular science articles) and summarize what is known and not known about a particular scientific topic. |
| | Develop research questions based on your own or others' observations. | Review and interpret existing lab notebooks and preliminary data from other lab members in order to identify unanswered questions. |
| | | Suggest follow-up questions based on patterns in data. |
| | | Make note of day-to-day observations that you don't understand and reframe into scientific questions. |
| | Formulate testable hypotheses and state their predictions. | Identify hypotheses and predictions in a scientific publication. |
| | | Evaluate peers' hypotheses for testability. |
| | | Sketch graphs or schematics of predicted study results based on hypotheses. |
| | | Elaborate the proposed model underlying a hypothesis using a cartoon or flow chart. |
| | | Write the "hypothesis & specific aims" portion of a mock grant proposal. |
| **STUDY DESIGN Plan, evaluate, and implement scientific investigations.** | Compare the strengths and limitations of various study designs. | Evaluate peers' study designs' alignment with hypotheses. |
| | | Identify and describe important study design elements of published studies (e.g., predictor and response variables, sample selection, replicates). |
| | | Distinguish between experimental and observational study designs, deductive and inductive approaches. |
| | | Describe the advantages and limitations of different types of studies (e.g., randomized controlled trials, retrospective studies, natural experiments, comparative studies). |
| | Design controlled experiments, including plans for analyzing the data. | Design experiment using simulations. |
| | | Draw a flow diagram or cartoon of proposed or published experimental design. |
| | | Identify and design necessary controls, both biological and methodological. |
| | | Explain how experimental design will account for or detect technical or biological variability. |
| | | Select appropriate measurement and statistical methods for a given research design. |

### Expanded BioSkills Guide: Process of Science

| Program-Level Learning Outcomes | Course-Level Learning Outcomes | Examples |
|---|---|---|
| **STUDY DESIGN** **Plan, evaluate, and implement scientific investigations.** ***(continued)*** | Execute protocols and accurately record measurements and observations. | Read and follow protocols, making note of where practices may have differed from protocol descriptions (i.e. mistakes, ambiguity). |
| | | Keep detailed notes on observations about samples/subjects made before, during, and after protocols. |
| | | Generate organized tables or spreadsheets to record measurements. |
| | | Maintain a laboratory notebook and carefully save and index digital data files. |
| | | Become proficient with common experimental techniques in a given subdiscipline. |
| | Identify methodological problems and suggest how to troubleshoot them. | Interpret positive and negative controls to evaluate success of an experiment. |
| | | Use notes and observations taken during study to pinpoint most likely source of problems. |
| | | Read about methods to propose possible sources of technical errors and appropriate corrections. |
| | Evaluate and suggest best practices for responsible research conduct (e.g., lab safety, record keeping, proper citation of sources). | Complete lab safety training, and maintain recommended practices (e.g., personal protective equipment, proper use and storage of chemicals). |
| | | Use consistent file names and metadata (e.g., collection date, variable naming) formats to save digital files so that they may be used by others in the future. |
| | | Process and store samples using appropriate techniques to preserve data quality (or make note of any improper handling of samples). |

### Expanded BioSkills Guide: Process of Science

<table>
<tr><th>Program-Level Learning Outcomes</th><th>Course-Level Learning Outcomes</th><th>Examples</th></tr>
<tr><td rowspan="12">DATA INTERPRETATION & EVALUATION<br>Interpret, evaluate, and draw conclusions from data in order to make evidence-based arguments about the natural world.</td><td rowspan="4">Analyze data, summarize resulting patterns, and draw appropriate conclusions.</td><td>Transform and display data for exploration and analysis.</td></tr>
<tr><td>Identify trends and distributions in data.</td></tr>
<tr><td>Relate data to conceptual models.</td></tr>
<tr><td>Present data and conclusions clearly, noting limitations.</td></tr>
<tr><td rowspan="3">Describe sources of error and uncertainty in data.</td><td>Select and use appropriate statistical methods to calculate the degree of certainty in results.</td></tr>
<tr><td>Differentiate between sources and effects of technical and biological variability and make appropriate conclusions with such variability in mind.</td></tr>
<tr><td>Describe any mistakes or unexpected conditions during data collection and explain how they could impact conclusions.</td></tr>
<tr><td rowspan="2">Make evidence-based arguments using your own and others' findings.</td><td>Debate the pros and cons of various scientific practices based on evidence (e.g., the use of monocultures by agribusiness).</td></tr>
<tr><td>Write an editorial outlining an argument for or against various scientific applications (e.g., the safety of using CRISPR technology in human embryos).</td></tr>
<tr><td rowspan="3">Relate conclusions to original hypothesis, consider alternative hypotheses, and suggest future research directions based on findings.</td><td>Determine whether hypothesis was supported or refuted by data.</td></tr>
<tr><td>Revise or refine hypothesis or model based on conclusions.</td></tr>
<tr><td>Identify experiments that could be used to resolve ambiguity in results.</td></tr>
<tr><td colspan="2" rowspan="4">DOING RESEARCH<br>Apply science process skills to address a research question in a course-based or independent research experience.</td><td>Identify a novel research question and propose an appropriate study design to test it.</td></tr>
<tr><td>Given a research question, formulate a hypothesis, identify a relevant online data set, and run appropriate analyses to test hypothesis.</td></tr>
<tr><td>Follow protocols to gather data in the field or lab, summarize and find patterns, and identify follow-up questions to address uncertainty in results.</td></tr>
<tr><td>After attempting an experiment or study, reflect on its success and failures and repeat with adjustments.</td></tr>
</table>

### Expanded BioSkills Guide: Quantitative Reasoning

| Program-Level Learning Outcomes | Course-Level Learning Outcomes | Examples |
|---|---|---|
| **NUMERACY** **Use basic mathematics (e.g., algebra, probability, unit conversions) in biological contexts.** | Perform basic calculations (e.g., percentages, frequencies, rates, means). | Perform calculations as part of experimental planning (e.g., plan serial dilutions, use dimensional analysis to convert units). |
| | | Use data to calculate rates of change (e.g., mutation rate, growth rate). |
| | | Summarize data sets using common descriptive statistics (e.g., mean, median, standard deviation). |
| | Select and apply appropriate equations (e.g., Hardy-Weinberg, Nernst, Gibbs free energy) to solve problems. | Use Punnett Squares and related equations to calculate predicted phenotype and genotype frequencies from crosses. |
| | | Plan how to prepare solutions and reaction mixes using $C_1*V_1=C_2*V_2$. |
| | | Identify appropriate equations to interpret a given data set or scenario (e.g., population growth equations, Nernst and Goldman equations for equilibrium and membrane potentials). |
| | | Translate words and concepts (e.g., descriptions of systems, hypotheses) into equations and terms (e.g., coefficients, rate of change). |
| | Interpret and manipulate mathematical relationships (e.g., scale, ratios, units) to make quantitative comparisons. | Convert between related units (e.g., given concentration, convert volume to mass). |
| | | Calculate and interpret fold-change in a variable over time. |
| | | Interpret units reported on graphs (e.g., differentiate between linear and log scales, interpret slope based on units of X and Y-axes). |
| | | Relate surface area to volume in various biological structures (e.g., plasma membranes, alveoli, leaves). |
| | Use probability and understanding of biological variability to reason about biological processes and statistical analyses. | Apply ‘either-or’ and ‘both-and’ rules to calculate combined probabilities. |
| | | Explain the strengths and limitations of using a p-value criterion (e.g., $<0.05$) to determine significance. |
| | | Explain the difference between different measures of error and variation (e.g., standard error the mean vs. standard deviation). |

#### Expanded BioSkills Guide: Quantitative Reasoning

| Program-Level Learning Outcomes | Course-Level Learning Outcomes | Examples |
|---|---|---|
| **NUMERACY** **Use basic mathematics (e.g., algebra, probability, unit conversions) in biological contexts.** ***(continued)*** | Use rough estimates informed by biological knowledge to check quantitative work. | Make order-of-magnitude estimates. |
| | | Extrapolate from data in order to make predictions. |
| | Describe how quantitative reasoning helps biologists understand the natural world. | Identify quantitative approaches that could be used when solving biological problems. |
| | | Discuss examples of how 'big data' has allowed biologists to answer new research questions (e.g., the identification of quantitative trait loci, the use of satellite data to inform ecological models). |
| **QUANTITATIVE & COMPUTATIONAL DATA ANALYSIS** **Apply the tools of graphing, statistics, and data science to analyze biological data.** | Record, organize, and annotate simple data sets. | Save and organize data with future users in mind (e.g., include units, clearly label columns and rows, use intuitive file names). |
| | | Identify and include relevant metadata in data tables (e.g., collection dates, origin of data). |
| | | Add appropriately annotated data to large public databases and discuss the importance of data annotation in the maintenance of these resources. |
| | | Use Excel, R, Python, Mathematica, or other programs to perform basic tasks in data management. |
| | Create and interpret informative graphs and other data visualizations. | Interpret tables and data visualizations (e.g., scatter plots, bar graphs, boxplots, histograms) in primary literature. |
| | | Choose and create best form of chart for data type (e.g., logarithmic scale for growth rates, bar graphs for categorical data, histograms for counts). |
| | | Modify visualization to emphasize important relationships between variables (e.g., add trend lines, color code subsets of data). |
| | | Make predictions and construct explanations based on your own or others' data visualizations. |
| | Select, carry out, and interpret statistical analyses. | Calculate and explain the uses of different types of descriptive statistics (e.g., mean vs median, standard deviation vs. standard error of the mean). |
| | | Use software (e.g., Excel, R) to calculate inferential statistics. |
| | | Interpret statistics in primary literature (e.g., error bars and *p*-values). |
| | | Select appropriate inferential statistical methods for research question (e.g., t-test for comparing mean of two groups, linear regression for modeling relationship between multiple variables, Chi-square for comparing distributions). |

#### Expanded BioSkills Guide: Quantitative Reasoning

| Program-Level Learning Outcomes | Course-Level Learning Outcomes | Examples |
|---|---|---|
| **QUANTITATIVE & COMPUTATIONAL DATA ANALYSIS** **Apply the tools of graphing, statistics, and data science to analyze biological data.** ***(continued)*** | Describe how biologists answer research questions using databases, large data sets, and data science tools. | Give examples of research tasks that can be aided by common bioinformatic tools (e.g., BLAST to find homologs, Clustal to identify differences between sequences, Primer3 to reduce likelihood of unintended PCR products). |
| | | Browse and describe the types of data available in various public databases (e.g., GenBank, UCSC genome browser, 1001 Genomes, NEON). |
| | | Discuss examples where data science has contributed to our understanding of biology (e.g., genomics and genetics, metabolomics and human health, satellite data and the impacts of climate change). |
| | Interpret the biological meaning of quantitative results. | Describe equations and coefficients in terms of their biological meaning (e.g., k as “carrying capacity”, $N_e$ as “effective population size”, $C_t$ values as “gene expression levels”). |
| | | Interpret what graph curves (e.g., linear, exponential, saturation, sigmoidal) mean in different biological contexts (e.g., population growth, enzyme kinetics). |
| | | Summarize data and relate back to hypotheses and other knowledge. |
| | | Write the discussion section of a lab report, including alternative interpretations of data. |

### Expanded BioSkills Guide: Modeling

| Program-Level Learning Outcomes | Course-Level Learning Outcomes | Examples |
|---|---|---|
| **PURPOSE OF MODELS Recognize the important roles that scientific models, of many different types (conceptual, mathematical, physical, etc.), play in predicting and communicating biological phenomena.** | Describe why biologists use simplified representations (models) when solving problems and communicating ideas. | Provide examples of models used by biologists (e.g., animal models of human disease, mathematical models of population genetics, 3D models of anatomical structures) and explain their advantages and disadvantages over "the real thing". |
| | | Describe ways you use models in your own studying and research (e.g., textbook schematics or simulations for learning about abstract ideas, conceptual models when formulating hypotheses, 3D models in lab). |
| | | Identify aspects of biological systems that would likely be simplified in a model and explain why (e.g., system dynamics are often omitted from 2D models because of difficulty in representing time). |
| | Given two models of the same biological process or system, compare their strengths, limitations, and assumptions. | Describe the purposes of different types of models (e.g., physical models for understanding 3D structure, mathematical models for predicting future events). |
| | | Discuss the tradeoffs between model accuracy and simplicity. |
| | | Identify and describe assumptions made in a given model or simulation (i.e. simplified conditions and unknown relationships). |
| | | Choose and justify which model would be better for a given research question based on which parameters are included in the model and which are simplified. |
| **MODEL APPLICATION Make inferences and solve problems using models and simulations.** | Summarize relationships and trends that can be inferred from a given model or simulation. | Determine how two variables relate by manipulating a model and interpreting its output (e.g., the influence of Keq on ΔG in enzymatic reaction coupling, the interplay of resistance and concentration gradient in flux). |
| | | Sketch a flow chart or cartoon of a biological process based on the output of a model or simulation. |
| | | Infer biological trends based on the shape of the model output curve (e.g., linear, exponential, saturation, sigmoidal). |
| | Use models and simulations to make predictions and refine hypotheses. | Predict the impact of changing parameters on various outputs (e.g., the effect of selection coefficients on allele frequencies, the effect of environmental resources on fitness over time). |
| | | Identify key components of a system based on their relative importance in a model's ability to explain data (e.g., master regulator transcription factors, keystone species). |
| | | Propose environmental or public health policy solutions based on models and simulations (e.g., priorities for habitat restoration). |

### Expanded BioSkills Guide: Modeling

| Program-Level Learning Outcomes | Course-Level Learning Outcomes | Examples |
|---|---|---|
| **MODELING**<br>**Build and evaluate models of biological systems.** | Build and revise conceptual models to propose how a biological system or process works. | Sketch flow charts, diagrams, or concept maps while problem solving to organize thinking. |
| | | Draw a cartoon or diagram of a biological process or system consistent with a given set of data. |
| | | Generate a concept map using index cards or online programs to identify and visualize connections between concepts (e.g., transcription, translation, and signaling; ventilation, heart rate, and $O_2$ levels). |
| | | Create diagrams or 3D models that emphasize important aspects of biological structures (e.g., ability to separate DNA base pairs for replication, amino acid R-group structure for protein function, tissue organization for organ function). |
| | Identify important components of a system and describe how they influence each other (e.g., positively or negatively). | Given a biological system (e.g., gut microbiome, carbon cycle, cellular respiration), list relevant components and categorize them as inputs, outputs, or mediators. |
| | | Simplify models by identifying and removing components that are not necessary to recreate patterns of interest. |
| | | Add quantitative signifiers to a concept map (i.e. "-" indicates two components covary indirectly). |
| | Evaluate conceptual, mathematical, or computational models by comparing their predictions with empirical data. | Conduct quality control tests by defining expected outcomes for a model or simulation, including conditions under which expected behaviors should occur (e.g., lac operon expression in presence of lactose and/or glucose). |
| | | Use statistics (e.g., Chi-square tests, t-tests) to compare model outputs to observed distributions. |
| | | When model predictions and empirical data don't match, propose variables that may be missing from model that could explain difference. |
| | | Iteratively modify a model or simulation until quality control is passed. |

### Expanded BioSkills Guide: Interdisciplinary Nature of Science

| Program-Level Learning Outcomes | Course-Level Learning Outcomes | Examples |
|---|---|---|
| **CONNECTING SCIENTIFIC KNOWLEDGE Integrate concepts across other STEM disciplines (e.g., chemistry, physics) and multiple fields of biology (e.g., cell biology, ecology).** | Given a biological problem, identify relevant concepts from other STEM disciplines or fields of biology. | Identify and use relevant knowledge from chemistry and physics when learning biology concepts (e.g., apply physics concepts when learning about microscopy or mass spectrometry, apply chemistry knowledge when describing molecular affinity). |
| | | Describe influences of physical forces or chemical interactions in biological systems (e.g., flux in electrophysiology, bulk flow in respiration, hydrogen bonding in enzyme/substrate binding, chemosensation or biomechanics in plant-pollinator interactions). |
| | | Use math to model biological systems. |
| | Build models or explanations of simple biological processes that include concepts from other STEM disciplines or multiple fields of biology. | Build a concept map connecting ideas from multiple disciplines in the context of a biological system (e.g., the kinetics, biochemistry, and functions of catalysis; the chemistry and physics of membrane potentials; the effects of abiotic factors and symbiotes on plant productivity). |
| | | Sketch models or write explanations of biological systems that incorporate concepts from multiple disciplines, including how components interact across scales (i.e. atoms, molecules, cellular structures, organs, organisms, ecosystems). |
| **INTERDISCIPLINARY PROBLEM SOLVING Consider interdisciplinary solutions to real-world problems.** | Describe examples of real-world problems that are too complex to be solved by applying biological approaches alone. | Reflect on and propose solutions to case studies of complex problems with important societal consequences (e.g., ocean acidification, malaria epidemic, ecological impacts of urbanization). |
| | | Identify stories in the news or other popular media that include the contributions of experts from multiple disciplines. |
| | | Attend and summarize seminars from local faculty engaged in interdisciplinary research. |
| | Suggest how collaborators in STEM and non-STEM disciplines could contribute to solutions of real-world problems. | Discuss and describe the contributions of different stakeholders in actual policy proposals (e.g., clean water initiatives, carbon taxes, vaccination requirements). |
| | | Identify gaps in own knowledge (e.g., as part of a case on diabetes, write questions for a chemist to improve understanding of symptoms and drug treatments). |

#### Expanded BioSkills Guide: Interdisciplinary Nature of Science

| Program-Level Learning Outcomes | Course-Level Learning Outcomes | Examples |
|---|---|---|
| **INTERDISCIPLINARY PROBLEM SOLVING Consider interdisciplinary solutions to real-world problems.** ***(continued)*** | Be able to explain biological concepts, data, and methods, including their limitations, using language understandable by collaborators in other disciplines. | Identify and define terms that can have different meanings in different contexts (e.g., regression, acid/base, energy). |
| | | Share data collection and analysis tasks with students from multiple disciplines (e.g., chemistry and biology students work together on a drug discovery project). |
| | | Define constraints and parameters of a system to be used by colleagues in other disciplines (e.g., biology and computer science students collaborate to write computer scripts to analyze genomics data). |
| | | Teach a student in another major about your research topic. |

### Expanded BioSkills Guide: Communication & Collaboration

| Program-Level Learning Outcomes | Course-Level Learning Outcomes | Examples |
|---|---|---|
| **COMMUNICATION Share ideas, data, and findings with others clearly and accurately.** | Use appropriate language and style to communicate science effectively to targeted audiences (e.g., general public, biology experts, collaborators in other disciplines). | Adjust level of detail in data presentations depending on scientific audience (e.g., lab meeting for experts vs. class presentations for general biology audience). |
| | | Build educational blogs, pamphlets, Wikipedia pages or magazine articles for audiences outside of the classroom. |
| | | Write evidence-based policy recommendations for private or government agencies (e.g., Nature Conservancy, State Fish & Wildlife, Food & Drug Administration). |
| | | Create tailored presentations or documents to inform the general public (e.g., bird-watching groups, museum visitors) about important new biological findings, avoiding overly sensational language. |
| | Use a variety of modes to communicate science (e.g., oral, written, visual). | Present data orally with supporting poster, slides, or chalkboard sketches. |
| | | Write blogs, essays, papers, or pamphlets to communicate findings. |
| | | Give mini-lectures in the classroom. |
| | | Write a scientific abstract, research paper, senior thesis, or grant proposal. |
| | | Present data as infographics. |
| **COLLABORATION Work productively in teams with people who have diverse backgrounds, skill sets, and perspectives.** | Work with teammates to establish and periodically update group plans and expectations (e.g., team goals, project timeline, rules for group interactions, individual and collaborative tasks). | Delegate tasks to accomplish larger projects (e.g., team-based learning, many-hands data collection, collaborative presentations). |
| | | Prepare a group contract establishing norms and expectations (e.g., modes of communication, frequency of meetings, paths for feedback). |
| | | Set aside team time to discuss progress and reorganize work distribution and decision-making processes as needed. |
| | Elicit, listen to, and incorporate ideas from teammates with different perspectives and backgrounds. | Learn from teammates in small groups (e.g., jigsaw reading of journal articles, think-pair-share). |
| | | Ask clarifying questions from partner. |
| | | Monitor group conversations for equitable participation. |
| | | Take the role of different stakeholders and have a discussion about a policy issue using scientific evidence. |

#### Expanded BioSkills Guide: Communication & Collaboration

| Program-Level Learning Outcomes | Course-Level Learning Outcomes | Examples |
|---|---|---|
| **COLLABORATION Work productively in teams with people who have diverse backgrounds, skill sets, and perspectives. *(continued)*** | Work effectively with teammates to complete projects. | Establish mode of communication and resource sharing that works for all group members (e.g., emails, online file sharing applications, in person meetings). |
| | | Seek outside help from instructor or TA when group reaches an impasse. |
| | | Meet before presentations or deadlines to integrate individual products into a cohesive whole. |
| | | Plan multiple opportunities to exchange drafts and share progress updates for whole-group feedback. |
| **COLLEGIAL REVIEW Provide and respond to constructive feedback in order to improve individual and team work.** | Evaluate feedback from others and revise work or behavior appropriately. | Modify posters, presentations, or papers based on comments by peers and instructors. |
| | | Listen to and weigh alternative points of view. |
| | | Ask others for specific types of feedback based on self-assessed weaknesses of work. |
| | Critique others' work and ideas constructively and respectfully. | Peer review papers and presentations, providing feedback on both content and style. |
| | | Where appropriate, use existing methods to formalize feedback and maximize likelihood of use (e.g., 'compliment sandwich' or 'keep-quit-start'). |
| | | Evaluate performance of other team members and make constructive suggestions. |
| **METACOGNITION Reflect on your own learning, performance, and achievements.** | Evaluate your own understanding and skill level. | Compare your responses on an exam or assignment to a key and identify areas for improvement. |
| | | Write down the "muddiest point" from a class or study session. |
| | | Score your work as part of a team project. |
| | Assess personal progress and contributions to your team and generate a plan to change your behavior as needed. | Use practice test or midterm outcomes to determine how to modify study strategies. |
| | | Develop time management strategies to meet competing deadlines. |
| | | Set and revisit goals to reflect on personal growth (How have you improved? How can you ensure you reach your goal?). |
| | | Monitor and adjust behavior based on informal and formal feedback from group members. |

### Expanded BioSkills Guide: Science & Society

| Program-Level Learning Outcomes | Course-Level Learning Outcomes | Examples |
|---|---|---|
| **ETHICS Demonstrate the ability to critically analyze ethical issues in the conduct of science.** | Identify and evaluate ethical considerations (e.g., use of animal or human subjects, conflicts of interest, confirmation bias) in a given research study. | Discuss animal welfare and rights in biomedical research and agriculture. |
| | | Outline relevant ethical considerations for a published study. |
| | | Complete appropriate ethics training before conducting research. |
| | | Identify relevant ethical considerations for experimental design before beginning an independent or course-based research study, and discuss appropriate accommodations. |
| | Critique how ethical controversies in biological research have been and can continue to be addressed by the scientific community. | Discuss historical cases of scientific misconduct or controversy (e.g., Rosalind Franklin and the history of women in science, Henrietta Lacks and informed consent). |
| | | Debate current status and proposed solutions to modern scientific controversies (e.g., call for moratorium on human gene editing, the sharing of pathogenic virus sequences). |
| | | Evaluate strengths and weaknesses of existing paths for ethics review (e.g., IRB and IACUC review processes). |
| | | Compare research policies and guidelines (e.g., stem cell research, embryonic gene editing) in the US with other countries. |
| **SOCIETAL INFLUENCES Consider the potential impacts of outside influences (historical, cultural, political, technological) on how science is practiced.** | Describe examples of how scientists' backgrounds and biases can influence science and how science is enhanced through diversity. | Research and summarize the contributions of scientists from diverse backgrounds. |
| | | Discuss cases where diversity in science led to innovation (e.g., maternal effect, sexual selection). |
| | | Reflect on how scientists' worldviews affect their interpretations (e.g., "ladder of life" model of evolution, Earth as the center of the universe). |
| | | Discuss the connections between social justice and science (e.g., the history of research on the genetic basis of race, funding for neglected tropical diseases). |
| | Identify and describe how systemic factors (e.g., socioeconomic, political) affect how and by whom science is conducted. | Critique the strengths and weaknesses of peer review for publication. |
| | | Describe cases where a new technology changed the types of data that can be collected and therefore the scientific questions that can be answered (e.g., DNA sequencing, imaging technologies, large public databases). |
| | | Listen to stories from the MeToo STEM movement and reflect on how scientific culture can affect who pursues and remains in science. |
| | | Discuss ways that education and hiring practices might be changed to lessen or eliminate opportunity gaps for underrepresented groups in science. |
| | | Discuss how funding determines what research is prioritized (e.g., neglected tropical diseases, climate change). |

### Expanded BioSkills Guide: Science & Society

| Program-Level Learning Outcomes | Course-Level Learning Outcomes | Examples |
|---|---|---|
| **SCIENCE'S IMPACT ON SOCIETY** **Apply scientific reasoning in daily life and recognize the impacts of science on a local and global scale.** | Apply evidence-based reasoning and biological knowledge in daily life (e.g., consuming popular media, deciding how to vote). | Practice skepticism when consuming popular media about scientific or non-scientific topics. |
| | | Reflect on how science is applied in personal decisions about health, use of technology, and interactions with the environment. |
| | | Share relevant biology knowledge with friends and relatives during conversations about current events or health decisions. |
| | Use examples to describe the relevance of science in everyday experiences. | Notice and describe local plant ecology, the biology of food and nutrition, representations and reports of science in popular culture. |
| | | Evaluate how evidence is used in government policy decisions (e.g., subsidies in agriculture, public health policy, funding of renewable energy). |
| | Identify and describe the broader societal impacts of biological research on different stakeholders. | Discuss past cases of biased research designs having negative repercussions (e.g., pharmacological trials on white male patients used to inform dosage in all populations). |
| | | Reflect on unanticipated impacts of scientific advances (e.g., environmental policy, genetic engineering, personal genomics). |
| | | Consider the perspectives of multiple stakeholders as part of lessons about current societal issues (e.g., comparing DNA found at crime scenes to genealogical records, impacts of human disturbance on wildlife health, climate-related habitat change). |
| | Describe the roles scientists have in facilitating public understanding of science. | Read and describe the purpose of various modes of presenting science for a general audience (e.g., Science section of the New York Times, museum exhibits, scientist interviews on news shows or podcasts). |
| | | Write summaries of new biological findings for a general audience, including a discussion of why they should care. |
| | | Discuss the importance of political advocacy as part of the role of professional scientists (e.g., voice of scientists in debates on vaccines, climate change, or misconceptions about race and genetics). |
| | | Share biology knowledge while volunteering for non-profit organizations or participating in local community meetings. |

**Supplemental Material 3. Supplemental Figures.**

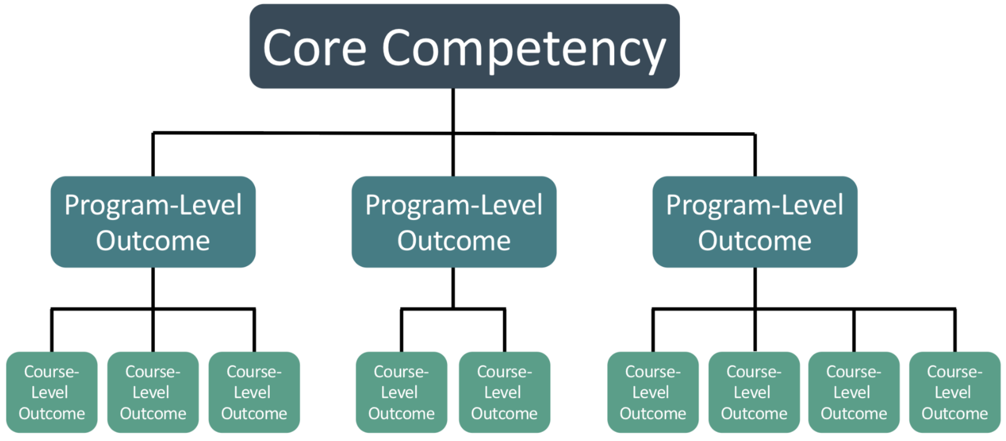


**Supplemental Figure 1. Nested BioSkills Guide structure.**
The BioSkills Guide has a two-tiered structure. Each of the six core competencies contains 2-6 program-level learning outcomes (20 total), and each program-level outcome contains 2-6 course-level outcomes (57 total).

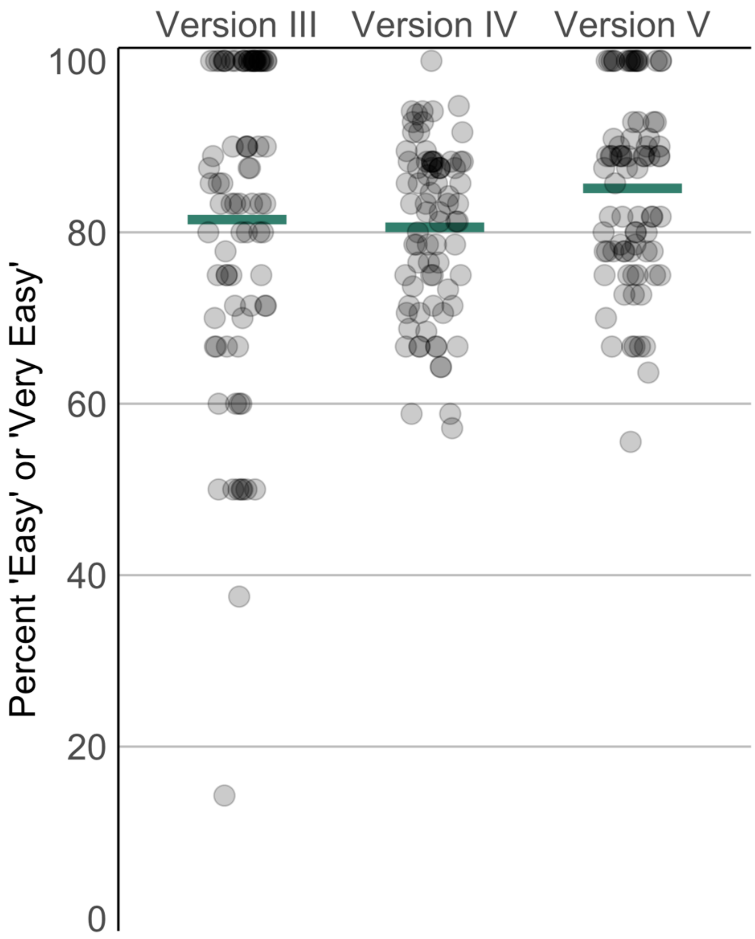


**Supplemental Figure 2. Change in 'ease of understanding' ratings over time.**
For each learning outcome, ratings were summarized by calculating the percent of respondents who selected 'Easy' or 'Very Easy'. Horizontal lines indicate means. Points are jittered to reveal distribution. 'Ease of understanding' questions were not included in validation phase surveys.

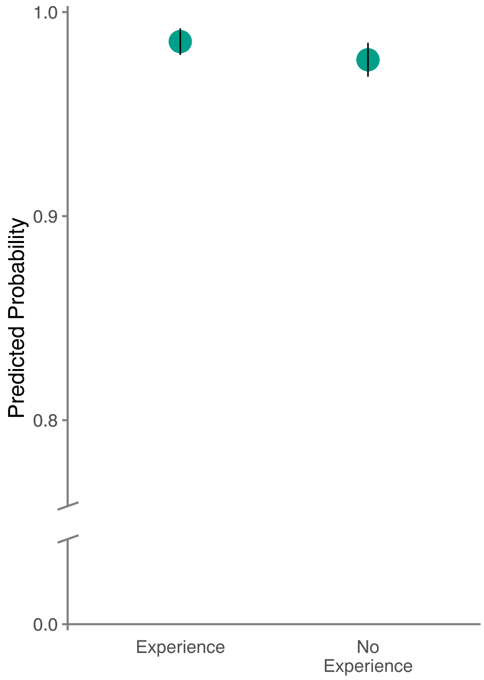


**Supplemental Figure 3. Effect of experience with ecology or evolutionary biology research on support for learning outcomes.**

Experience in ecology or evolutionary biology research was retained as a main effect in the best fitting model for RQ2b. Predicted probabilities are shown. Vertical lines indicate 95% confidence intervals. Note that y-axis is truncated.

**Supplemental Material 4. Supplemental Tables.**

**Supplemental Table 1. Summary of revisions.**

[a] The number of learning outcomes that were removed, added, or reworded is shown for each round of revision. Rewordings included substantial rewrites and single word changes. The total number of outcomes (both program- and course-level) in the starting draft is also shown (i.e. there were 78 outcomes in Version IV).

[b] One outcome underwent minor revision after the national validation survey. Details of editing are in Methods.

| | Number of Outcomes [a] | | | |
|---|---|---|---|---|
| **Round** | **Before Revision** | **Removed** | **Added** | **Reworded** |
| **Version I -> Version II** | 86 | 1 | 1 | 30 |
| **Version II -> Version III** | 86 | 7 | 2 | 57 |
| **Version III -> Version IV** | 81 | 6 | 3 | 57 |
| **Version IV -> Version V** | 78 | 4 | 6 | 64 |
| **Version V -> Version VI** | 80 | 3 | 0 | 29 |
| **Version VI -> Final** | 77 | 0 | 0 | 1 |
| **Final Version [b]** | 77 | 0 | 0 | 1 |

**Supplemental Table 2. Summary of self-reported demographics of survey respondents during development phase.**

[a] Count and percent of total (out of 93, unless otherwise noted) of respondents who selected indicated response, aggregated across all three surveys from the development phase. Unknown indicates that respondent did not answer that question.

[b] Mean importance rating (1 = 'Very Unimportant', 5 = 'Very Important') was calculated across all outcomes reviewed by respondents who selected that response.

[c] Demographic questions were revised slightly between review of Version III and review of Version IV. These demographic characteristics can only be determined from respondents who reviewed Version IV or V (n=72).

[d] Because of survey revision, this demographic characteristic can only be determined from respondents who reviewed Version III (n=21).

[e] These questions allowed respondents to select all that apply, so percentages do not sum to 100.

[f] These characteristics were inferred from responses to questions about Graduate Training Subdiscipline and Current Research Subdiscipline.

| Demographic | Response | n (%) [a] | Mean Importance [b] |
|---|---|---|---|
| Institution Type | Associate's Granting | 10 (10.8%) | 4.6 |
| | Bachelor's Granting | 9 (9.7%) | 4.4 |
| | Master's Granting | 18 (19.4%) | 4.6 |
| | Doctoral Granting | 42 (45.2%) | 4.4 |
| | Other | 1 (1.1%) | 4.3 |
| | Unknown | 13 (14%) | 4.3 |
| Current Position | Graduate Student | 3 (3.2%) | 4.7 |
| | Postdoc | 9 (9.7%) | 4.5 |
| | Lecturer/Instructor | 17 (18.3%) | 4.5 |
| | Assistant/Associate/Full Professor | 44 (47.3%) | 4.5 |
| | Staff | 2 (2.2%) | 4.4 |
| | Other | 4 (4.3%) | 4.7 |
| | Unknown | 14 (15.1%) | 4.3 |
| Job Responsibility [c] | Teaching | 25 (34.7%) | 4.6 |
| | Research | 10 (13.9%) | 4.5 |
| | Teaching & Research Equally | 14 (19.4%) | 4.5 |
| | Other | 5 (6.9%) | 4.3 |
| | Unknown | 18 (25%) | 4.5 |
| Course Levels Taught [e] | Non-Majors Lower (100-200)[2] | 27 (17.2%) | 4.5 |
| | Majors Lower (100-200) | 51 (32.5%) | 4.5 |
| | Upper (300-400) | 41 (26.1%) | 4.6 |
| | Graduate (500+) | 16 (10.2%) | 4.5 |
| | Unknown | 22 (14%) | 4.4 |
| Teaching Subdiscipline [c] | Molecular/Cell/Development (MCD) | 12 (16.7%) | 4.5 |
| | Physiology | 6 (8.3%) | 4.6 |
| | Ecology/Evolution | 12 (16.7%) | 4.7 |
| | General | 11 (15.3%) | 4.3 |
| | Other | 13 (18.1%) | 4.4 |
| | Unknown | 18 (25%) | 4.5 |

| | | | |
|---|---|---|---|
| **Expertise Subdiscipline [d]** | MCD | 8 (28.6%) | 4.6 |
| | Physiology | 3 (10.7%) | 4.4 |
| | Ecology/Evolution | 6 (21.4%) | 4.3 |
| | Discipline-Based Education Research (DBER) | 5 (17.9%) | 4.5 |
| | Other | 2 (7.1%) | 4.5 |
| | Unknown | 4 (14.3%) | 4.1 |
| **Current Research Subdiscipline [ce]** | No Current Research | 8 (10.1%) | 4.5 |
| | MCD | 6 (7.6%) | 4.4 |
| | Physiology | 2 (2.5%) | 4.7 |
| | Ecology/Evolution | 5 (6.3%) | 4.8 |
| | DBER | 37 (46.8%) | 4.5 |
| | Other | 3 (3.8%) | 4.5 |
| | Unknown | 18 (22.8%) | 4.5 |
| **Graduate Training Subdiscipline [ce]** | MCD | 12 (14.6%) | 4.5 |
| | Physiology | 8 (9.8%) | 4.4 |
| | Ecology/Evolution | 19 (23.2%) | 4.6 |
| | DBER | 10 (12.2%) | 4.5 |
| | Other | 15 (18.3%) | 4.4 |
| | Unknown | 18 (22%) | 4.5 |
| **Currently Engaged in Biology Research [cf]** | Not Currently Engaged in Biology Research | 39 (54.2%) | 4.5 |
| | Current Research in Biology | 15 (20.8%) | 4.6 |
| | Unknown | 18 (25%) | 4.5 |
| **DBER Experience [f]** | No DBER Experience | 26 (28%) | 4.5 |
| | DBER Experience | 45 (48.4%) | 4.5 |
| | Unknown | 22 (23.7%) | 4.4 |
| **Familiarity with Vision and Change** | Extremely Familiar | 28 (30.1%) | 4.6 |
| | Very Familiar | 24 (25.8%) | 4.4 |
| | Somewhat Familiar | 13 (14%) | 4.5 |
| | Slightly Familiar | 2 (2.2%) | 4.4 |
| | Not at all Familiar | 3 (3.2%) | 4.1 |
| | Unknown | 23 (24.7%) | 4.4 |
| **Previous Involvement [c]** | Previously Involved | 10 (13.9%) | 4.5 |
| | New | 44 (61.1%) | 4.5 |
| | Unknown | 18 (25%) | 4.5 |

**Supplemental Table 3. Counts of unique participants and institutions, by institution type.**

[a] Institution classification of all participants was determined by matching institution name with Carnegie classification dataset (Indiana University Center for Postsecondary Research, 2016).

[b] Total n and distribution for development phase is distinct from Supplemental Table 2 because these data also include workshop, round table, and interview participants.

| Phase | Institution Type [a] | Participants, n (% of unique) | Institutions, n (% of unique) |
|---|---|---|---|
| **Development [b]** | Associate's | 37 (14.4%) | 24 (27.6%) |
| | Bachelor's | 13 (5.1%) | 10 (11.5%) |
| | Master's | 31 (12.1%) | 16 (18.4%) |
| | Doctoral | 95 (37%) | 29 (33.3%) |
| | International | 37 (14.4%) | 4 (4.6%) |
| | Other | 4 (1.6%) | 4 (4.6%) |
| | Unknown | 40 (15.6%) | NA |
| **Validation** | Associate's | 86 (20.6%) | 59 (26.2%) |
| | Bachelor's | 50 (12%) | 36 (16%) |
| | Master's | 67 (16.1%) | 43 (19.1%) |
| | Doctoral | 116 (27.8%) | 77 (34.2%) |
| | International | 7 (1.7%) | 6 (2.7%) |
| | Other | 5 (1.2%) | 4 (1.8%) |
| | Unknown | 86 (20.6%) | NA |

**Supplemental Table 4. Summary of learning outcome importance ratings across development and validation phases.**

[a] Importance ratings (1 = 'Very Unimportant', 5 = 'Very Important') were individually summarized for each learning outcome by calculating percent support and the mean rating. Then, minimum, maximum, and mean were calculated across all learning outcomes for both summary measures.

[b] 'n total' is the number of unique survey respondents who participated in that survey. 'n outcome' is the number of respondents who reviewed each individual outcome (respondents were randomly assigned a subset of outcomes). 'n observations' is the number of rating data points collected across all outcomes and respondents.

| | | Percent support [a] | | | Mean rating [a] | | | n [b] | | |
|---|---|---|---|---|---|---|---|---|---|---|
| **Phase** | Round | min | max | mean | min | max | mean | total | outcome | observations |
| **Develop-ment** | Version III | 16.7 | 100 | 85.9 | 2.7 | 5 | 4.4 | 21 | 5-10 | 618 |
| | Version IV | 50 | 100 | 92.8 | 3.4 | 4.9 | 4.5 | 45 | 12-19 | 1197 |
| | Version V | 72.7 | 100 | 94.4 | 3.8 | 5 | 4.5 | 27 | 8-14 | 786 |
| **Validation** | Pilot | 72.7 | 100 | 94.9 | 3.9 | 4.9 | 4.5 | 20 | 11-12 | 905 |
| | National | 73.5 | 99.5 | 91.7 | 4 | 4.9 | 4.5 | 397 | 200-225 | 16667 |
| | **Combined** | **74.3** | **99.6** | **91.9** | **4** | **4.9** | **4.5** | **417** | **211-237** | **17572** |

**Supplemental Table 5. Summary of descriptive statistics of learning outcome 'ease of understanding' ratings across development phase.**

[a] 'Ease of understanding' ratings (1 = 'Very Difficult', 5 = 'Very Easy') were individually summarized for each learning outcome by calculating the percent of respondents who selected 'Easy' or 'Very Easy' and the mean rating. Then, minimum, maximum, and mean were calculated across all outcomes for both summary measures. Participant counts can be seen in Supplemental Table 4.

[b] 'Ease of understanding' questions were not included in validation phase surveys.

| | Percent 'Easy' or 'Very Easy' [a] | | | Mean rating [a] | | |
|---|---|---|---|---|---|---|
| **Round [b]** | min | max | mean | min | max | mean |
| **Version III** | 14.3 | 100 | 81.5 | 2.6 | 5 | 4.3 |
| **Version IV** | 57.1 | 100 | 80.6 | 3.4 | 4.8 | 4.2 |
| **Version V** | 55.6 | 100 | 85.1 | 3.6 | 4.8 | 4.4 |

**Supplemental Table 6. Self-reported demographics of survey respondents during validation phase.**

[a] Number and percent (out of 417) of unique respondents who selected indicated response, aggregated across both surveys from the validation phase. Unknown indicates that respondent did not answer that question.

[b] Mean importance rating (1 = 'Very Unimportant', 5 = 'Very Important') was calculated across all outcomes reviewed by respondents who selected that response.

[c] These questions allowed respondents to select all that apply, so percentages do not sum to 100.

[d] These characteristics were inferred from responses to questions about Graduate Training Subdiscipline and Current Research Subdiscipline.

| Demographic | Response | n (%) [a] | Mean Importance [b] |
|---|---|---|---|
| **Institution Type** | Associate's Granting | 92 (22.1%) | 4.5 |
| | Bachelor's Granting | 96 (23%) | 4.5 |
| | Master's Granting | 51 (12.2%) | 4.5 |
| | Doctoral Granting | 121 (29%) | 4.5 |
| | Other | 7 (1.7%) | 4.4 |
| | Unknown | 50 (12%) | 4.4 |
| **Current Position** | Graduate Student | 5 (1.2%) | 4.5 |
| | Postdoc | 5 (1.2%) | 4.5 |
| | Lecturer/Instructor | 83 (19.9%) | 4.5 |
| | Assistant/Associate/Full Professor | 253 (60.7%) | 4.5 |
| | Staff | 5 (1.2%) | 4.6 |
| | Other | 16 (3.8%) | 4.6 |
| | Unknown | 50 (12%) | 4.4 |
| **Job Responsibility** | Teaching | 258 (61.9%) | 4.5 |
| | Research | 10 (2.4%) | 4.5 |
| | Teaching & Research Equally | 79 (18.9%) | 4.4 |
| | Other | 20 (4.8%) | 4.6 |
| | Unknown | 50 (12%) | 4.4 |
| **Teaching Subdiscipline** | Molecular/Cell/Development | 108 (25.9%) | 4.4 |
| | Physiology | 49 (11.8%) | 4.4 |
| | Ecology/Evolution | 65 (15.6%) | 4.5 |
| | General | 103 (24.7%) | 4.5 |
| | Other | 38 (9.1%) | 4.4 |
| | Unknown | 54 (12.9%) | 4.4 |
| **Course Levels [c]** | Non-Majors Lower (100-200) | 196 (47%) | 4.5 |
| | Majors Lower (100-200) | 265 (63.5%) | 4.5 |
| | Upper (300-400) | 221 (53%) | 4.5 |
| | Graduate (500+) | 56 (13.4%) | 4.4 |
| | Unknown | 55 (13.2%) | 4.4 |
| **Course Levels, Aggregated** | Lower-Level Only (100-200) | 129 (30.9%) | 4.5 |
| | Advanced Only (300-500+) | 40 (9.6%) | 4.5 |
| | Lower & Advanced | 193 (46.3%) | 4.5 |
| | Unknown | 55 (13.2%) | 4.4 |

| | | | |
|---|---|---|---|
| **Current Research Subdiscipline [c]** | No Current Research | 86 (20.6%) | 4.5 |
| | Molecular/Cell/Development (MCD) | 84 (20.1%) | 4.4 |
| | Physiology | 17 (4.1%) | 4.5 |
| | Ecology/Evolution | 68 (16.3%) | 4.5 |
| | Discipline-Based Education Research (DBER) | 112 (26.9%) | 4.5 |
| | Unknown | 65 (15.6%) | 4.4 |
| **Current Research Subdiscipline, Aggregated** | No Current Research | 84 (20.1%) | 4.5 |
| | MCD Only | 61 (14.6%) | 4.4 |
| | Physiology Only | 11 (2.6%) | 4.5 |
| | Ecology/Evolution Only | 53 (12.7%) | 4.5 |
| | DBER Only | 90 (21.6%) | 4.5 |
| | Other Only | 21 (5%) | 4.5 |
| | More than 2 Subdisciplines | 6 (1.4%) | 4.5 |
| | MCD & Physiology | 1 (0.2%) | 4.8 |
| | MCD & Ecology/Evolution | 5 (1.2%) | 4.3 |
| | MCD & DBER | 12 (2.9%) | 4.4 |
| | Physiology & Ecology/Evolution | 2 (0.5%) | 4.6 |
| | Physiology & DBER | 1 (0.2%) | 4.7 |
| | Ecology/Evolution & DBER | 5 (1.2%) | 4.6 |
| | Unknown | 65 (15.6%) | 4.4 |
| **Graduate Training Subdiscipline [c]** | MCD | 182 (43.6%) | 4.5 |
| | Physiology | 40 (9.6%) | 4.5 |
| | Ecology/Evolution | 126 (30.2%) | 4.5 |
| | DBER | 17 (4.1%) | 4.6 |
| | Other | 40 (9.6%) | 4.4 |
| | Unknown | 52 (12.5%) | 4.4 |
| **Graduate Training Subdiscipline, Aggregated** | MCD Only | 165 (39.6%) | 4.4 |
| | Physiology Only | 26 (6.2%) | 4.5 |
| | Ecology/Evolution Only | 115 (27.6%) | 4.5 |
| | DBER Only | 11 (2.6%) | 4.5 |
| | Other Only | 24 (5.8%) | 4.3 |
| | More than 2 Subdisciplines | 2 (0.5%) | 4.8 |
| | MCD & Physiology | 8 (1.9%) | 4.7 |
| | MCD & Ecology/Evolution | 5 (1.2%) | 4.6 |
| | MCD & DBER | 3 (0.7%) | 4.6 |
| | Physiology & Ecology/Evolution | 3 (0.7%) | 4.5 |
| | Physiology & DBER | 1 (0.2%) | 4.5 |
| | Ecology/Evolution & DBER | 2 (0.5%) | 4.8 |
| | Unknown | 52 (12.5%) | 4.4 |
| **DBER Experience [d]** | No DBER Experience | 236 (56.6%) | 4.5 |
| | DBER Experience | 116 (27.8%) | 4.5 |

| | | | |
|---|---|---|---|
| | Unknown | 65 (15.6%) | 4.4 |
| **Currently Engaged in Biology Research** [d] | Not Currently Engaged in Biology Research | 170 (40.8%) | 4.5 |
| | Current Research in Biology | 182 (43.6%) | 4.5 |
| | Unknown | 65 (15.6%) | 4.4 |
| **Familiarity with Vision and Change** | Extremely Familiar | 110 (26.4%) | 4.5 |
| | Very Familiar | 146 (35%) | 4.4 |
| | Somewhat Familiar | 68 (16.3%) | 4.5 |
| | Slightly Familiar | 16 (3.8%) | 4.2 |
| | Not at all Familiar | 26 (6.2%) | 4.4 |
| | Unknown | 51 (12.2%) | 4.4 |
| **Gender** | Female | 241 (57.8%) | 4.5 |
| | Male | 120 (28.8%) | 4.4 |
| | Other | 2 (0.5%) | 4.7 |
| | Unknown | 54 (12.9%) | 4.4 |
| **Previous Involvement** | Previously Involved | 29 (7%) | 4.5 |
| | New | 334 (80.1%) | 4.5 |
| | Unknown | 54 (12.9%) | 4.4 |

**Supplemental Table 7. Descriptive statistics of importance ratings for all outcomes during validation.**

[a] Importance ratings (1 = 'Very Unimportant', 5 = 'Very Important') for each learning outcome were summarized by calculating percent support (percent 'Important' or 'Very Important' out of all ratings), mean, maximum, and minimum ratings. n=211-237 per outcome.

[b] This is the only outcome that was revised after the national validation survey. Details of editing are in Methods.

| Process of Science | | | | | |
|---|---|---|---|---|---|
| **Outcome #** | **Outcome** | **% Support [a]** | **Mean** | **Max** | **Min** |
| **1** | SCIENTIFIC THINKING Explain how science generates knowledge of the natural world. | 97 | 4.8 | 5 | 1 |
| **1.1** | Explain how scientists use inference and evidence-based reasoning to generate knowledge. | 97.4 | 4.6 | 5 | 1 |
| **1.2** | Describe the iterative nature of science and how new evidence can lead to the revision of scientific knowledge. | 97.9 | 4.7 | 5 | 1 |
| **2** | INFORMATION LITERACY Locate, interpret, and evaluate scientific information. | 98.3 | 4.8 | 5 | 1 |
| **2.1** | Find and evaluate the credibility of a variety of sources of scientific information, including popular science media and scientific journals. | 97 | 4.7 | 5 | 1 |
| **2.2** | Interpret, summarize, and evaluate evidence in primary literature. | 94.8 | 4.6 | 5 | 1 |
| **2.3** | Evaluate claims in scientific papers, popular science media, and other sources using evidence-based reasoning. | 97 | 4.7 | 5 | 1 |
| **3** | QUESTION FORMULATION Pose testable questions and hypotheses to address gaps in knowledge. | 95.7 | 4.6 | 5 | 1 |
| **3.1** | Recognize gaps in our current understanding of a biological system or process and identify what specific information is missing. | 83 | 4.1 | 5 | 1 |
| **3.2** | Develop research questions based on your own or others' observations. | 89.8 | 4.3 | 5 | 1 |
| **3.3** | Formulate testable hypotheses and state their predictions. | 95.3 | 4.6 | 5 | 1 |
| **4** | STUDY DESIGN Plan, evaluate, and implement scientific investigations. | 93.6 | 4.5 | 5 | 2 |
| **4.1** | Compare the strengths and limitations of various study designs. | 87.2 | 4.2 | 5 | 2 |
| **4.2** | Design controlled experiments, including plans for analyzing the data. | 91.5 | 4.5 | 5 | 2 |
| **4.3** | Execute protocols and accurately record measurements and observations. | 93.6 | 4.5 | 5 | 1 |
| **4.4** | Identify methodological problems and suggest solutions or alternative approaches. | 83.8 | 4.1 | 5 | 1 |
| **4.5** | Evaluate and suggest best practices for responsible research conduct (e.g., lab safety, record keeping, proper citation of sources). | 82 | 4.2 | 5 | 2 |
| **5** | DATA INTERPRETATION & EVALUATION Interpret, evaluate, and draw conclusions from data in order to make evidence-based arguments about the natural world. | 98.3 | 4.8 | 5 | 1 |
| **5.1** | Analyze data, summarize resulting patterns, and draw appropriate conclusions. | 99.1 | 4.8 | 5 | 1 |
| **5.2** | Describe sources of error and uncertainty in data. | 93.6 | 4.4 | 5 | 1 |
| **5.3** | Make evidence-based arguments using your own and others' findings. | 97.8 | 4.7 | 5 | 1 |
| **5.4** | Relate conclusions to original hypothesis, consider alternative hypotheses, and suggest future research directions based on findings. | 95.7 | 4.6 | 5 | 1 |
| **6** | DOING RESEARCH Apply science process skills to address a research question in a course-based or independent research experience. | 93.2 | 4.4 | 5 | 1 |

| Quantitative Reasoning | | | | | |
|---|---|---|---|---|---|
| **Outcome #** | **Outcome** | **% Support** | **Mean** | **Max** | **Min** |
| **1** | NUMERACY Use basic mathematics (e.g., algebra, probability, unit conversions) in biological contexts. | 98.6 | 4.8 | 5 | 2 |
| **1.1** | Perform basic calculations (e.g., percentages, frequencies, rates, means). | 99.6 | 4.9 | 5 | 3 |
| **1.2** | Select and apply appropriate equations (e.g., Hardy-Weinberg, Nernst, Gibbs free energy) to solve problems. | 86.5 | 4.3 | 5 | 2 |
| **1.3** | Interpret and manipulate mathematical relationships (e.g., scale, ratios, units) to make quantitative comparisons. | 98.2 | 4.7 | 5 | 3 |
| **1.4** | Use probability and understanding of biological variability to reason about biological processes and statistical analyses. | 96 | 4.6 | 5 | 2 |
| **1.5** | Use rough estimates informed by biological knowledge to check quantitative work. | 92.8 | 4.5 | 5 | 1 |

| | | | | | |
|---|---|---|---|---|---|
| **1.6** | Describe how quantitative reasoning helps biologists understand the natural world. | 91.9 | 4.5 | 5 | 2 |
| **2** | QUANTITATIVE & COMPUTATIONAL DATA ANALYSIS Apply the tools of graphing, statistics, and data science to analyze biological data. | 98.2 | 4.8 | 5 | 3 |
| **2.1** | Record, organize, and annotate simple data sets. | 98.7 | 4.8 | 5 | 3 |
| **2.2** | Create and interpret informative graphs and other data visualizations. | 99.6 | 4.9 | 5 | 3 |
| **2.3** | Select, carry out, and interpret statistical analyses. | 95.9 | 4.5 | 5 | 2 |
| **2.4** | Describe how biologists answer research questions using databases, large data sets, and data science tools. | 87.9 | 4.3 | 5 | 2 |
| **2.5** | Interpret the biological meaning of quantitative results. | 99.1 | 4.7 | 5 | 3 |

#### Modeling

| Outcome # | Outcome | % Support | Mean | Max | Min |
|---|---|---|---|---|---|
| 1 | PURPOSE OF MODELS Recognize the important roles that scientific models, of many different types (conceptual, mathematical, physical, etc.), play in predicting and communicating biological phenomena. | 93.9 | 4.5 | 5 | 2 |
| **1.1** | Describe why biologists use simplified representations (models) when solving problems and communicating ideas. | 88.3 | 4.3 | 5 | 2 |
| **1.2** | Given two models of the same biological process or system, compare their strengths, limitations, and assumptions. | 84.1 | 4.3 | 5 | 2 |
| **2** | MODEL APPLICATION Make inferences and solve problems using models and simulations. | 88.8 | 4.3 | 5 | 2 |
| **2.1** | Summarize relationships and trends that can be inferred from a given model or simulation. | 93.5 | 4.3 | 5 | 1 |
| **2.2** | Use models and simulations to make predictions and refine hypotheses. | 89.8 | 4.3 | 5 | 1 |
| **3** | MODELING Build and evaluate models of biological systems. | 75.5 | 4 | 5 | 1 |
| **3.1** | Build and revise conceptual models to propose how a biological system or process works. [b] | 86.4 | 4.2 | 5 | 1 |
| **3.2** | Identify important components of a system and describe how they influence each other (e.g., positively or negatively). | 93.9 | 4.5 | 5 | 1 |
| **3.3** | Evaluate conceptual, mathematical, or computational models by comparing their predictions with empirical data. | 82.6 | 4.2 | 5 | 1 |

#### Interdisciplinary Nature of Science

| Outcome # | Outcome | % Support | Mean | Max | Min |
|---|---|---|---|---|---|
| **1** | CONNECTING SCIENTIFIC KNOWLEDGE Integrate concepts across other STEM disciplines (e.g., chemistry, physics) and multiple fields of biology (e.g., cell biology, ecology). | 95.1 | 4.5 | 5 | 1 |
| **1.1** | Given a biological problem, identify relevant concepts from other STEM disciplines or fields of biology. | 89.4 | 4.2 | 5 | 1 |
| **1.2** | Build models or explanations of simple biological processes that include concepts from other STEM disciplines or multiple fields of biology. | 82.4 | 4.1 | 5 | 1 |
| **2** | INTERDISCIPLINARY PROBLEM SOLVING Consider interdisciplinary solutions to real-world problems. | 88.4 | 4.3 | 5 | 1 |
| **2.1** | Describe examples of real-world problems that are too complex to be solved by applying biological approaches alone. | 74 | 4 | 5 | 1 |
| **2.2** | Suggest how collaborators in STEM and non-STEM disciplines could contribute to solutions of real-world problems. | 74.3 | 4 | 5 | 1 |
| **2.3** | Be able to explain biological concepts, data, and methods, including their limitations, using language understandable by collaborators in other disciplines. | 91.6 | 4.5 | 5 | 1 |

#### Communication & Collaboration

| Outcome # | Outcome | % Support | Mean | Max | Min |
|---|---|---|---|---|---|
| **1** | COMMUNICATION Share ideas, data, and findings with others clearly and accurately. | 97.8 | 4.8 | 5 | 1 |
| **1.1** | Use appropriate language and style to communicate science effectively to targeted audiences (e.g., general public, biology experts, collaborators in other disciplines). | 96.1 | 4.5 | 5 | 3 |
| **1.2** | Use a variety of modes to communicate science (e.g., oral, written, visual). | 97 | 4.6 | 5 | 3 |
| **2** | COLLABORATION Work productively in teams with people who have diverse backgrounds, skill sets, and perspectives. | 97 | 4.6 | 5 | 1 |

| | | | | | |
|---|---|---|---|---|---|
| **2.1** | Work with teammates to establish and periodically update group plans and expectations (e.g., team goals, project timeline, rules for group interactions, individual and collaborative tasks). | 84.8 | 4.2 | 5 | 1 |
| **2.2** | Elicit, listen to, and incorporate ideas from teammates with different perspectives and backgrounds. | 94.4 | 4.5 | 5 | 1 |
| **2.3** | Work effectively with teammates to complete projects. | 97.8 | 4.6 | 5 | 1 |
| **3** | COLLEGIAL REVIEW Provide and respond to constructive feedback in order to improve individual and team work. | 93.5 | 4.3 | 5 | 1 |
| **3.1** | Evaluate feedback from others and revise work or behavior appropriately. | 94.3 | 4.4 | 5 | 1 |
| **3.2** | Critique others' work and ideas constructively and respectfully. | 94.8 | 4.4 | 5 | 1 |
| **4** | METACOGNITION Reflect on your own learning, performance, and achievements. | 92.2 | 4.5 | 5 | 1 |
| **4.1** | Evaluate your own understanding and skill level. | 93.5 | 4.4 | 5 | 1 |
| **4.2** | Assess personal progress and contributions to your team and generate a plan to change your behavior as needed. | 89.5 | 4.3 | 5 | 1 |

**Science & Society**

| **Outcome #** | **Outcome** | **% Support** | **Mean** | **Max** | **Min** |
|---|---|---|---|---|---|
| **1** | ETHICS Demonstrate the ability to critically analyze ethical issues in the conduct of science. | 92.3 | 4.5 | 5 | 1 |
| **1.1** | Identify and evaluate ethical considerations (e.g., use of animal or human subjects, conflicts of interest, confirmation bias) in a given research study. | 90.8 | 4.3 | 5 | 1 |
| **1.2** | Critique how ethical controversies in biological research have been and can continue to be addressed by the scientific community. | 87 | 4.2 | 5 | 1 |
| **2** | SOCIETAL INFLUENCES Consider the potential impacts of outside influences (historical, cultural, political, technological) on how science is practiced. | 82.7 | 4.2 | 5 | 1 |
| **2.1** | Describe examples of how scientists' backgrounds and biases can influence science and how science is enhanced through diversity. | 83.1 | 4.2 | 5 | 1 |
| **2.2** | Identify and describe how systemic factors (e.g., socioeconomic, political) affect how and by whom science is conducted. | 78.9 | 4.1 | 5 | 1 |
| **3** | SCIENCE'S IMPACT ON SOCIETY Apply scientific reasoning in daily life and recognize the impacts of science on a local and global scale. | 96.6 | 4.7 | 5 | 1 |
| **3.1** | Apply evidence-based reasoning and biological knowledge in daily life (e.g., consuming popular media, deciding how to vote). | 95.8 | 4.7 | 5 | 1 |
| **3.2** | Use examples to describe the relevance of science in everyday experiences. | 94.1 | 4.5 | 5 | 1 |
| **3.3** | Identify and describe the broader societal impacts of biological research on different stakeholders. | 89 | 4.3 | 5 | 1 |
| **3.4** | Describe the roles scientists have in facilitating public understanding of science. | 86.9 | 4.2 | 5 | 1 |

**Supplemental Table 8. Skills that validation survey respondents suggested were missing from the BioSkills Guide.**

[a] Skills are summarized from comments in national validation survey. Annotations indicate that multiple people suggested adding that skill (i.e. "x2" indicates two respondents). In most cases, suggested skills are more specific or more challenging versions of existing outcomes in the BioSkills Guide.

| Core Competency | Missing Essential Skill [a] |
|---|---|
| Process of Science | Distinguish between the terms "hypothesis", "theory", and "fact". |
| | Describe how paradigms shift in biology. |
| | Describe the differences between various types of scientific literature. |
| | Identify assumptions and biases in scientific arguments. |
| | Use observation in the process of science and explain its importance. (x4) |
| | Entrepreneurialism. |
| Quantitative Reasoning | Do simple calculations without the use of a calculator. |
| | Use logic in the process of science (i.e. planning and implementing studies, writing, forming arguments). |
| | Consider hypotheses when designing and interpreting statistical tests. |
| | Identify the limitations of quantitative results. |
| | Use computational tools to analyze large datasets. (x2) |
| | Be familiar with and be able to self-teach a variety of scientific software programs. (x2) |
| Modeling | Define what a model is and the different ways we model biological phenomenon. |
| | Identify alternative assumptions that could be included in a model. |
| | Consider model biases. |
| | Construct simple quantitative models based on data. |
| | Project the implications of alternative model assumptions. |
| | Identify the objectives of a model before beginning construction. |
| | Build models that integrate multiple processes (e.g. positive and negative feedback loops). |
| Interdisciplinary Nature of Science | Describe the limits of science in addressing political or ethical issues. (x2) |
| | Consider differences in epistemology during interdisciplinary conversations. |
| Communication & Collaboration | Use a variety of modes to communicate science to a wide audience, including individuals with disabilities. |
| | Communicate science accurately and with sound logic. |
| | Write about science in plain language. |
| | Listen to and consider opposing views while maintaining respect and civility. |
| | Work productively in teams with people who have different abilities. |
| | Identify barriers to collaboration. |
| | Develop study habits that work well with your learning style. |
| | Evaluate use of prior knowledge. |
| Science & Society | Act ethically in academics and other work settings (cheating, plagiarism). (x4) |
| | Describe the strengths and limitations of the peer review process. |
| | Describe and critique the treatment of minority groups in biological and medical research. (x2) |
| | Explain why biology cannot be used to define race. |
| | Reflect on and describe the perceptions of science by the general public. (x2) |

**Supplemental Table 9. Sensitivity analysis comparing learning outcome support for respondents retained or excluded in RQ2 analysis.**

[a] All models contained respondent and learning outcome as random effects. Additional fixed effects added to models are indicated in left column. Likelihood Ratio (LR) tests were used to determine whether a model including learning outcomes and respondents as random effects has a better fit than a model that does not include these random effects. LR tests revealed that models with random effects were a better fit in all cases.

| Model [a] | AIC |
|---|---|
| **Random effects only (SA0)** | 7652.64 |
| **Exclusion indicator only (SA1)** | 7651.33 |
| **Competency only (SA2)** | 7629.10 |
| **Competency + Exclusion indicator (SA3)** | 7627.81 |
| **Competency X Exclusion indicator interaction (SA4)** | 7628.82 |

**Supplemental Table 10. RQ2 descriptive statistics: Distribution of respondent characteristics.**

[a] Total n for RQ2 was 346 respondents. Details of data processing are in Methods and Supplemental Methods.

| Demographic | Response | n [a] (%) |
|---|---|---|
| Institution Type | Associate's Granting | 82 (23.7%) |
| | Bachelor's Granting | 95 (27.5%) |
| | Master's Granting | 50 (14.4%) |
| | Doctoral Granting | 119 (34.4%) |
| Discipline-Based Education Research Experience | No DBER Experience | 232 (67.0%) |
| | DBER Experience | 114 (33.0%) |
| Currently Engaged in Biology Research | Not Currently Engaged in Biology Research | 167 (48.3%) |
| | Current Research in Biology | 179 (51.7%) |
| Experience with Ecology/Evolution Research | No Ecology/Evolution Experience | 218 (63.0%) |
| | Ecology/Evolution Experience | 128 (37.0%) |
| Familiarity with Vision and Change | Low Familiarity | 102 (29.5%) |
| | High Familiarity | 244 (70.5%) |

**Supplemental Table 11. RQ2 descriptive statistics: Number of learning outcomes per competency.**

[a] 77 total outcomes.

| | | n[a] (%) |
|---|---|---|
| Competency | Process of Science | 23 (29.9%) |
| | Quantitative Reasoning | 13 (16.9%) |
| | Modeling | 10 (13.0%) |
| | Interdisciplinary Nature of Science | 7 (9.1%) |
| | Communication & Collaboration | 13 (16.9%) |
| | Science & Society | 11 (14.3%) |

**Supplemental Table 12. Details of cross-classified multilevel binary logistic regression models of competency and respondent demographics predicting learning outcome support.**

[a] Best fitting models for each research question are shown. Both models contained respondent and learning outcome as random effects. Additional fixed effects in models are indicated in top row. For RQ2a, the initial complex model used 'Support' as the dependent variable and included a random effect for learning outcome, a random effect for respondent, and a fixed effect for learning outcome competency. For RQ2b, the initial complex model used 'Support' as the dependent variable and included a random effect for learning outcome, a random effect for respondent, and five interactions as fixed effects: competency X institution type, competency X experience in DBER, competency X engagement in disciplinary biology research, competency X experience in ecology/evolution, and competency X Vision and Change familiarity.

[b] OR = odds ratio, SE = standard error. *** p < 0.001; ** p < 0.01; * p <0.05.

[c] ref = reference category. Quantitative reasoning was used as Competency reference category because it was the most highly rated overall.

[d] For categorical variables with three or more categories (i.e. Competency, Institution Type), the Wald Chi-squared Test evaluates the joint significance of all coefficients for that variable (e.g., tests whether or not all coefficients related to Competency are significant overall). Furthermore, interpreting individual coefficients is not recommended for logistic regression models (Long & Freese, 2014; Mustillo, Lizardo, & McVeigh, 2018). Instead, for interpretation, see predicted probabilities (Figure 4) and Wald Chi-squared Test.

[e] AIC = Akaike's Information Criterion. AIC for model with random effects only = 6412.06.

[f] A significant Likelihood Ratio test indicates that a model including learning outcomes and respondents as random effects is a better fit than a model that does not include these random effects. LR test for model with only random effects = 1856.24***. LR tests revealed that models with random effects were a better fit in all cases.

| **Model [a]** | | **Main Effect: Competency (RQ2a)** | | **Interactions with Competency: Institution Type, Discipline-Based Education Research, Biology Research; Main Effect: Eco/Evo Experience (RQ2b)** | |
|---|---|---|---|---|---|
| | | **OR [b]** | ***SE* [b]** | **OR** | ***SE*** |
| **Competency, ref [c]= Quantitative Reasoning** | Process of Science | 0.672 | *0.235* | 1.003 | *0.494* |
| | Modeling | 0.162 *** | *0.066* | 0.197** | *0.109* |
| | Interdisciplinary Nature of Science | 0.153 *** | *0.069* | 0.215** | *0.124* |
| | Communication & Collaboration | 0.585 | *0.227* | 1.086 | *0.576* |
| | Science & Society | 0.254 *** | *0.102* | 0.202** | *0.109* |
| **Institution Type, ref = Doctoral Granting** | Associate's Granting | | | 2.079 | *0.995* |
| | Bachelor's Granting | | | 0.634 | *0.260* |
| | Master's Granting | | | 0.760 | *0.401* |
| **Competency X Institution Type, ref = Quantitative Reasoning X Doctoral Granting** | Process of Sci. X Associate's Granting | | | 0.455 | *0.209* |
| | Process of Sci. X Bachelor's Granting | | | 1.521 | *0.559* |
| | Process of Sci. X Master's Granting | | | 3.093* | *1.587* |
| | Modeling X Associate's Granting | | | 0.532 | *0.256* |
| | Modeling X Bachelor's Granting | | | 0.819 | *0.310* |
| | Modeling X Master's Granting | | | 0.562 | *0.277* |
| | Interdisc. Nature of Sci. X Associate's Granting | | | 0.429 | *0.203* |
| | Interdisc. Nature of Sci. X Bachelor's Granting | | | 1.661 | *0.652* |
| | Interdisc. Nature of Sci. X Master's Granting | | | 0.976 | *0.482* |
| | Communic. & Collabor. X Associate's Granting | | | 0.633 | *0.306* |

| | | | | | |
|---|---|---|---|---|---|
| | Communic. & Collabor. X Bachelor's Granting | | | 1.286 | *0.481* |
| | Communic. & Collabor. X Master's Granting | | | 2.745 | *1.474* |
| | Sci. & Society X Associate's Granting | | | 0.815 | *0.372* |
| | Sci. & Society X Bachelor's Granting | | | 1.694 | *0.636* |
| | Sci. & Society X Master's Granting | | | 1.388 | *0.692* |
| **Discipline-Based Education Research, ref = No Experience** | Experience in Discipline-Based Education Research | | | 0.750 | *0.274* |
| **Competency X Discipline-Based Education Research, ref = Quantitative Reasoning X No Experience** | Process of Sci. X Experience | | | 0.983 | *0.332* |
| | Modeling X Experience | | | 4.021*** | *1.478* |
| | Interdisc. Nature of Sci. X Experience | | | 1.139 | *0.408* |
| | Communication & Collaboration X Experience | | | 0.823 | *0.285* |
| | Sci. & Society X Experience | | | 1.809 | *0.621* |
| **Biology Research, ref = Not Currently Engaged** | Currently Engaged in Biology Research | | | 2.226* | *0.880* |
| **Competency X Biology Research, ref = Quantitative Reasoning X Not Currently Engaged** | Process of Sci. X Currently Engaged | | | 0.375** | *0.138* |
| | Modeling X Currently Engaged | | | 0.505 | *0.201* |
| | Interdisc. Nature of Sci. X Currently Engaged | | | 0.479 | *0.181* |
| | Communic. & Collabor. X Currently Engaged | | | 0.287** | *0.112* |
| | Sci. & Society X Currently Engaged | | | 0.690 | *0.265* |
| **Ecology/ Evolution, ref = No Experience** | Eco/Evo Experience | | | 1.632* | *0.384* |
| **Constant** | | 105.201*** | *32.446* | 68.839*** | *33.421* |
| **Wald Chi-squared Test** [d] | | *Competency, $\chi^2$=37.82**** | | *Competency x Institution Type, $\chi^2$=35.76***<br>*Competency x Discipline-Based Education Research, $\chi^2$=27.06****<br>*Competency x Biology Research, $\chi^2$=13.89** | |
| **ΔAIC (relative to model with only random effects)** [e] | | -22.21 | | -52.18 | |
| **Likelihood Ratio Test** [f] | | 1678.50*** | | 1648.14*** | |

#### Supplemental Material 5. Supplemental Methods.

##### Workshop and Round Table Design and Implementation

We employed 5 workshops and 2 round tables over the course of the development phase (Table 2). Workshops were held at biology education learning community meetings at universities around the Northwest United States and British Columbia, unless specified otherwise. During workshops, we instructed participants to self-select into six smaller groups based on which competency they wished to focus on. We then directed groups to brainstorm outcomes essential to their competency, with the goal of eliciting a range of perspectives and ideas. Next we provided handouts of the current draft of learning outcomes for their competency and asked groups to discuss and record: (1) whether each outcome was important for a graduating general biology major (yes/no/maybe), (2) any essential outcomes they felt were missing, and (3) any comments on the wording or content of the draft.

Round tables were held at national biology education research meetings and used to collect targeted feedback on parts of the BioSkills Guide for which it was more difficult to find appropriate revisions. During round tables, we also instructed participants to talk in groups, but the topic of discussion and specific instructions varied depending on the issue at hand (elaborated in detailed descriptions below). We transcribed and summarized all written feedback from workshops and round tables for use during revision sessions.

##### Interview Design and Implementation

We conducted 25 interviews over the course of the development phase (Table 2). Interviews were either semi-structured or unstructured depending on their purpose (elaborated in detailed descriptions below). We conducted interviews in a variety of settings depending on participant availability (in person, video chat, or over the phone). When possible, the interviews were recorded, otherwise the interviewer took notes during the interview and expanded upon them immediately after the interview. Detailed notes and recordings (when available) from interviews were analyzed to identify major themes that then informed revisions.

##### Detailed Description of the Initial Drafting of the BioSkills Guide

The initial draft of the BioSkills Guide was composed by a group of 8 biology faculty members at a large, public research university in the Northwest United States. This work was initiated at an all-faculty departmental meeting where the Vision and Change core competencies were presented and discussed, along with several other guiding documents related to science competencies (e.g., AAMC & HHMI, 2009; NRC, 2012). Four department competency priorities were selected and broadly defined: Process of Science, Quantitative Reasoning (which included some Modeling), Communication & Collaboration, and Science & Society. Interdisciplinary Nature of Science was understood to run through all four competencies. Four working groups of two faculty each then drafted learning outcomes for one of the four competencies. Drafts were then shared with members of other working groups and 12 additional interested faculty, postdocs, and grad students at a series of departmental round table meetings (n=20 participants total, 5-12 per competency, Table 2). Participants suggested additions and changes, which the working groups used to revise the drafts.

We built on this initial draft by aligning it with Vision and Change and broader work in biology education research. We began by drafting learning outcomes for Interdisciplinary Nature of Science, disentangling Modeling outcomes from Quantitative Reasoning outcomes, and checking for gaps in coverage within the remaining competencies. We accomplished these tasks by reviewing the literature, leading unstructured interviews with competency experts (n=11), and hosting a round table at a national biology education research meeting (n≈24; see note about estimation in Table 2). At the round table, we asked participants to discuss and record suggestions for priorities and appropriate challenge level for the Interdisciplinary Nature of Science competency. We recruited competency experts for interviews based on their history of publication or conference presentations in areas related to underdeveloped portions of the guide (i.e. Modeling, Interdisciplinary Nature of Science, and Science and Society). Finally, we revised learning outcomes for all six competencies to a common tone and formatting. The initial draft ("Version I") contained 86 outcomes: 23 "program-level" and 63 "course-level" (see Supplemental Figure 1 for overview of guide structure).

**Detailed Description of the Process of Iterative Review and Revision of the BioSkills Guide**

After initial drafting was complete, Version I learning outcomes were reviewed by our project advisory board, who then provided written feedback. We then clarified feedback via a virtual meeting. Two authors (AWC, AJC) discussed all feedback and collectively decided to add one outcome, remove one outcome, and revise 30 outcomes (Supplemental Table 1).

To assess Version II of the BioSkills Guide, we led the first of five workshops (n≈30; see note about estimation in Table 2). Participants were primarily biology instructors and postdocs, but also included some graduate students and undergraduates. Feedback from the workshop was discussed by two authors (AWC, AJC). We decided to remove seven outcomes, add two outcomes, and revise 57 outcomes (Supplemental Table 1). Additionally, workshop participants raised concerns about the order of outcomes within the guide, leading to some rearrangements of outcomes within competencies.

Version III was reviewed via web survey (n=21 total, n=6-10 per competency) and a small workshop (n=6 total, n=2 per competency) (Table 2). Of the 81 outcomes in Version III, we removed six, added three, and revised 57 to generate Version IV (Supplemental Table 1).

Version IV was reviewed via web survey (n=45 total, n=12-19 per competency), workshop (n=32 total, n=4-6 per competency), a round table (n=21), and interviews (n=14) (Table 2). The workshop was held at a regional meeting of biology community college instructors (approximately 66% of participants were from community colleges). During the round table, we asked participants to split into groups focusing on one of four areas of low consensus (based on mixed survey ratings and comments: Modeling, Interdisciplinary Nature of Science, a Process of Science outcome on "doing research", and attitude-/affect-related outcomes) and instructed them to discuss and record ideas for revision.

We conducted interviews for different purposes with three different populations using an opportunistic sampling approach: competency experts (n=6), survey respondents (n=5), and community college faculty (n=3). We recruited competency experts to provide guidance on revising outcomes for less frequently taught or understood competencies (e.g., Modeling, Interdisciplinary Nature of Science), where survey ratings were low or mixed, but comments did not suggest specific revisions. These interviews were unstructured to allow competency experts to direct the conversation to what they felt was most essential about that competency and therefore must be retained during revision. Interviews with past survey respondents were semi-structured, with questions varying depending on which competencies they had reviewed. The purpose of these interviews was to gain additional insight on learning outcomes with low ratings and determine what about the outcomes should be revised (e.g., level of challenge, unclear terminology). Interviews with community college faculty were unstructured and involved asking participants to comment on the outcomes and identify points of connection (or lack thereof) between the BioSkills Guide and their own classroom practices. The purpose of these interviews was to identify areas of the guide that required revision to be valuable in a two-year setting. Feedback on Version IV prompted us to remove four outcomes, add six, and revise 64.

Version V was reviewed via web survey (n=27 total, n=8-14 per competency) and two workshops (n=21 and n=8 total, with n=2-5 per competency). As a result, we removed three outcomes, added none, and revised 29 to generate Version VI.

**Detailed Description of the Pilot Validation**
A smaller-scale pilot validation was conducted to test the new questionnaire (which had been shortened and reformatted for validation phase) and our final draft of outcomes before inviting a large number of educators nationwide to participate. We invited 45 biology educators from the local Partnership for Undergraduate Life Sciences Education (PULSE) network to review Version VI outcomes via web survey. We chose this population because they represented a range of institution types and were expected to have spent time thinking deeply about the undergraduate biology curriculum, since they had participated in a PULSE workshop which includes professional development on Vision and Change recommendations. Twenty people completed the pilot validation survey (n=11-12 per competency, 44% participation rate). Of the 77 outcomes in Version VI, 74 had greater than 80% support (Table 3, Supplemental Table 4). Of the three remaining outcomes, support ranged from 73%-75%. Two of the three were from the Modeling competency and had been strongly advocated for in interviews with experts during review of Version IV. The third was revised, as described in Methods.

**Missing Data and Sensitivity Analysis for Excluded Cases**
Of our 417 initial respondents who rated at least one outcome (and thus were included in RQ1b), 71 did not provide all five demographic characteristics of interest (i.e. institution type, familiarity with Vision and Change, etc.), and therefore could not be included in our analyses for RQ2a and b. Of these 71 individuals, the majority (n=48) left the web survey before viewing all pages with their assigned outcomes (i.e. breakoff cases) and therefore never saw the demographic questions. Of the individuals that *did* view all pages with their assigned outcomes, 17 did not respond to all demographic questions required for RQ2, and 2 did not respond to any demographic questions at all. We also removed 4 individuals who indicated “other” for institution type.

As a sensitivity analysis, we evaluated whether or not the odds of supporting an outcome (i.e. the dependent variable in our RQ2 analyses) differed across respondents who were excluded from the RQ2 analysis (n=71) and respondents who were included in the RQ2 analysis (n=346). In other words, were respondents who supported competency learning outcomes less more likely to leave the survey early or skip demographic questions? We explored this question using backward model selection beginning with complex cross-classified multilevel binary logistic regression models predicting whether particular respondents will support particular learning outcomes, as described further in Methods. All models contained learning outcome and respondent as random effects (random intercepts).

We first fit a model containing one fixed effect: a binary “exclusion indicator” variable for whether or not a respondent was excluded from the RQ2 analysis (=0 if respondent was included; =1 if not). Removing this variable from the model did not affect model fit relative to a model with only random effects (SA1-SA0, ΔAIC = -1.31, Supplemental Table 9). Thus, respondents that were *excluded* from the RQ2 analysis did not differ in their odds of supporting a learning outcome compared to respondents that were *included* in the RQ2 analysis.

We then examined whether or not the odds of supporting learning outcomes for different competencies differed across respondents that were *excluded* from the RQ2 analysis and respondents that were *included* in the RQ2 analysis. We carried out backward model selection starting with a complex model (SA4) including the interaction between outcome competency (e.g., Process of Science, Modeling) and the exclusion indicator from SA1. The best fitting and most parsimonious model was the one with only competency as a fixed effect (SA2). Neither removing the competency X exclusion indicator interaction (SA4) nor removing the inclusion of the exclusion indicator as a fixed effect (SA3) affected model fit relative to a model with just competency as a fixed effect (SA2) (SA3-SA4, ΔAIC = -1.01; SA2-SA3, ΔAIC = 1.29; Supplemental Table 9). Thus, respondents that were *excluded* from the RQ2 analysis again did not differ in their odds of supporting a learning outcome compared to respondents that were *included* in the RQ2 analysis, within each competency (i.e., exclusion indicator x competency interaction) nor when including competency only as a main effect (i.e., no exclusion indicator x competency interaction).

##### Data Recoding for Statistical Models

After data processing as described above and in Methods, we recoded variables as follows: Three respondents who indicated "other" for institution type (out of 7 total) were recoded based on Carnegie classification of institution name provided or description of institution in comments (e.g., "we are part of a larger R1, but our campus strictly grants Associate's degrees" was assigned to "Associate's Granting"). The remaining four respondents were removed from analysis as mentioned above. Vision and Change familiarity was recoded to a binary variable: 'Extremely' or 'Very Familiar' were recoded as 'High Familiarity' and 'Somewhat', 'Slightly', or 'Not at All Familiar' were recoded as 'Low Familiarity'. Experience in Discipline-Based Education Research and Experience in Ecology/Evolution Research were coded as binary variables based on selecting the corresponding field when answering questions about field of current research and/or field of graduate training. Current engagement in biology research was coded as a binary variable based on selecting a disciplinary biology field (e.g., "Molecular/Cell/Developmental Biology", "Physiology", "Ecology/Evolutionary Biology") when answering question about field of current research. Finally, importance ratings for each course-level and program-level learning outcome were recoded: 'Important' or 'Very Important' were recoded as 'Support', and 'Neither Important nor Unimportant', 'Unimportant', or 'Very Unimportant' were recoded as 'No Support'.

**Supplemental Material 6. BioSkills development phase questionnaire.**

Questionnaire used during development phase survey for review of Version V. Questionnaires for Versions III and IV were very similar, except for revision of learning outcomes.

### BioSkills: Core Competencies Learning Outcomes

**Welcome Page**

**Thank you for your help developing learning outcomes for the core competencies, or essential skills, for biology undergraduate education.** Based on the recommendations of the 2011 AAAS report *Vision and Change in Undergraduate Biology Education*, this NSF-supported project is intended to use the perspectives and priorities of a wide range of biology educators to elaborate six "core competencies" (listed below) into measurable learning outcomes. The learning outcomes will be revised based on your and others' feedback, and then be made available to the biology education community as a resource to facilitate planning and assessment of skills training. It is essential that this work is done collaboratively to ensure the final set of learning outcomes (which we're calling the "BioSkills Guide") has broad utility for biology educators teaching at different institution types, course levels, and biology subdisciplines. Thank you for being a part of this work!

You will be asked to provide feedback on learning outcomes for just two of the six core competencies, although we would love your feedback on additional competencies if you have time. You do **not** need to have experience teaching the core competencies as long as you have served as instructor of record for a college-level biology course. Your feedback is valuable, and we sincerely appreciate all comments and suggestions.

You may exit and return to the survey as needed. The survey automatically resumes where you left off if you use the same browser and do not clear cookies. If you have any questions about the project or this survey, please do not hesitate to contact me, Dr. Alexa Clemmons, or Dr. Alison Crowe.

**Vision and Change Core Competencies**

- **Process of Science**
- **Quantitative Reasoning**
- **Modeling & Simulations**
- **Interdisciplinary Nature of Science**
- **Communication & Collaboration**
- **Science & Society**

Have you ever served as the instructor of record for a college-level biology course?

- ○ Yes
- ○ No

**Screened Out**

Thank you for your interest in the BioSkills Guide! At this time we are soliciting feedback from college biology instructors, but in the future we plan to widen our scope.

If you would be interested in providing feedback then, please enter your email address. If you enter your email address we will also notify you when the BioSkills Guide is ready for distribution.

**Process of Science**

#### [***After the welcome page, blocks of questions (each block corresponding to one of the six core competencies) were randomly assigned. All respondents were given 2 blocks of questions, with the option to complete additional.***]

In this portion of the survey, we would like you to rate the importance and ease of understanding of our current draft of learning outcomes for one particular core competency: **Process of Science.** Later, you will be asked to comment on whether the outcomes are appropriately categorized under this competency.

In the current draft of the BioSkills Guide, each core competency contains multiple **program**-level learning outcomes. In addition, each program-level learning outcome contains multiple **course**-level learning outcomes. Throughout the survey, you will switch between evaluating program- and course-level outcomes. We will use the figure below to remind you of this structure and cue when you are switching between levels.

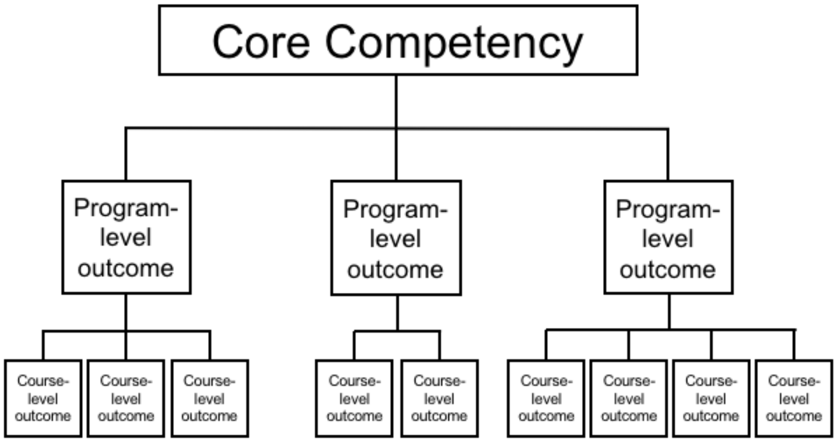


**Important note:** Please keep in mind that we intend for the BioSkills Guide to contain the learning outcomes that we, as a community, think all **graduating general biology majors** should achieve. Therefore, as you complete the survey, please rate the outcomes based on whether they are both desirable and reasonable to accomplish in a **four-year program**, not introductory courses alone (1-2 years only) or in a graduate program (5+ years).

Importance and Ease of Understanding: **Process of Science**

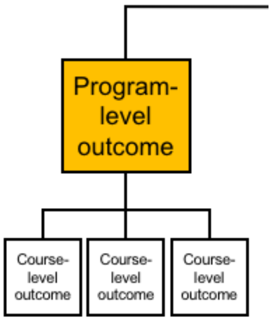


The next three questions will ask you to evaluate the following **program**-level outcome:

**Scientific Thinking: Explain how science generates knowledge of the natural world.**

How important or unimportant is it for graduating general biology majors to achieve this outcome?

- Very Important
- Important
- Neither Important Nor Unimportant
- Unimportant
- Very Unimportant

How easy or difficult is it for **you** to understand this outcome?

- Very Easy
- Easy
- Neither Easy Nor Difficult
- Difficult
- Very Difficult

Please share any feedback you have about the content or wording of this outcome.

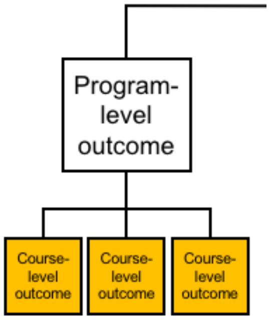


The next three questions will ask you to evaluate the following **course**-level outcome:

**Explain how scientists use inference, a skeptical mindset, and evidence-based reasoning to generate knowledge.**

How important or unimportant is it for graduating general biology majors to achieve this outcome?

- Very Important
- Important
- Neither Important Nor Unimportant
- Unimportant
- Very Unimportant

How easy or difficult is it for **you** to understand this outcome?

- Very Easy
- Easy

- Neither Easy Nor Difficult
- Difficult
- Very Difficult

Please share any feedback you have about the content or wording of this outcome.

**Course**-level outcome:

**Describe the iterative nature of science and how new evidence can lead to the revision of scientific knowledge.**

How important or unimportant is it for graduating general biology majors to achieve this outcome?

- Very Important
- Important
- Neither Important Nor Unimportant
- Unimportant
- Very Unimportant

How easy or difficult is it for **you** to understand this outcome?

- Very Easy
- Easy
- Neither Easy Nor Difficult
- Difficult
- Very Difficult

Please share any feedback you have about the content or wording of this outcome.

#### Importance and Ease of Understanding: **Process of Science**

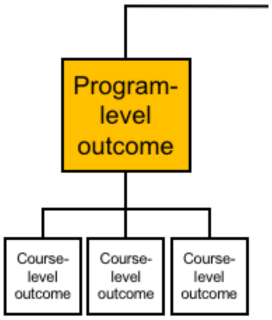


The next three questions will ask you to evaluate the following **program**-level outcome:

**Information Literacy: Locate, interpret, and evaluate scientific information.**

How important or unimportant is it for graduating general biology majors to achieve this outcome?

- Very Important
- Important
- Neither Important Nor Unimportant
- Unimportant
- Very Unimportant

How easy or difficult is it for **you** to understand this outcome?

- Very Easy
- Easy
- Neither Easy Nor Difficult
- Difficult
- Very Difficult

Please share any feedback you have about the content or wording of this outcome.

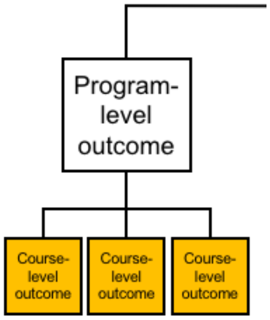


The next three questions will ask you to evaluate the following **course**-level outcome:

**Find and evaluate credibility of a variety of sources of scientific information, including popular science media and scientific journals.**

How important or unimportant is it for graduating general biology majors to achieve this outcome?

- Very Important
- Important
- Neither Important Nor Unimportant
- Unimportant
- Very Unimportant

How easy or difficult is it for **you** to understand this outcome?

- Very Easy
- Easy
- Neither Easy Nor Difficult
- Difficult
- Very Difficult

Please share any feedback you have about the content or wording of this outcome.

#### **Course**-level outcome:

**Interpret, evaluate, and summarize evidence in primary literature.**

How important or unimportant is it for graduating general biology majors to achieve this outcome?

- Very Important
- Important
- Neither Important Nor Unimportant
- Unimportant
- Very Unimportant

How easy or difficult is it for **you** to understand this outcome?

- Very Easy
- Easy
- Neither Easy Nor Difficult
- Difficult
- Very Difficult

Please share any feedback you have about the content or wording of this outcome.

**Course**-level outcome:

**Evaluate claims in scientific papers, popular science media, and other sources using evidence-based reasoning.**

How important or unimportant is it for graduating general biology majors to achieve this outcome?

- Very Important
- Important
- Neither Important Nor Unimportant
- Unimportant
- Very Unimportant

How easy or difficult is it for **you** to understand this outcome?

- Very Easy
- Easy
- Neither Easy Nor Difficult
- Difficult
- Very Difficult

Please share any feedback you have about the content or wording of this outcome.

Importance and Ease of Understanding: **Process of Science**

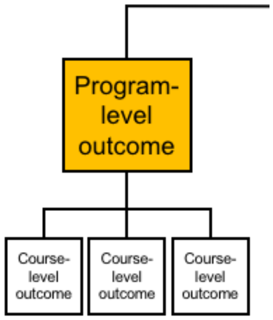


The next three questions will ask you to evaluate the following **program**-level outcome:

**Question Formulation: Pose testable questions and hypotheses to address gaps in knowledge.**

How important or unimportant is it for graduating general biology majors to achieve this outcome?

- Very Important
- Important
- Neither Important Nor Unimportant
- Unimportant
- Very Unimportant

How easy or difficult is it for **you** to understand this outcome?

- Very Easy
- Easy
- Neither Easy Nor Difficult
- Difficult
- Very Difficult

Please share any feedback you have about the content or wording of this outcome.

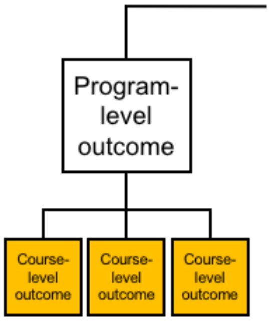


The next three questions will ask you to evaluate the following **course**-level outcome:

**Identify gaps in current understanding of a biological system or process and articulate what specific information is missing.**

How important or unimportant is it for graduating general biology majors to achieve this outcome?

- Very Important
- Important
- Neither Important Nor Unimportant
- Unimportant
- Very Unimportant

How easy or difficult is it for **you** to understand this outcome?

- Very Easy
- Easy
- Neither Easy Nor Difficult
- Difficult
- Very Difficult

Please share any feedback you have about the content or wording of this outcome.

##### **Course**-level outcome:

**Develop questions based on your own or others' observations.**

How important or unimportant is it for graduating general biology majors to achieve this outcome?

- Very Important
- Important
- Neither Important Nor Unimportant
- Unimportant
- Very Unimportant

How easy or difficult is it for **you** to understand this outcome?

- Very Easy
- Easy
- Neither Easy Nor Difficult
- Difficult
- Very Difficult

Please share any feedback you have about the content or wording of this outcome.

##### **Course**-level outcome:

**Formulate testable hypotheses and state their predictions.**

How important or unimportant is it for graduating general biology majors to achieve this outcome?

- Very Important
- Important
- Neither Important Nor Unimportant
- Unimportant
- Very Unimportant

How easy or difficult is it for **you** to understand this outcome?

- Very Easy
- Easy
- Neither Easy Nor Difficult
- Difficult
- Very Difficult

Please share any feedback you have about the content or wording of this outcome.

Importance and Ease of Understanding: **Process of Science**

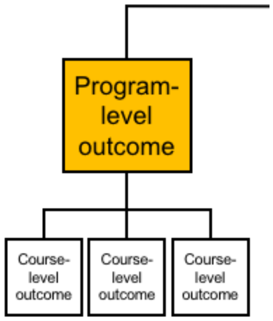


The next three questions will ask you to evaluate the following **program**-level outcome:

**Study Design: Plan, evaluate, and implement scientific investigations.**

How important or unimportant is it for graduating general biology majors to achieve this outcome?

- Very Important
- Important
- Neither Important Nor Unimportant
- Unimportant
- Very Unimportant

How easy or difficult is it for **you** to understand this outcome?

- Very Easy
- Easy
- Neither Easy Nor Difficult
- Difficult
- Very Difficult

Please share any feedback you have about the content or wording of this outcome.

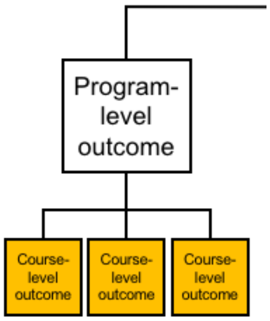


The next three questions will ask you to evaluate the following **course**-level outcome:

**Compare the strengths and limitations of various study designs.**

How important or unimportant is it for graduating general biology majors to achieve this outcome?

- Very Important
- Important
- Neither Important Nor Unimportant
- Unimportant
- Very Unimportant

How easy or difficult is it for **you** to understand this outcome?

- Very Easy
- Easy
- Neither Easy Nor Difficult
- Difficult
- Very Difficult

Please share any feedback you have about the content or wording of this outcome.

**Course**-level outcome:

**Design controlled experiments, including appropriate data analysis plans.**

How important or unimportant is it for graduating general biology majors to achieve this outcome?

- Very Important
- Important
- Neither Important Nor Unimportant
- Unimportant
- Very Unimportant

How easy or difficult is it for **you** to understand this outcome?

- Very Easy
- Easy
- Neither Easy Nor Difficult
- Difficult
- Very Difficult

Please share any feedback you have about the content or wording of this outcome.

**Course**-level outcome:

**Execute protocols and accurately record measurements and observations.**

How important or unimportant is it for graduating general biology majors to achieve this outcome?

- Very Important
- Important
- Neither Important Nor Unimportant
- Unimportant
- Very Unimportant

How easy or difficult is it for **you** to understand this outcome?

- Very Easy
- Easy
- Neither Easy Nor Difficult
- Difficult
- Very Difficult

Please share any feedback you have about the content or wording of this outcome.

**Course**-level outcome:

**Identify methodological problems and suggest solutions or alternative approaches.**

How important or unimportant is it for graduating general biology majors to achieve this outcome?

- Very Important
- Important
- Neither Important Nor Unimportant
- Unimportant
- Very Unimportant

How easy or difficult is it for **you** to understand this outcome?

- Very Easy
- Easy
- Neither Easy Nor Difficult
- Difficult
- Very Difficult

Please share any feedback you have about the content or wording of this outcome.

**Course**-level outcome:

**Evaluate and suggest best practices for responsible research conduct (e.g., data management, lab safety, proper citation of sources).**

How important or unimportant is it for graduating general biology majors to achieve this outcome?

- Very Important
- Important
- Neither Important Nor Unimportant
- Unimportant
- Very Unimportant

How easy or difficult is it for **you** to understand this outcome?

- Very Easy
- Easy
- Neither Easy Nor Difficult
- Difficult
- Very Difficult

Please share any feedback you have about the content or wording of this outcome.

Importance and Ease of Understanding: **Process of Science**

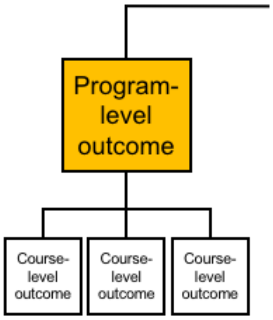


The next three questions will ask you to evaluate the following **program**-level outcome:

**Data Interpretation & Evaluation: Interpret, evaluate, and draw conclusions from data in order to make evidence-based arguments about the natural world.**

How important or unimportant is it for graduating general biology majors to achieve this outcome?

- Very Important
- Important
- Neither Important Nor Unimportant
- Unimportant
- Very Unimportant

How easy or difficult is it for **you** to understand this outcome?

- Very Easy
- Easy
- Neither Easy Nor Difficult
- Difficult
- Very Difficult

Please share any feedback you have about the content or wording of this outcome.

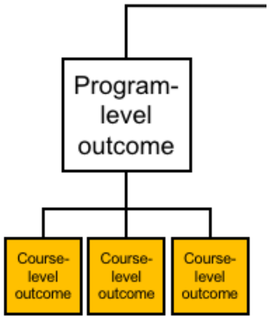


The next three questions will ask you to evaluate the following **course**-level outcome:

**Analyze data and summarize resulting patterns.**

How important or unimportant is it for graduating general biology majors to achieve this outcome?

- Very Important
- Important
- Neither Important Nor Unimportant
- Unimportant
- Very Unimportant

How easy or difficult is it for **you** to understand this outcome?

- Very Easy
- Easy
- Neither Easy Nor Difficult
- Difficult
- Very Difficult

Please share any feedback you have about the content or wording of this outcome.

**Course**-level outcome:

**Describe sources of error and uncertainty in data.**

How important or unimportant is it for graduating general biology majors to achieve this outcome?

- Very Important
- Important
- Neither Important Nor Unimportant
- Unimportant
- Very Unimportant

How easy or difficult is it for **you** to understand this outcome?

- Very Easy
- Easy
- Neither Easy Nor Difficult
- Difficult
- Very Difficult

Please share any feedback you have about the content or wording of this outcome.

#### **Course**-level outcome:

**Engage in data-driven argumentation using your own and others' findings.**

How important or unimportant is it for graduating general biology majors to achieve this outcome?

- Very Important
- Important
- Neither Important Nor Unimportant
- Unimportant
- Very Unimportant

How easy or difficult is it for **you** to understand this outcome?

- Very Easy
- Easy
- Neither Easy Nor Difficult
- Difficult
- Very Difficult

Please share any feedback you have about the content or wording of this outcome.

**Course**-level outcome:

**Relate conclusions to original hypothesis and suggest future research directions based on findings.**

How important or unimportant is it for graduating general biology majors to achieve this outcome?

- Very Important
- Important
- Neither Important Nor Unimportant
- Unimportant
- Very Unimportant

How easy or difficult is it for **you** to understand this outcome?

- Very Easy
- Easy
- Neither Easy Nor Difficult
- Difficult
- Very Difficult

Please share any feedback you have about the content or wording of this outcome.

Importance and Ease of Understanding: **Process of Science**

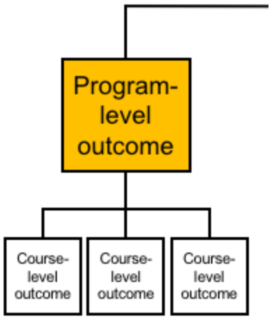


The next three questions will ask you to evaluate the following **program**-level outcome:

**Doing Research: Integrate process of science skills to address a research question in a course-based or independent research experience.**

How important or unimportant is it for graduating general biology majors to achieve this outcome?

- Very Important

- Important
- Neither Important Nor Unimportant
- Unimportant
- Very Unimportant

How easy or difficult is it for **you** to understand this outcome?

- Very Easy
- Easy
- Neither Easy Nor Difficult
- Difficult
- Very Difficult

Please share any feedback you have about the content or wording of this outcome.

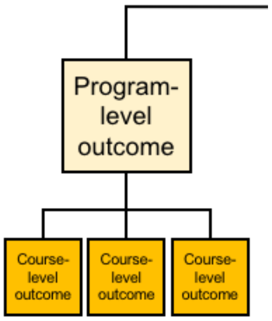


Next, we would like you to indicate whether the current categorization of **course**-level outcomes within program-level outcomes makes sense to you. You will also be asked to suggest missing outcomes.

Some related outcomes may currently be categorized under other competencies. If you would like to see a complete draft of the BioSkills Guide for context, please click here. This is a preliminary draft, and we ask that you *please do not share it with others until it is published.*

#### Categorization: **Process of Science**

Currently, we have categorized each of the course-level outcomes listed below under the program-level outcome:

**Scientific Thinking: Explain how science generates knowledge of the natural world.**

In your opinion, is it accurate to categorize each of the following course-level outcomes under this program-level outcome?

| | Yes | No |
|---|---|---|

| | Yes | No |
|---|---|---|
| Explain how scientists use inference, a skeptical mindset, and evidence-based reasoning to generate knowledge. | ○ | ○ |
| Describe the iterative nature of science and how new evidence can lead to the revision of scientific knowledge. | ○ | ○ |

Program-level outcome:

**Information Literacy: Locate, interpret, and evaluate scientific information.**

In your opinion, is it accurate to categorize each of the following course-level outcomes under this program-level outcome?

| | Yes | No |
|---|---|---|
| Find and evaluate credibility of a variety of sources of scientific information, including popular science media and scientific journals. | ○ | ○ |
| Interpret, evaluate, and summarize evidence in primary literature. | ○ | ○ |
| Evaluate claims in scientific papers, popular science media, and other sources using evidence-based reasoning. | ○ | ○ |

Program-level outcome:

**Question Formulation: Pose testable questions and hypotheses to address gaps in knowledge.**

In your opinion, is it accurate to categorize each of the following course-level outcomes under this program-level outcome?

| | Yes | No |
|---|---|---|
| Identify gaps in current understanding of a biological system or process and articulate what specific information is missing. | ○ | ○ |
| Develop questions based on your own or others' observations. | ○ | ○ |
| Formulate testable hypotheses and state their predictions. | ○ | ○ |

Optional: Please share any comments you have on the categorization of these outcomes, including any suggestions for alternative categorization if applicable.

Do you think any essential **course**-level outcomes are missing from this list?

Categorization: **Process of Science**

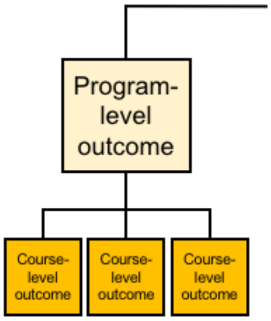


Currently, we have categorized each of the course-level outcomes listed below under the program-level outcome:

**Study Design: Plan, evaluate, and implement scientific investigations.**

In your opinion, is it accurate to categorize each of the following course-level outcomes under this program-level outcome?

| | Yes | No |
|---|---|---|
| Compare the strengths and limitations of various study designs. | O | O |
| Design controlled experiments, including appropriate data analysis plans. | O | O |
| Execute protocols and accurately record measurements and observations. | O | O |
| Identify methodological problems and suggest solutions or alternative approaches. | O | O |
| Evaluate and suggest best practices for responsible research conduct (e.g., data management, lab safety, proper citation of sources). | O | O |

Currently, we have categorized each of the course-level outcomes listed below under the program-level outcome:

**Data Interpretation & Evaluation: Interpret, evaluate, and draw conclusions from data in order to make evidence-based arguments about the natural world.**

In your opinion, is it accurate to categorize each of the following course-level outcomes under this program-level outcome?

| | Yes | No |
|---|---|---|
| Analyze data and summarize resulting patterns. | O | O |

| | Yes | No |
|---|---|---|
| Describe sources of error and uncertainty in data. | ○ | ○ |
| Engage in data-driven argumentation using your own and others' findings. | ○ | ○ |
| Relate conclusions to original hypothesis and suggest future research directions based on findings. | ○ | ○ |

Optional: Please share any comments you have on the categorization of these outcomes, including any suggestions for alternative categorization if applicable.

Do you think any essential **course**-level outcomes are missing from this list?

Categorization: **Process of Science**

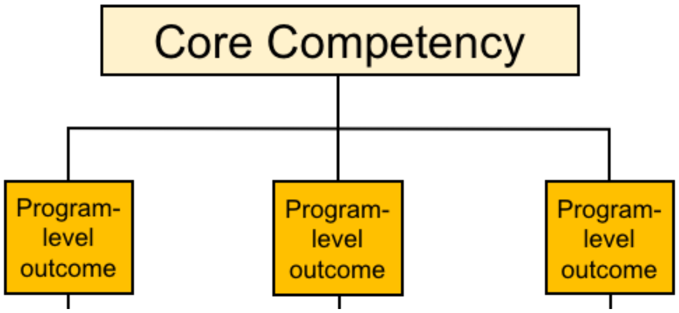


Next, we would like you to indicate if you think any of the **program**-level learning outcomes are currently miscategorized.

If you would like to see the complete, preliminary draft of the BioSkills Guide for context, please click here. As a reminder, the other core competencies are: **Quantitative Reasoning, Modeling & Simulations, Interdisciplinary Nature of Science, Communication & Collaboration, and Science & Society.**

Currently, we have categorized the program-level outcomes listed below under the core competency:

**Process of Science**

In your opinion, is it accurate to categorize each of the following **program**-level outcomes under this core competency?

| | Yes | No |
|---|---|---|

| | Yes | No |
| --- | --- | --- |
| Scientific Thinking: Explain how science generates knowledge of the natural world. | ○ | ○ |
| Information Literacy: Locate, interpret, and evaluate scientific information. | ○ | ○ |
| Question Formulation: Pose testable questions and hypotheses to address gaps in knowledge. | ○ | ○ |
| Study Design: Plan, evaluate, and implement scientific investigations. | ○ | ○ |
| Data Interpretation & Evaluation: Interpret, evaluate, and draw conclusions from data in order to make evidence-based arguments about the natural world. | ○ | ○ |
| Doing Research: Integrate process of science skills to address a research question in a course-based or independent research experience. | ○ | ○ |

Optional: Please share any comments you have on the categorization of these outcomes, including any suggestions for alternative categorization if applicable.

You have now reviewed all of the program-level learning outcomes and course-level learning outcomes for this core competency.

Given this review, do you think any essential **program**-level learning outcomes are missing from the **Process of Science** core competency?

Optional: Please share any other feedback on the **Process of Science** core competency.

**Quantitative Reasoning**

In this portion of the survey, we would like you to rate the importance and ease of understanding of our current draft of learning outcomes for one particular core competency: **Quantitative Reasoning.** Later, you will be asked to comment on whether the outcomes are appropriately categorized under this competency.

In the current draft of the BioSkills Guide, each core competency contains multiple **program**-level learning outcomes. In addition, each program-level learning outcome contains multiple **course**-level learning outcomes. Throughout the survey, you will switch between evaluating program- and course-level outcomes. We will use the figure below to remind you of this structure and cue when you are switching between levels.

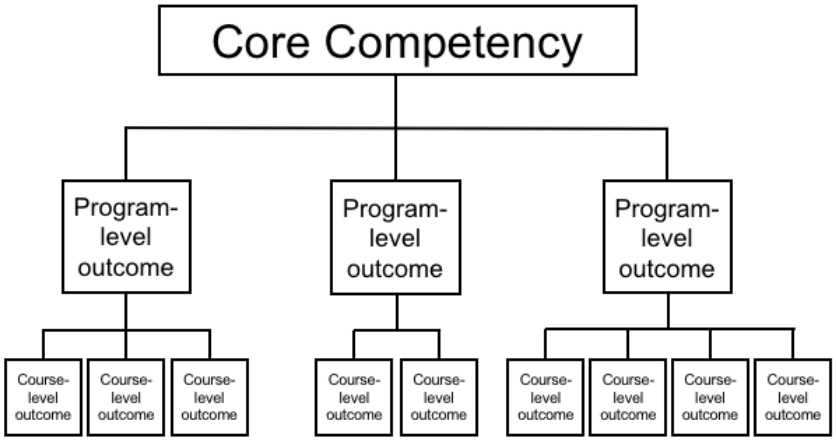


**Important note:** Please keep in mind that we intend for the BioSkills Guide to contain the learning outcomes that we, as a community, think all **graduating general biology majors** should achieve. Therefore, as you complete the survey, please rate the outcomes based on whether they are both desirable and reasonable to accomplish in a **four-year program**, not introductory courses alone (1-2 years only) or in a graduate program (5+ years).

#### Importance and Ease of Understanding: **Quantitative Reasoning**

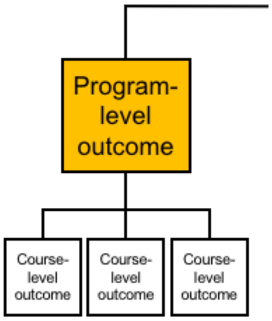


The next three questions will ask you to evaluate the following **program**-level outcome:

**Numeracy: Use basic mathematics (e.g., algebra, probability, unit conversions) in biological contexts.**

How important or unimportant is it for graduating general biology majors to achieve this outcome?

- Very Important
- Important
- Neither Important Nor Unimportant
- Unimportant
- Very Unimportant

How easy or difficult is it for **you** to understand this outcome?

- ◯ Very Easy
- ◯ Easy
- ◯ Neither Easy Nor Difficult
- ◯ Difficult
- ◯ Very Difficult

Please share any feedback you have about the content or wording of this outcome.

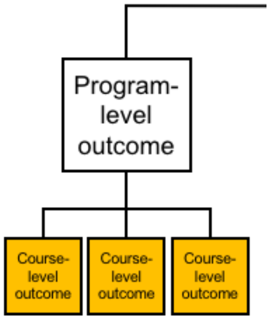


The next three questions will ask you to evaluate the following **course**-level outcome:

**Describe how quantitative reasoning helps biologists understand the natural world.**

How important or unimportant is it for graduating general biology majors to achieve this outcome?

- ◯ Very Important
- ◯ Important
- ◯ Neither Important Nor Unimportant
- ◯ Unimportant
- ◯ Very Unimportant

How easy or difficult is it for **you** to understand this outcome?

- ◯ Very Easy
- ◯ Easy
- ◯ Neither Easy Nor Difficult
- ◯ Difficult
- ◯ Very Difficult

Please share any feedback you have about the content or wording of this outcome.

**Course**-level outcome:

**Perform basic calculations (e.g., percentages, frequencies, rates, means).**

How important or unimportant is it for graduating general biology majors to achieve this outcome?

- Very Important
- Important
- Neither Important Nor Unimportant
- Unimportant
- Very Unimportant

How easy or difficult is it for **you** to understand this outcome?

- Very Easy
- Easy
- Neither Easy Nor Difficult
- Difficult
- Very Difficult

Please share any feedback you have about the content or wording of this outcome.

**Course**-level outcome:

**Select and apply appropriate equations to solve problems (e.g., Hardy-Weinberg equations, Nernst equation, logistic population growth).**

How important or unimportant is it for graduating general biology majors to achieve this outcome?

- Very Important
- Important
- Neither Important Nor Unimportant
- Unimportant
- Very Unimportant

How easy or difficult is it for **you** to understand this outcome?

- Very Easy

- Easy
- Neither Easy Nor Difficult
- Difficult
- Very Difficult

Please share any feedback you have about the content or wording of this outcome.

**Course**-level outcome:

**Interpret and manipulate mathematical relationships (e.g., scale, ratios, units) to make quantitative comparisons.**

How important or unimportant is it for graduating general biology majors to achieve this outcome?

- Very Important
- Important
- Neither Important Nor Unimportant
- Unimportant
- Very Unimportant

How easy or difficult is it for **you** to understand this outcome?

- Very Easy
- Easy
- Neither Easy Nor Difficult
- Difficult
- Very Difficult

Please share any feedback you have about the content or wording of this outcome.

**Course**-level outcome:

**Use probability to reason about biological processes and about statistical analyses.**

How important or unimportant is it for graduating general biology majors to achieve this outcome?

- Very Important
- Important

- Neither Important Nor Unimportant
- Unimportant
- Very Unimportant

How easy or difficult is it for **you** to understand this outcome?

- Very Easy
- Easy
- Neither Easy Nor Difficult
- Difficult
- Very Difficult

Please share any feedback you have about the content or wording of this outcome.

##### **Course**-level outcome:

**Use rough estimates informed by biological knowledge to check quantitative work.**

How important or unimportant is it for graduating general biology majors to achieve this outcome?

- Very Important
- Important
- Neither Important Nor Unimportant
- Unimportant
- Very Unimportant

How easy or difficult is it for **you** to understand this outcome?

- Very Easy
- Easy
- Neither Easy Nor Difficult
- Difficult
- Very Difficult

Please share any feedback you have about the content or wording of this outcome.

#### Importance and Ease of Understanding: **Quantitative Reasoning**

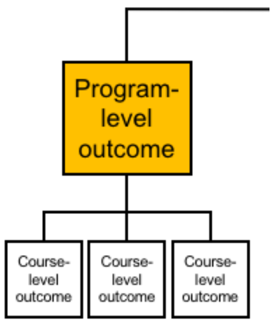


The next three questions will ask you to evaluate the following **program**-level outcome:

**Quantitative & Computational Data Analysis: Apply the tools of graphing, statistics, and data science to analyze biological data.**

How important or unimportant is it for graduating general biology majors to achieve this outcome?

- Very Important
- Important
- Neither Important Nor Unimportant
- Unimportant
- Very Unimportant

How easy or difficult is it for **you** to understand this outcome?

- Very Easy
- Easy
- Neither Easy Nor Difficult
- Difficult
- Very Difficult

Please share any feedback you have about the content or wording of this outcome.

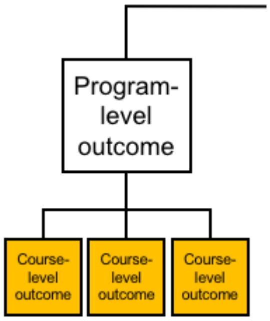


The next three questions will ask you to evaluate the following **course**-level outcome:

**Record, organize, and annotate simple data sets.**

How important or unimportant is it for graduating general biology majors to achieve this outcome?

- Very Important
- Important
- Neither Important Nor Unimportant
- Unimportant
- Very Unimportant

How easy or difficult is it for **you** to understand this outcome?

- Very Easy
- Easy
- Neither Easy Nor Difficult
- Difficult
- Very Difficult

Please share any feedback you have about the content or wording of this outcome.

**Course**-level outcome:

**Create and interpret informative graphs and other data visualizations.**

How important or unimportant is it for graduating general biology majors to achieve this outcome?

- Very Important
- Important
- Neither Important Nor Unimportant
- Unimportant

- Very Unimportant

How easy or difficult is it for **you** to understand this outcome?

- Very Easy
- Easy
- Neither Easy Nor Difficult
- Difficult
- Very Difficult

Please share any feedback you have about the content or wording of this outcome.

##### **Course**-level outcome:

**Select, carry out, and interpret statistical analyses.**

How important or unimportant is it for graduating general biology majors to achieve this outcome?

- Very Important
- Important
- Neither Important Nor Unimportant
- Unimportant
- Very Unimportant

How easy or difficult is it for **you** to understand this outcome?

- Very Easy
- Easy
- Neither Easy Nor Difficult
- Difficult
- Very Difficult

Please share any feedback you have about the content or wording of this outcome.

##### **Course**-level outcome:

**Describe examples of how scientists use databases, large data sets, and data science tools to answer a variety of biological questions.**

How important or unimportant is it for graduating general biology majors to achieve this outcome?

- Very Important
- Important
- Neither Important Nor Unimportant
- Unimportant
- Very Unimportant

How easy or difficult is it for **you** to understand this outcome?

- Very Easy
- Easy
- Neither Easy Nor Difficult
- Difficult
- Very Difficult

Please share any feedback you have about the content or wording of this outcome.

##### **Course**-level outcome:

**Explain the biological meaning of quantitative results.**

How important or unimportant is it for graduating general biology majors to achieve this outcome?

- Very Important
- Important
- Neither Important Nor Unimportant
- Unimportant
- Very Unimportant

How easy or difficult is it for **you** to understand this outcome?

- Very Easy
- Easy
- Neither Easy Nor Difficult
- Difficult
- Very Difficult

Please share any feedback you have about the content or wording of this outcome.

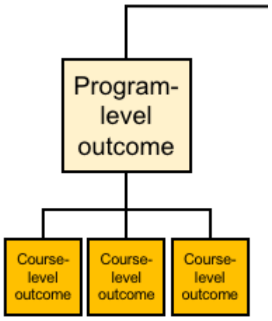


Next, we would like you to indicate whether the current categorization of **course**-level outcomes within program-level outcomes makes sense to you. You will also be asked to suggest missing outcomes.

Some related outcomes may currently be categorized under other competencies. If you would like to see a complete draft of the BioSkills Guide for context, please click here. This is a preliminary draft, and we ask that you *please do not share it with others until it is published.*

#### Categorization: **Quantitative Reasoning**

Currently, we have categorized each of the course-level outcomes listed below under the program-level outcome:

**Numeracy: Use basic mathematics (e.g., algebra, probability, unit conversions) in biological contexts.**

In your opinion, is it accurate to categorize each of the following course-level outcomes under this program-level outcome?

| | Yes | No |
|---|---|---|
| Describe how quantitative reasoning helps biologists understand the natural world. | O | O |
| Perform basic calculations (e.g., percentages, frequencies, rates, means). | O | O |
| Select and apply appropriate equations to solve problems (e.g., Hardy-Weinberg equations, Nernst equation, logistic population growth). | O | O |
| Interpret and manipulate mathematical relationships (e.g., scale, ratios, units) to make quantitative comparisons. | O | O |
| Use probability to reason about biological processes and about statistical analyses. | O | O |
| Use rough estimates informed by biological knowledge to check quantitative work. | O | O |

Program-level outcome:

**Quantitative & Computational Data Analysis: Apply the tools of graphing, statistics, and data science to analyze biological data.**

In your opinion, is it accurate to categorize each of the following course-level outcomes under this program-level outcome?

| | Yes | No |
| --- | --- | --- |
| Record, organize, and annotate simple data sets. | | |
| Create and interpret informative graphs and other data visualizations. | | |
| Select, carry out, and interpret statistical analyses. | | |
| Describe examples of how scientists use databases, large data sets, and data science tools to answer a variety of biological questions. | | |
| Explain the biological meaning of quantitative results. | | |

Optional: Please share any comments you have on the categorization of these outcomes, including any suggestions for alternative categorization if applicable.

Do you think any essential **course**-level outcomes are missing from this list?

#### Categorization: **Quantitative Reasoning**

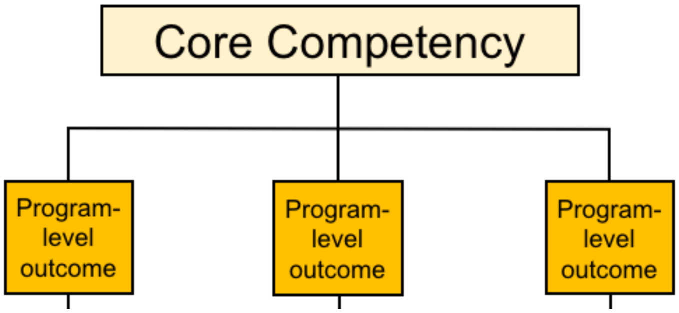


Next, we would like you to indicate if you think any of the **program**-level learning outcomes are currently miscategorized.

If you would like to see the complete, preliminary draft of the BioSkills Guide for context, please click here. As a reminder, the other core competencies are: **Process of Science, Modeling & Simulations, Interdisciplinary Nature of Science, Communication & Collaboration, and Science & Society.**

Currently, we have categorized the program-level outcomes listed below under the core competency:

**Quantitative Reasoning**

In your opinion, is it accurate to categorize each of the following program-level outcomes under this core competency?

| | Yes | No |
|---|---|---|
| Numeracy: Use basic mathematics (e.g., algebra, probability, unit conversions) in biological contexts. | O | O |
| Quantitative & Computational Data Analysis: Apply the tools of graphing, statistics, and data science to analyze biological data. | O | O |

Optional: Please share any comments you have on the categorization of these outcomes, including any suggestions for alternative categorization if applicable.

You have now reviewed all of the program-level learning outcomes and course-level learning outcomes for this core competency.

Given this review, do you think any essential **program**-level learning outcomes are missing from the **Quantitative Reasoning** core competency?

Optional: Please share any other feedback on the **Quantitative Reasoning** core competency.

**Option to Continue**

Thank you for all of your feedback so far! We know that your time is valuable. We would love your feedback on additional outcomes, if you have the time.

Would you like to evaluate another set of outcomes?

#### [***This question was shown after first 2 randomly assigned blocks of questions, and then subsequently after each additional block of questions until all 6 blocks were complete.***]

- O Yes
- O No

**Modeling & Simulation**

In this portion of the survey, we would like you to rate the importance and ease of understanding of our current draft of learning outcomes for one particular core competency: **Modeling and Simulations.** Later, you will be asked to comment on whether the outcomes are appropriately categorized under this competency.

In the current draft of the BioSkills Guide, each core competency contains multiple **program**-level learning outcomes. In addition, each program-level learning outcome contains multiple **course**-level learning outcomes. Throughout the survey, you will switch between evaluating program- and course-level outcomes. We will use the figure below to remind you of this structure and cue when you are switching between levels.

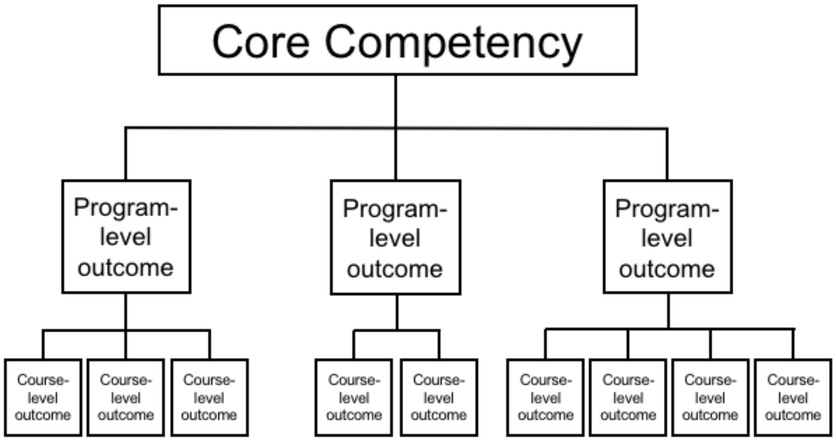


**Important note:** Please keep in mind that we intend for the BioSkills Guide to contain the learning outcomes that we, as a community, think all **graduating general biology majors** should achieve. Therefore, as you complete the survey, please rate the outcomes based on whether they are both desirable and reasonable to accomplish in a **four-year program**, not introductory courses alone (1-2 years only) or in a graduate program (5+ years).

#### Importance and Ease of Understanding: **Modeling & Simulations**

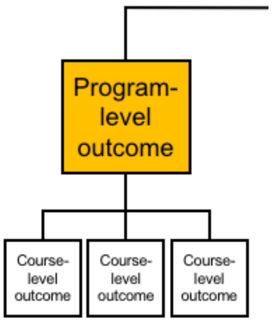


The next three questions will ask you to evaluate the following **program**-level outcome:

**Purpose of Models: Recognize the important roles that scientific models of many different types (conceptual, mathematical, physical, etc.) play in predicting and communicating biological phenomena.**

How important or unimportant is it for graduating general biology majors to achieve this outcome?

- Very Important
- Important
- Neither Important Nor Unimportant
- Unimportant
- Very Unimportant

How easy or difficult is it for **you** to understand this outcome?

- Very Easy
- Easy
- Neither Easy Nor Difficult
- Difficult
- Very Difficult

Please share any feedback you have about the content or wording of this outcome.

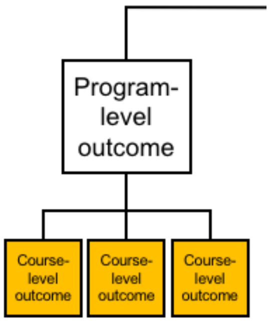


The next three questions will ask you to evaluate the following **course**-level outcome:

**Describe how and why scientists use simplified representations (models) of biological systems when solving problems and communicating ideas.**

How important or unimportant is it for graduating general biology majors to achieve this outcome?

- Very Important
- Important
- Neither Important Nor Unimportant
- Unimportant
- Very Unimportant

How easy or difficult is it for **you** to understand this outcome?

- Very Easy
- Easy
- Neither Easy Nor Difficult
- Difficult
- Very Difficult

Please share any feedback you have about the content or wording of this outcome.

**Course**-level outcome:

**Given two models of the same biological process or system, compare their strengths, limitations, and assumptions.**

How important or unimportant is it for graduating general biology majors to achieve this outcome?

- Very Important
- Important
- Neither Important Nor Unimportant
- Unimportant
- Very Unimportant

How easy or difficult is it for **you** to understand this outcome?

- Very Easy
- Easy
- Neither Easy Nor Difficult
- Difficult
- Very Difficult

Please share any feedback you have about the content or wording of this outcome.

Importance and Ease of Understanding: **Modeling & Simulations**

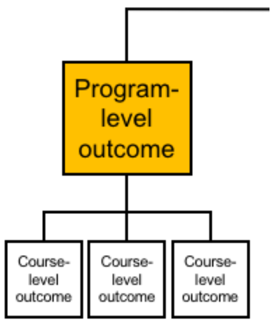


The next three questions will ask you to evaluate the following **program**-level outcome:

**Model Application: Make inferences and solve problems using models and simulations.**

How important or unimportant is it for graduating general biology majors to achieve this outcome?

- ◯ Very Important
- ◯ Important
- ◯ Neither Important Nor Unimportant
- ◯ Unimportant
- ◯ Very Unimportant

How easy or difficult is it for **you** to understand this outcome?

- ◯ Very Easy
- ◯ Easy
- ◯ Neither Easy Nor Difficult
- ◯ Difficult
- ◯ Very Difficult

Please share any feedback you have about the content or wording of this outcome.

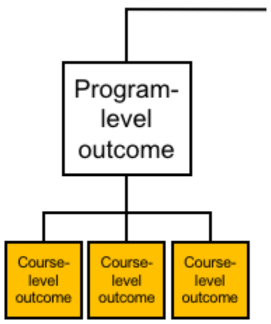

The next three questions will ask you to evaluate the following **course**-level outcome:

**Summarize relationships and trends that can be inferred from a given model or simulation.**

How important or unimportant is it for graduating general biology majors to achieve this outcome?

- ○ Very Important
- ○ Important
- ○ Neither Important Nor Unimportant
- ○ Unimportant
- ○ Very Unimportant

How easy or difficult is it for **you** to understand this outcome?

- ○ Very Easy
- ○ Easy
- ○ Neither Easy Nor Difficult
- ○ Difficult
- ○ Very Difficult

Please share any feedback you have about the content or wording of this outcome.

**Course**-level outcome:

**Use models and simulations to make predictions and refine hypotheses.**

How important or unimportant is it for graduating general biology majors to achieve this outcome?

- ○ Very Important
- ○ Important
- ○ Neither Important Nor Unimportant
- ○ Unimportant
- ○ Very Unimportant

How easy or difficult is it for **you** to understand this outcome?

- ○ Very Easy
- ○ Easy
- ○ Neither Easy Nor Difficult
- ○ Difficult
- ○ Very Difficult

Please share any feedback you have about the content or wording of this outcome.

Importance and Ease of Understanding: **Modeling & Simulations**

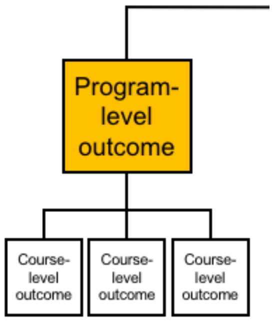


The next three questions will ask you to evaluate the following **program**-level outcome:

**Modeling: Build and evaluate models of biological systems.**

How important or unimportant is it for graduating general biology majors to achieve this outcome?

- Very Important
- Important
- Neither Important Nor Unimportant
- Unimportant
- Very Unimportant

How easy or difficult is it for **you** to understand this outcome?

- Very Easy
- Easy
- Neither Easy Nor Difficult
- Difficult
- Very Difficult

Please share any feedback you have about the content or wording of this outcome.

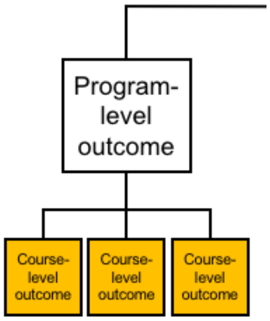


The next three questions will ask you to evaluate the following **course**-level outcome:

**Build and revise conceptual models (e.g., diagrams, concept maps, flow charts) to propose how a biological system or process works.**

How important or unimportant is it for graduating general biology majors to achieve this outcome?

- Very Important
- Important
- Neither Important Nor Unimportant
- Unimportant
- Very Unimportant

How easy or difficult is it for **you** to understand this outcome?

- Very Easy
- Easy
- Neither Easy Nor Difficult
- Difficult
- Very Difficult

Please share any feedback you have about the content or wording of this outcome.

**Course**-level outcome:

**Identify important components of a system and describe how they influence each other (e.g., positively or negatively).**

How important or unimportant is it for graduating general biology majors to achieve this outcome?

- Very Important
- Important

- Neither Important Nor Unimportant
- Unimportant
- Very Unimportant

How easy or difficult is it for **you** to understand this outcome?

- Very Easy
- Easy
- Neither Easy Nor Difficult
- Difficult
- Very Difficult

Please share any feedback you have about the content or wording of this outcome.

**Course**-level outcome:

**Using instructor-provided tools, set parameters of mathematical or computational models and interpret output.**

How important or unimportant is it for graduating general biology majors to achieve this outcome?

- Very Important
- Important
- Neither Important Nor Unimportant
- Unimportant
- Very Unimportant

How easy or difficult is it for **you** to understand this outcome?

- Very Easy
- Easy
- Neither Easy Nor Difficult
- Difficult
- Very Difficult

Please share any feedback you have about the content or wording of this outcome.

**Course**-level outcome:

**Evaluate models by comparing their predictions with empirical data.**

How important or unimportant is it for graduating general biology majors to achieve this outcome?

- Very Important
- Important
- Neither Important Nor Unimportant
- Unimportant
- Very Unimportant

How easy or difficult is it for **you** to understand this outcome?

- Very Easy
- Easy
- Neither Easy Nor Difficult
- Difficult
- Very Difficult

Please share any feedback you have about the content or wording of this outcome.

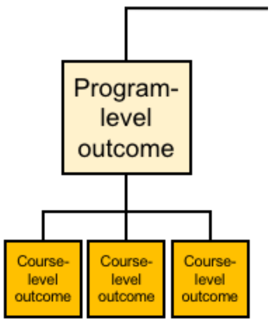


Next, we would like you to indicate whether the current categorization of **course**-level outcomes within program-level outcomes makes sense to you. You will also be asked to suggest missing outcomes.

Some related outcomes may currently be categorized under other competencies. If you would like to see a complete draft of the BioSkills Guide for context, please click here. This is a preliminary draft, and we ask that you *please do not share it with others until it is published.*

Categorization: **Modeling & Simulations**

Currently, we have categorized each of the course-level outcomes listed below under the program-level outcome:

**Purpose of Models: Recognize the important roles that scientific models of many different types (conceptual, mathematical, physical, etc.) play in predicting and communicating biological phenomena.**

In your opinion, is it accurate to categorize each of the following course-level outcomes under this program-level outcome?

| | Yes | No |
|---|---|---|
| Describe how and why scientists use simplified representations (models) of biological systems when solving problems and communicating ideas. | O | O |
| Given two models of the same biological process or system, compare their strengths, limitations, and assumptions. | O | O |

Program-level outcome:

**Model Application: Make inferences and solve problems using models and simulations.**

In your opinion, is it accurate to categorize each of the following course-level outcomes under this program-level outcome?

| | Yes | No |
|---|---|---|
| Summarize relationships and trends that can be inferred from a given model or simulation. | O | O |
| Use models and simulations to make predictions and refine hypotheses. | O | O |

Program-level outcome:

**Modeling: Build and evaluate models of biological systems.**

In your opinion, is it accurate to categorize each of the following course-level outcomes under this program-level outcome?

| | Yes | No |
|---|---|---|
| Build and revise conceptual models (e.g., diagrams, concept maps, flow charts) to propose how a biological system or process works. | O | O |
| Identify important components of a system and describe how they influence each other (e.g., positively or negatively). | O | O |
| Using instructor-provided tools, set parameters of mathematical or computational models and interpret output. | O | O |
| Evaluate models by comparing their predictions with empirical data. | O | O |

Optional: Please share any comments you have on the categorization of these outcomes, including any suggestions for alternative categorization if applicable.

Do you think any essential **course**-level outcomes are missing from this list?

Categorization: **Modeling & Simulations**

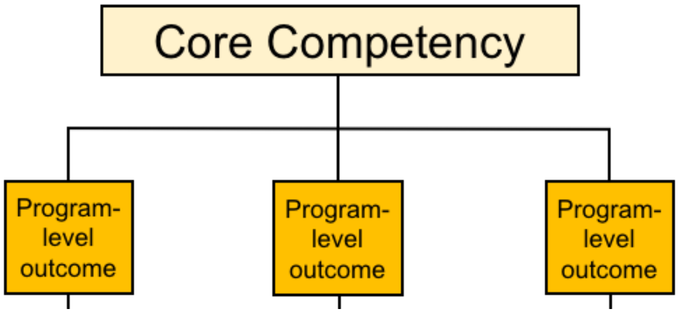


Next, we would like you to indicate if you think any of the **program**-level learning outcomes are currently miscategorized.

If you would like to see the complete, preliminary draft of the BioSkills Guide for context, please click here. As a reminder, the other core competencies are: **Process of Science, Quantitative Reasoning, Interdisciplinary Nature of Science, Communication & Collaboration, and Science & Society.**

Currently, we have categorized the program-level outcomes listed below under the core competency:

**Modeling & Simulations**

In your opinion, is it accurate to categorize each of the following program-level outcomes under this core competency?

| | Yes | No |
|---|---|---|
| Purpose of Models: Recognize the important roles that scientific models of many different types (conceptual, mathematical, physical, etc.) play in predicting and communicating biological phenomena. | ○ | ○ |
| Model Application: Make inferences and solve problems using models and simulations. | ○ | ○ |
| Modeling: Build and evaluate models of biological systems. | ○ | ○ |

Optional: Please share any comments you have on the categorization of these outcomes, including any suggestions for alternative categorization if applicable.

You have now reviewed all of the program-level learning outcomes and course-level learning outcomes for this core competency.

Given this review, do you think there are any essential **program**-level learning outcomes missing from the **Modeling & Simulations** core competency?

Optional: Please share any other feedback on the **Modeling & Simulations** core competency.

**Interdisciplinary Nature of Science**

In this portion of the survey, we would like you to rate the importance and ease of understanding of our current draft of learning outcomes for one particular core competency: **Interdisciplinary Nature of Science.** Later, you will be asked to comment on whether the outcomes are appropriately categorized under this competency.

In the current draft of the BioSkills Guide, each core competency contains multiple **program**-level learning outcomes. In addition, each program-level learning outcome contains multiple **course**-level learning outcomes. Throughout the survey, you will switch between evaluating program- and course-level outcomes. We will use the figure below to remind you of this structure and cue when you are switching between levels.

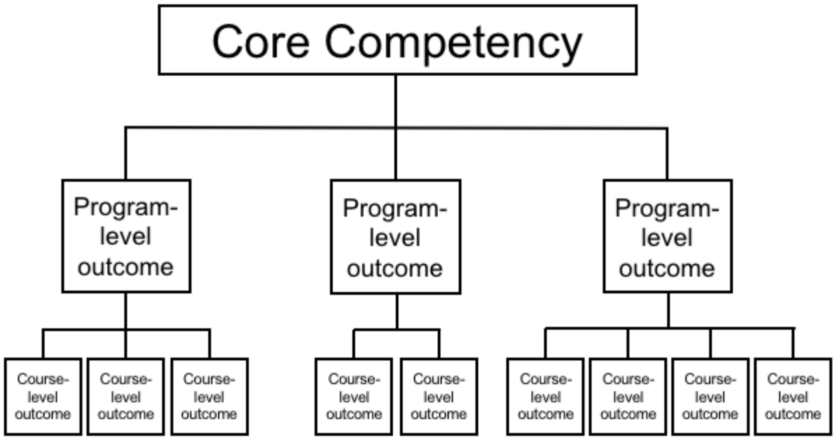


**Important note:** Please keep in mind that we intend for the BioSkills Guide to contain the learning outcomes that we, as a community, think all **graduating general biology majors** should achieve. Therefore, as you complete the survey, please rate the outcomes based on whether they are both desirable and reasonable to accomplish in a **four-year program**, not introductory courses alone (1-2 years only) or in a graduate program (5+ years).

#### Importance and Ease of Understanding: **Interdisciplinary Nature of Science**

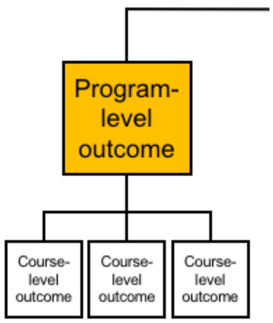


The next three questions will ask you to evaluate the following **program**-level outcome:

**Connecting Scientific Knowledge: Integrate concepts from other STEM disciplines (e.g., chemistry, physics) and across multiple fields of biology (e.g., cell biology, ecology).**

How important or unimportant is it for graduating general biology majors to achieve this outcome?

- Very Important
- Important
- Neither Important Nor Unimportant
- Unimportant
- Very Unimportant

How easy or difficult is it for **you** to understand this outcome?

- Very Easy
- Easy
- Neither Easy Nor Difficult
- Difficult
- Very Difficult

Please share any feedback you have about the content or wording of this outcome.

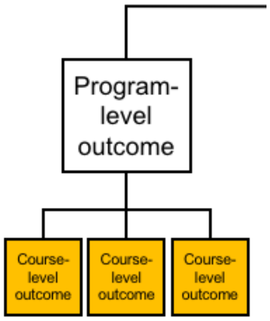


The next three questions will ask you to evaluate the following **course**-level outcome:

**Given a biological problem, identify relevant concepts from other disciplines.**

How important or unimportant is it for graduating general biology majors to achieve this outcome?

- Very Important
- Important
- Neither Important Nor Unimportant
- Unimportant
- Very Unimportant

How easy or difficult is it for **you** to understand this outcome?

- Very Easy
- Easy
- Neither Easy Nor Difficult
- Difficult
- Very Difficult

Please share any feedback you have about the content or wording of this outcome.

**Course**-level outcome:

**Build models or explanations of simple biological processes that include concepts from multiple fields of biology and/or other STEM disciplines.**

How important or unimportant is it for graduating general biology majors to achieve this outcome?

- Very Important
- Important

- Neither Important Nor Unimportant
- Unimportant
- Very Unimportant

How easy or difficult is it for **you** to understand this outcome?

- Very Easy
- Easy
- Neither Easy Nor Difficult
- Difficult
- Very Difficult

Please share any feedback you have about the content or wording of this outcome.

#### Importance and Ease of Understanding: **Interdisciplinary Nature of Science**

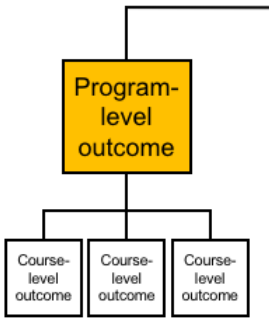


The next three questions will ask you to evaluate the following **program**-level outcome:

**Interdisciplinary Problem Solving: Consider interdisciplinary solutions to real-world problems.**

How important or unimportant is it for graduating general biology majors to achieve this outcome?

- Very Important
- Important
- Neither Important Nor Unimportant
- Unimportant
- Very Unimportant

How easy or difficult is it for **you** to understand this outcome?

- Very Easy
- Easy
- Neither Easy Nor Difficult
- Difficult
- Very Difficult

Please share any feedback you have about the content or wording of this outcome.

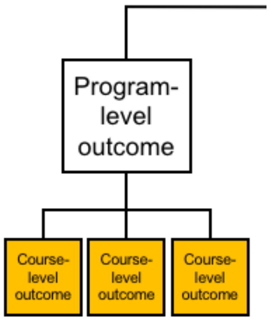


The next three questions will ask you to evaluate the following **course**-level outcome:

**Describe examples of real-world problems that are too complex to be solved by applying biological approaches alone.**

How important or unimportant is it for graduating general biology majors to achieve this outcome?

- Very Important
- Important
- Neither Important Nor Unimportant
- Unimportant
- Very Unimportant

How easy or difficult is it for **you** to understand this outcome?

- Very Easy
- Easy
- Neither Easy Nor Difficult
- Difficult
- Very Difficult

Please share any feedback you have about the content or wording of this outcome.

**Course**-level outcome:

**Suggest how collaborators in other disciplines could contribute to solutions of real-world problems.**

How important or unimportant is it for graduating general biology majors to achieve this outcome?

- Very Important
- Important
- Neither Important Nor Unimportant
- Unimportant
- Very Unimportant

How easy or difficult is it for **you** to understand this outcome?

- Very Easy
- Easy
- Neither Easy Nor Difficult
- Difficult
- Very Difficult

Please share any feedback you have about the content or wording of this outcome.

**Course**-level outcome:

**Be able to explain biological concepts, data, and methods, including their limitations, using language understandable by collaborators in other disciplines.**

How important or unimportant is it for graduating general biology majors to achieve this outcome?

- Very Important
- Important
- Neither Important Nor Unimportant
- Unimportant
- Very Unimportant

How easy or difficult is it for **you** to understand this outcome?

- Very Easy
- Easy
- Neither Easy Nor Difficult
- Difficult
- Very Difficult

Please share any feedback you have about the content or wording of this outcome.

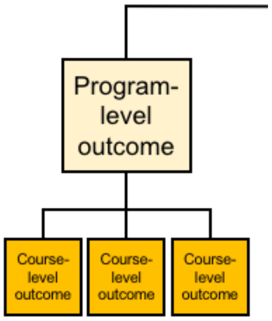


Next, we would like you to indicate whether the current categorization of **course**-level outcomes within program-level outcomes makes sense to you. You will also be asked to suggest missing outcomes.

Some related outcomes may currently be categorized under other competencies. If you would like to see a complete draft of the BioSkills Guide for context, please click here. This is a preliminary draft, and we ask that you *please do not share it with others until it is published.*

#### Categorization: **Interdisciplinary Nature of Science**

Currently, we have categorized each of the course-level outcomes listed below under the program-level outcome:

**Connecting Scientific Knowledge: Integrate concepts from other STEM disciplines (e.g., chemistry, physics) and across multiple fields of biology (e.g., cell biology, ecology).**

In your opinion, is it accurate to categorize each of the following course-level outcomes under this program-level outcome?

| | Yes | No |
|---|---|---|
| Given a biological problem, identify relevant concepts from other disciplines. | ○ | ○ |
| Build models or explanations of simple biological processes that include concepts from multiple fields of biology and/or other STEM disciplines. | ○ | ○ |

Program-level outcome:

**Interdisciplinary Problem Solving: Consider interdisciplinary solutions to real-world problems.**

In your opinion, is it accurate to categorize each of the following course-level outcomes under this program-level outcome?

| | Yes | No |
|---|---|---|
| Describe examples of real-world problems that are too complex to be solved by applying biological approaches alone. | ○ | ○ |
| Suggest how collaborators in other disciplines could contribute to solutions of real-world problems. | ○ | ○ |
| Be able to explain biological concepts, data, and methods, including their limitations, using language understandable by collaborators in other disciplines. | ○ | ○ |

Optional: Please share any comments you have on the categorization of these outcomes, including any suggestions for alternative categorization if applicable.

Do you think any essential **course**-level outcomes are missing from this list?

Categorization: **Interdisciplinary Nature of Science**

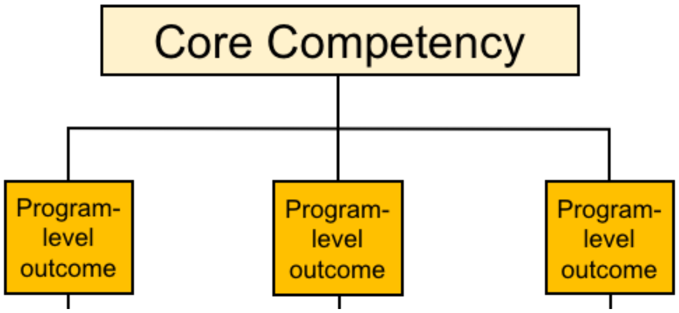


Next, we would like you to indicate if you think any of the **program**-level learning outcomes are currently miscategorized.

If you would like to see the complete, preliminary draft of the BioSkills Guide for context, please click here. As a reminder, the other core competencies are: **Process of Science, Quantitative Reasoning, Modeling & Simulations, Communication & Collaboration, and Science & Society.**

Currently, we have categorized the program-level outcomes listed below under the core competency:

**Interdisciplinary Nature of Science**

In your opinion, is it accurate to categorize each of the following program-level outcomes under this core competency?

| | Yes | No |
|---|---|---|
| Connecting Scientific Knowledge: Integrate concepts from other STEM disciplines (e.g., chemistry, physics) and across multiple fields of biology (e.g., cell biology, ecology). | O | O |
| Interdisciplinary Problem Solving: Consider interdisciplinary solutions to real-world problems. | O | O |

Optional: Please share any comments you have on the categorization of these outcomes, including any suggestions for alternative categorization if applicable.

You have now reviewed all of the program-level learning outcomes and course-level learning outcomes for this core competency.

Given this review, do you think there are any essential **program**-level learning outcomes missing from the **Interdisciplinary Nature of Science** core competency?

Optional: Please share any other feedback on the **Interdisciplinary Nature of Science** core competency.

**Communication & Collaboration**

In this portion of the survey, we would like you to rate the importance and ease of understanding of our current draft of learning outcomes for one particular core competency: **Communication & Collaboration.** Later, you will be asked to comment on whether the outcomes are appropriately categorized under this competency.

In the current draft of the BioSkills Guide, each core competency contains multiple **program**-level learning outcomes. In addition, each program-level learning outcome contains multiple **course**-level learning outcomes. Throughout the survey, you will switch between evaluating program- and course-level outcomes. We will use the figure below to remind you of this structure and cue when you are switching between levels.

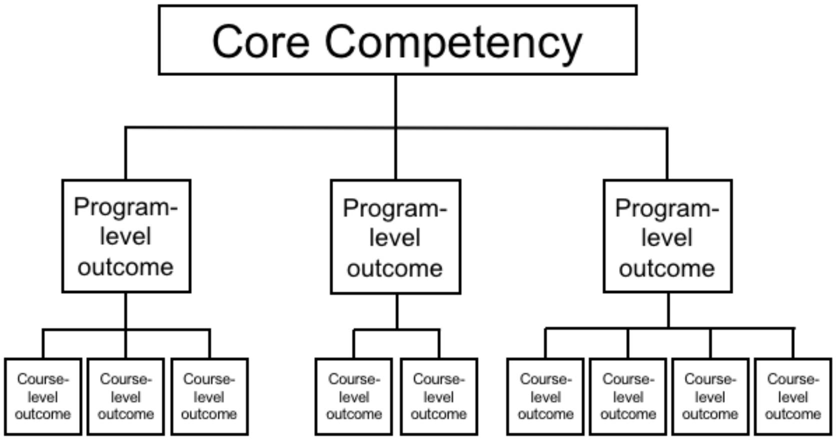


**Important note:** Please keep in mind that we intend for the BioSkills Guide to contain the learning outcomes that we, as a community, think all **graduating general biology majors** should achieve. Therefore, as you complete the survey, please rate the outcomes based on whether they are both desirable and reasonable to accomplish in a **four-year program**, not introductory courses alone (1-2 years only) or in a graduate program (5+ years).

Importance and Ease of Understanding: **Communication & Collaboration**

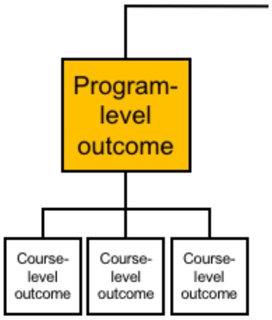


The next three questions will ask you to evaluate the following **program**-level outcome:

**Communication: Share ideas, data, and findings with others clearly and accurately.**

How important or unimportant is it for graduating general biology majors to achieve this outcome?

- ○ Very Important
- ○ Important
- ○ Neither Important Nor Unimportant
- ○ Unimportant
- ○ Very Unimportant

How easy or difficult is it for **you** to understand this outcome?

- ○ Very Easy

- Easy
- Neither Easy Nor Difficult
- Difficult
- Very Difficult

Please share any feedback you have about the content or wording of this outcome.

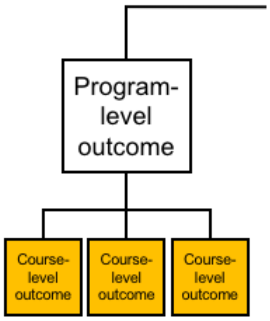


The next three questions will ask you to evaluate the following **course**-level outcome:

**Use an appropriate voice to communicate science to targeted audiences (e.g., general public, biology experts, collaborators in other disciplines).**

How important or unimportant is it for graduating general biology majors to achieve this outcome?

- Very Important
- Important
- Neither Important Nor Unimportant
- Unimportant
- Very Unimportant

How easy or difficult is it for **you** to understand this outcome?

- Very Easy
- Easy
- Neither Easy Nor Difficult
- Difficult
- Very Difficult

Please share any feedback you have about the content or wording of this outcome.

**Course**-level outcome:

**Use multiple modes to communicate science (e.g., oral, written, visual).**

How important or unimportant is it for graduating general biology majors to achieve this outcome?

- ○ Very Important
- ○ Important
- ○ Neither Important Nor Unimportant
- ○ Unimportant
- ○ Very Unimportant

How easy or difficult is it for **you** to understand this outcome?

- ○ Very Easy
- ○ Easy
- ○ Neither Easy Nor Difficult
- ○ Difficult
- ○ Very Difficult

Please share any feedback you have about the content or wording of this outcome.

**Course**-level outcome:

**Describe the purpose and parts of different forms of scientific communication (e.g., journal articles, posters, grant proposals).**

How important or unimportant is it for graduating general biology majors to achieve this outcome?

- ○ Very Important
- ○ Important
- ○ Neither Important Nor Unimportant
- ○ Unimportant
- ○ Very Unimportant

How easy or difficult is it for **you** to understand this outcome?

- ○ Very Easy
- ○ Easy
- ○ Neither Easy Nor Difficult
- ○ Difficult

◯ Very Difficult

Please share any feedback you have about the content or wording of this outcome.

Importance and Ease of Understanding: **Communication & Collaboration**

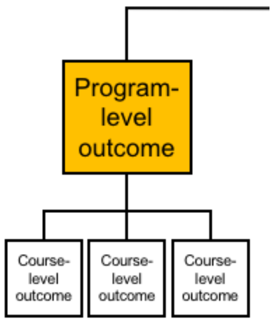


The next three questions will ask you to evaluate the following **program**-level outcome:

**Collaboration: Work productively in teams with people who have diverse backgrounds, skill sets, and perspectives.**

How important or unimportant is it for graduating general biology majors to achieve this outcome?

◯ Very Important
◯ Important
◯ Neither Important Nor Unimportant
◯ Unimportant
◯ Very Unimportant

How easy or difficult is it for **you** to understand this outcome?

◯ Very Easy
◯ Easy
◯ Neither Easy Nor Difficult
◯ Difficult
◯ Very Difficult

Please share any feedback you have about the content or wording of this outcome.

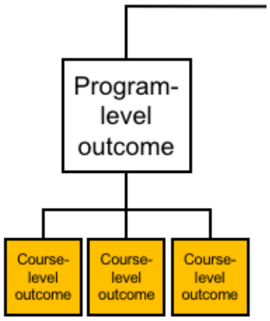


The next three questions will ask you to evaluate the following **course**-level outcome:

**Break team projects into tasks and decide how they can be productively shared.**

How important or unimportant is it for graduating general biology majors to achieve this outcome?

- ○ Very Important
- ○ Important
- ○ Neither Important Nor Unimportant
- ○ Unimportant
- ○ Very Unimportant

How easy or difficult is it for **you** to understand this outcome?

- ○ Very Easy
- ○ Easy
- ○ Neither Easy Nor Difficult
- ○ Difficult
- ○ Very Difficult

Please share any feedback you have about the content or wording of this outcome.

**Course**-level outcome:

**Work with teammates to establish and periodically update group expectations (e.g., project timeline, rules for group interactions).**

How important or unimportant is it for graduating general biology majors to achieve this outcome?

- Very Important
- Important
- Neither Important Nor Unimportant
- Unimportant
- Very Unimportant

How easy or difficult is it for **you** to understand this outcome?

- Very Easy
- Easy
- Neither Easy Nor Difficult
- Difficult
- Very Difficult

Please share any feedback you have about the content or wording of this outcome.

**Course**-level outcome:

**Elicit, listen to, and incorporate ideas from teammates with diverse perspectives and backgrounds.**

How important or unimportant is it for graduating general biology majors to achieve this outcome?

- Very Important
- Important
- Neither Important Nor Unimportant
- Unimportant
- Very Unimportant

How easy or difficult is it for **you** to understand this outcome?

- Very Easy
- Easy
- Neither Easy Nor Difficult
- Difficult
- Very Difficult

Please share any feedback you have about the content or wording of this outcome.

Course-level outcome:

**Work effectively with teammates to complete projects.**

How important or unimportant is it for graduating general biology majors to achieve this outcome?

- Very Important
- Important
- Neither Important Nor Unimportant
- Unimportant
- Very Unimportant

How easy or difficult is it for **you** to understand this outcome?

- Very Easy
- Easy
- Neither Easy Nor Difficult
- Difficult
- Very Difficult

Please share any feedback you have about the content or wording of this outcome.

Importance and Ease of Understanding: **Communication & Collaboration**

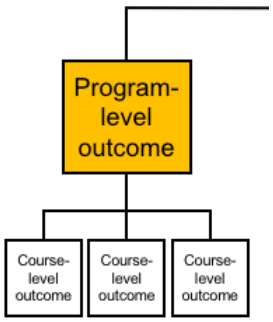


The next three questions will ask you to evaluate the following **program**-level outcome:

**Collegial Review: Provide and respond to constructive feedback in order to improve individual and team work.**

How important or unimportant is it for graduating general biology majors to achieve this outcome?

- Very Important
- Important
- Neither Important Nor Unimportant
- Unimportant
- Very Unimportant

How easy or difficult is it for **you** to understand this outcome?

- Very Easy
- Easy
- Neither Easy Nor Difficult
- Difficult
- Very Difficult

Please share any feedback you have about the content or wording of this outcome.

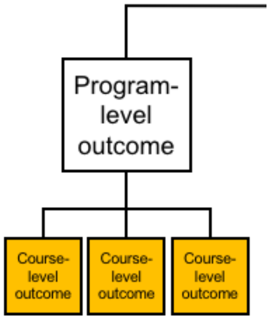


The next three questions will ask you to evaluate the following **course**-level outcome:

**Evaluate feedback from others and revise work or behavior appropriately.**

How important or unimportant is it for graduating general biology majors to achieve this outcome?

- Very Important
- Important
- Neither Important Nor Unimportant
- Unimportant
- Very Unimportant

How easy or difficult is it for **you** to understand this outcome?

- Very Easy

- Easy
- Neither Easy Nor Difficult
- Difficult
- Very Difficult

Please share any feedback you have about the content or wording of this outcome.

##### **Course**-level outcome:

**Critique others' work and ideas constructively and respectfully.**

How important or unimportant is it for graduating general biology majors to achieve this outcome?

- Very Important
- Important
- Neither Important Nor Unimportant
- Unimportant
- Very Unimportant

How easy or difficult is it for **you** to understand this outcome?

- Very Easy
- Easy
- Neither Easy Nor Difficult
- Difficult
- Very Difficult

Please share any feedback you have about the content or wording of this outcome.

#### Importance and Ease of Understanding: **Communication & Collaboration**

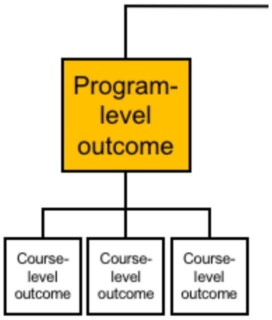


The next three questions will ask you to evaluate the following **program**-level outcome:

**Metacognition: Reflect on your own learning, performance, and achievements.**

How important or unimportant is it for graduating general biology majors to achieve this outcome?

- Very Important
- Important
- Neither Important Nor Unimportant
- Unimportant
- Very Unimportant

How easy or difficult is it for **you** to understand this outcome?

- Very Easy
- Easy
- Neither Easy Nor Difficult
- Difficult
- Very Difficult

Please share any feedback you have about the content or wording of this outcome.

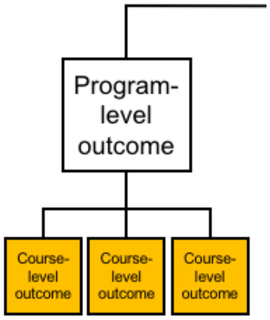

The next three questions will ask you to evaluate the following **course**-level outcome:

**Evaluate your own understanding and skill level.**

How important or unimportant is it for graduating general biology majors to achieve this outcome?

- ○ Very Important
- ○ Important
- ○ Neither Important Nor Unimportant
- ○ Unimportant
- ○ Very Unimportant

How easy or difficult is it for **you** to understand this outcome?

- ○ Very Easy
- ○ Easy
- ○ Neither Easy Nor Difficult
- ○ Difficult
- ○ Very Difficult

Please share any feedback you have about the content or wording of this outcome.

**Course**-level outcome:

**Assess personal progress and contributions to your team and generate a plan to change your behavior as needed.**

How important or unimportant is it for graduating general biology majors to achieve this outcome?

- ○ Very Important
- ○ Important
- ○ Neither Important Nor Unimportant
- ○ Unimportant
- ○ Very Unimportant

How easy or difficult is it for **you** to understand this outcome?

- ○ Very Easy
- ○ Easy
- ○ Neither Easy Nor Difficult
- ○ Difficult

Very Difficult

Please share any feedback you have about the content or wording of this outcome.

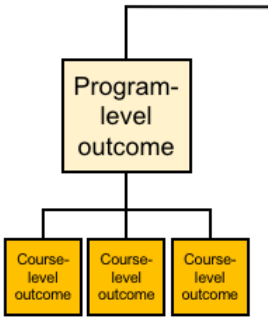


Next, we would like you to indicate whether the current categorization of **course**-level outcomes within program-level outcomes makes sense to you. You will also be asked to suggest missing outcomes.

Some related outcomes may currently be categorized under other competencies. If you would like to see a complete draft of the BioSkills Guide for context, please click here. This is a preliminary draft, and we ask that you *please do not share it with others until it is published.*

#### Categorization: **Communication & Collaboration**

Currently, we have categorized each of the course-level outcomes listed below under the program-level outcome:

**Communication: Share ideas, data, and findings with others clearly and accurately.**

In your opinion, is it accurate to categorize each of the following course-level outcomes under this program-level outcome?

| | Yes | No |
|---|---|---|
| Use an appropriate voice to communicate science to targeted audiences (e.g., general public, biology experts, collaborators in other disciplines). | | |
| Use multiple modes to communicate science (e.g., oral, written, visual). | | |
| Describe the purpose and parts of different forms of scientific communication (e.g., journal articles, posters, grant proposals). | | |

Program-level outcome:

**Collaboration: Work productively in teams with people who have diverse backgrounds, skill sets, and perspectives.**

In your opinion, is it accurate to categorize each of the following course-level outcomes under this program-level outcome?

| | Yes | No |
|---|---|---|
| Break team projects into tasks and decide how they can be productively shared. | O | O |
| Work with teammates to establish and periodically update group expectations (e.g., project timeline, rules for group interactions). | O | O |
| Elicit, listen to, and incorporate ideas from teammates with diverse perspectives and backgrounds. | O | O |
| Work effectively with teammates to complete projects. | O | O |

Program-level outcome:

**Collegial Review: Provide and respond to constructive feedback in order to improve individual and team work.**

In your opinion, is it accurate to categorize each of the following course-level outcomes under this program-level outcome?

| | Yes | No |
|---|---|---|
| Evaluate feedback from others and revise work or behavior appropriately. | O | O |
| Critique others' work and ideas constructively and respectfully. | O | O |

Program-level outcome:

**Metacognition: Reflect on your own learning, performance, and achievements.**

In your opinion, is it accurate to categorize each of the following course-level outcomes under this program-level outcome?

| | Yes | No |
|---|---|---|
| Evaluate your own understanding and skill level. | O | O |
| Assess personal progress and contributions to your team and generate a plan to change your behavior as needed. | O | O |

Optional: Please share any comments you have on the categorization of these outcomes, including any suggestions for alternative categorization if applicable.

Do you think any essential **course**-level outcomes are missing from this list?

Categorization: **Communication & Collaboration**

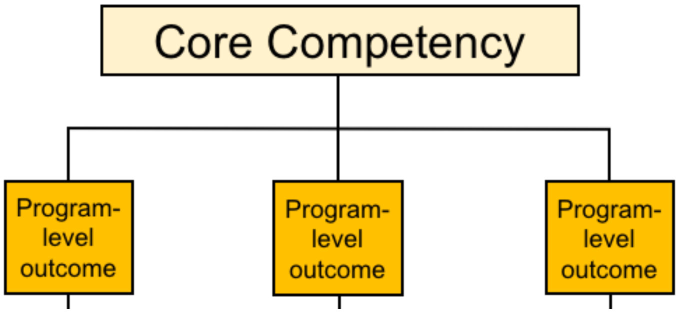


Next, we would like you to indicate if you think any of the **program**-level learning outcomes are currently miscategorized.

If you would like to see the complete, preliminary draft of the BioSkills Guide for context, please click here. As a reminder, the other core competencies are: **Process of Science, Quantitative Reasoning, Modeling & Simulations, Interdisciplinary Nature of Science, and Science & Society.**

Currently, we have categorized the program-level outcomes listed below under the core competency:

**Communication & Collaboration**

In your opinion, is it accurate to categorize each of the following program-level outcomes under this core competency?

| | Yes | No |
|---|---|---|
| Communication: Share ideas, data, and findings with others clearly and accurately. | ○ | ○ |
| Collaboration: Work productively in teams with people who have diverse backgrounds, skill sets, and perspectives. | ○ | ○ |
| Collegial Review: Provide and respond to constructive feedback in order to improve individual and team work. | ○ | ○ |
| Metacognition: Reflect on your own learning, performance, and achievements. | ○ | ○ |

Optional: Please share any comments you have on the categorization of these outcomes, including any suggestions for alternative categorization if applicable.

You have now reviewed all of the program-level learning outcomes and course-level learning outcomes for this core competency.

Given this review, do you think any essential **program**-level learning outcomes are missing from the **Communication & Collaboration** core competency?

Optional: Please share any other feedback on the **Communication & Collaboration** core competency.

**Science & Society**

In this portion of the survey, we would like you to rate the importance and ease of understanding of our current draft of learning outcomes for one particular core competency: **Science & Society.** Later, you will be asked to comment on whether the outcomes are appropriately categorized under this competency.

In the current draft of the BioSkills Guide, each core competency contains multiple **program**-level learning outcomes. In addition, each program-level learning outcome contains multiple **course**-level learning outcomes. Throughout the survey, you will switch between evaluating program- and course-level outcomes. We will use the figure below to remind you of this structure and cue when you are switching between levels.

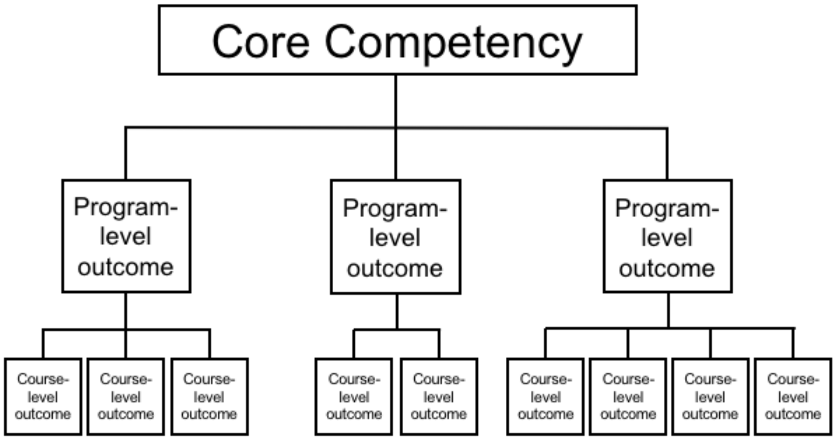


**Important note:** Please keep in mind that we intend for the BioSkills Guide to contain the learning outcomes that we, as a community, think all **graduating general biology majors** should achieve. Therefore, as you complete the survey, please rate the outcomes based on whether they are both desirable and reasonable to accomplish in a **four-year program**, not introductory courses alone (1-2 years only) or in a graduate program (5+ years).

#### Importance and Ease of Understanding: **Science & Society**

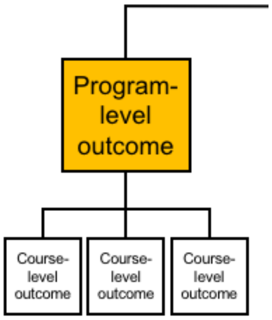


The next three questions will ask you to evaluate the following **program**-level outcome:

**Ethics: Demonstrate the ability to think critically about ethical issues in the conduct of science.**

How important or unimportant is it for graduating general biology majors to achieve this outcome?

- Very Important
- Important
- Neither Important Nor Unimportant
- Unimportant
- Very Unimportant

How easy or difficult is it for **you** to understand this outcome?

- Very Easy
- Easy
- Neither Easy Nor Difficult
- Difficult
- Very Difficult

Please share any feedback you have about the content or wording of this outcome.

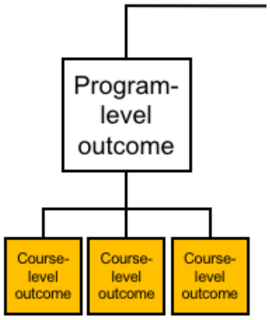

The next three questions will ask you to evaluate the following **course**-level outcome:

**Evaluate ethical considerations (e.g., use of animal or human subjects, conflicts of interest, biased study design) in a given research study.**

How important or unimportant is it for graduating general biology majors to achieve this outcome?

- ◯ Very Important
- ◯ Important
- ◯ Neither Important Nor Unimportant
- ◯ Unimportant
- ◯ Very Unimportant

How easy or difficult is it for **you** to understand this outcome?

- ◯ Very Easy
- ◯ Easy
- ◯ Neither Easy Nor Difficult
- ◯ Difficult
- ◯ Very Difficult

Please share any feedback you have about the content or wording of this outcome.

**Course**-level outcome:

**Provide examples of ethical controversies in biological research and critique how they have been addressed by the scientific community.**

How important or unimportant is it for graduating general biology majors to achieve this outcome?

- ◯ Very Important
- ◯ Important
- ◯ Neither Important Nor Unimportant
- ◯ Unimportant
- ◯ Very Unimportant

How easy or difficult is it for **you** to understand this outcome?

- ◯ Very Easy
- ◯ Easy
- ◯ Neither Easy Nor Difficult
- ◯ Difficult

○ Very Difficult

Please share any feedback you have about the content or wording of this outcome.

Importance and Ease of Understanding: **Science & Society**

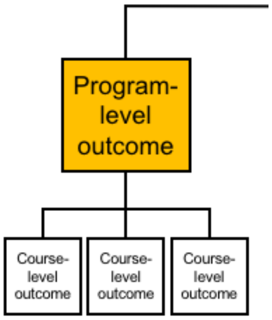


The next three questions will ask you to evaluate the following **program**-level outcome:

**Societal Influences: Consider the potential impacts of outside influences (historical, cultural, political, technological) on how science is practiced.**

How important or unimportant is it for graduating general biology majors to achieve this outcome?

○ Very Important
○ Important
○ Neither Important Nor Unimportant
○ Unimportant
○ Very Unimportant

How easy or difficult is it for **you** to understand this outcome?

○ Very Easy
○ Easy
○ Neither Easy Nor Difficult
○ Difficult
○ Very Difficult

Please share any feedback you have about the content or wording of this outcome.

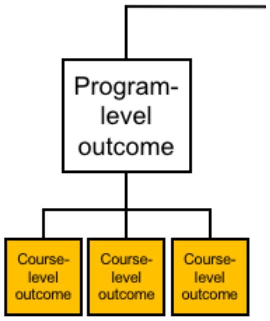


The next three questions will ask you to evaluate the following **course**-level outcome:

**Describe examples of how scientists' backgrounds and biases can influence science and how science is enhanced through diversity.**

How important or unimportant is it for graduating general biology majors to achieve this outcome?

- Very Important
- Important
- Neither Important Nor Unimportant
- Unimportant
- Very Unimportant

How easy or difficult is it for **you** to understand this outcome?

- Very Easy
- Easy
- Neither Easy Nor Difficult
- Difficult
- Very Difficult

Please share any feedback you have about the content or wording of this outcome.

**Course**-level outcome:

**Identify and describe how systemic factors (e.g., socioeconomic, political) affect how and by whom science is conducted.**

How important or unimportant is it for graduating general biology majors to achieve this outcome?

- Very Important
- Important
- Neither Important Nor Unimportant
- Unimportant
- Very Unimportant

How easy or difficult is it for **you** to understand this outcome?

- Very Easy
- Easy
- Neither Easy Nor Difficult
- Difficult
- Very Difficult

Please share any feedback you have about the content or wording of this outcome.

#### Importance and Ease of Understanding: **Science & Society**

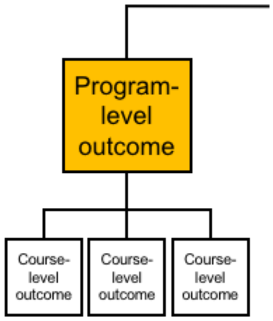


The next three questions will ask you to evaluate the following **program**-level outcome:

**Science's Impact on Society: Apply scientific reasoning in daily life and recognize the impacts of science on a local and global scale.**

How important or unimportant is it for graduating general biology majors to achieve this outcome?

- Very Important
- Important
- Neither Important Nor Unimportant
- Unimportant

○ Very Unimportant

How easy or difficult is it for **you** to understand this outcome?

○ Very Easy
○ Easy
○ Neither Easy Nor Difficult
○ Difficult
○ Very Difficult

Please share any feedback you have about the content or wording of this outcome.

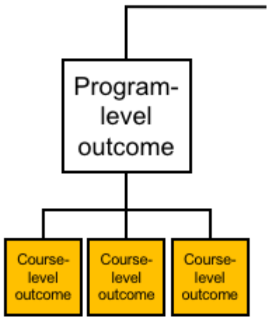


The next three questions will ask you to evaluate the following **course**-level outcome:

**Apply evidence-based reasoning and biological knowledge in daily life (e.g., consuming popular media, deciding how to vote).**

How important or unimportant is it for graduating general biology majors to achieve this outcome?

○ Very Important
○ Important
○ Neither Important Nor Unimportant
○ Unimportant
○ Very Unimportant

How easy or difficult is it for **you** to understand this outcome?

○ Very Easy
○ Easy
○ Neither Easy Nor Difficult
○ Difficult
○ Very Difficult

Please share any feedback you have about the content or wording of this outcome.

**Course**-level outcome:

**Use examples to describe the relevance of science in everyday experiences.**

How important or unimportant is it for graduating general biology majors to achieve this outcome?

- Very Important
- Important
- Neither Important Nor Unimportant
- Unimportant
- Very Unimportant

How easy or difficult is it for **you** to understand this outcome?

- Very Easy
- Easy
- Neither Easy Nor Difficult
- Difficult
- Very Difficult

Please share any feedback you have about the content or wording of this outcome.

**Course**-level outcome:

**Identify and describe the broader societal impacts of biological research on different stakeholders.**

How important or unimportant is it for graduating general biology majors to achieve this outcome?

- Very Important
- Important
- Neither Important Nor Unimportant
- Unimportant
- Very Unimportant

How easy or difficult is it for **you** to understand this outcome?

- Very Easy
- Easy
- Neither Easy Nor Difficult
- Difficult
- Very Difficult

Please share any feedback you have about the content or wording of this outcome.

**Course**-level outcome:

**Describe the role scientists have in providing public access to scientific knowledge.**

How important or unimportant is it for graduating general biology majors to achieve this outcome?

- Very Important
- Important
- Neither Important Nor Unimportant
- Unimportant
- Very Unimportant

How easy or difficult is it for **you** to understand this outcome?

- Very Easy
- Easy
- Neither Easy Nor Difficult
- Difficult
- Very Difficult

Please share any feedback you have about the content or wording of this outcome.

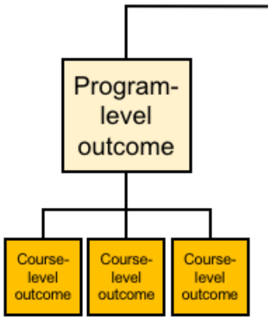


Next, we would like you to indicate whether the current categorization of **course**-level outcomes within program-level outcomes makes sense to you. You will also be asked to suggest missing outcomes.

Some related outcomes may currently be categorized under other competencies. If you would like to see a complete draft of the BioSkills Guide for context, please click here. This is a preliminary draft, and we ask that you *please do not share it with others until it is published*.

#### Categorization: **Science & Society**

Currently, we have categorized each of the course-level outcomes listed below under the program-level outcome:

**Ethics: Demonstrate the ability to think critically about ethical issues in the conduct of science.**

In your opinion, is it accurate to categorize each of the following course-level outcomes under this program-level outcome?

| | Yes | No |
|---|---|---|
| Evaluate ethical considerations (e.g., use of animal or human subjects, conflicts of interest, biased study design) in a given research study. | O | O |
| Provide examples of ethical controversies in biological research and critique how they have been addressed by the scientific community. | O | O |

Program-level outcome:

**Societal Influences: Consider the potential impacts of outside influences (historical, cultural, political, technological) on how science is practiced.**

In your opinion, is it accurate to categorize the following course-level outcome under this program-level outcome?

| | Yes | No |
|---|---|---|
| Describe examples of how scientists' backgrounds and biases can influence science and how science is enhanced through diversity. | O | O |

| | Yes | No |
|---|---|---|
| Identify and describe how systemic factors (e.g., socioeconomic, political) affect how and by whom science is conducted. | ○ | ○ |

Program-level outcome:

**Science’s Impact on Society: Apply scientific reasoning in daily life and recognize the impacts of science on a local and global scale.**

In your opinion, is it accurate to categorize each of the following course-level outcomes under this program-level outcome?

| | Yes | No |
|---|---|---|
| Apply evidence-based reasoning and biological knowledge in daily life (e.g., consuming popular media, deciding how to vote). | ○ | ○ |
| Use examples to describe the relevance of science in everyday experiences. | ○ | ○ |
| Identify and describe the broader societal impacts of biological research on different stakeholders. | ○ | ○ |
| Describe the role scientists have in providing public access to scientific knowledge. | ○ | ○ |

Optional: Please share any comments you have on the categorization of these outcomes, including any suggestions for alternative categorization if applicable.

Do you think any essential **course**-level outcomes are missing from this list?

Categorization: **Science & Society**

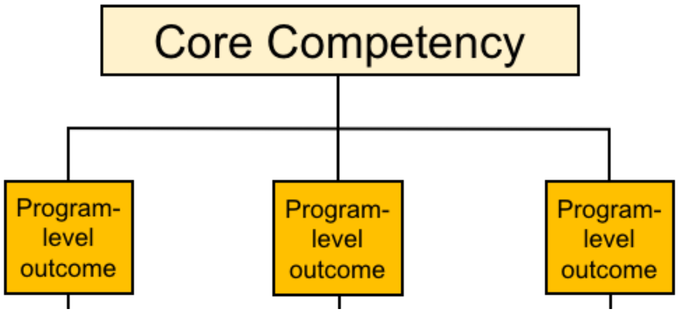

Next, we would like you to indicate if you think any of **program**-level learning outcomes are currently miscategorized.

If you would like to see the complete, preliminary draft of the BioSkills Guide for context, please click here. As a reminder, the other core competencies are: **Process of Science, Quantitative Reasoning, Modeling & Simulations, Interdisciplinary Nature of Science, and Communication & Collaboration.**

Currently, we have categorized the program-level outcomes listed below under the core competency:

**Science & Society**

In your opinion, is it accurate to categorize each of the following program-level outcomes under this core competency?

| | Yes | No |
|---|---|---|
| Ethics: Demonstrate the ability to think critically about ethical issues in the conduct of science. | O | O |
| Societal Influences: Consider the potential impacts of outside influences (historical, cultural, political, technological) on how science is practiced. | O | O |
| Science's Impact on Society: Apply scientific reasoning in daily life and recognize the impacts of science on a local and global scale. | O | O |

Optional: Please share any comments you have on the categorization of these outcomes, including any suggestions for alternative categorization if applicable.

You have now reviewed all of the program-level learning outcomes and course-level learning outcomes for this core competency.

Given this review, do you think any essential **program**-level learning outcomes are missing from the **Science & Society** core competency?

Optional: Please share any other feedback on the **Science & Society** core competency.

**Demographics**

#### Demographic Questions

We ask that you complete the following demographic questions so that we can determine if we are gathering feedback from a representative population. We will not link specific responses with any individual identifying information when sharing the results of this survey.

What is the name of your current institution? (This will be used to gather additional institutional demographic information.)

____________________

Which of the following best describes your institution type?

- ○ Associate’s Degree-Granting
- ○ Bachelor’s Degree-Granting
- ○ Master’s Degree-Granting
- ○ Doctoral Degree-Granting
- ○ ________ Other (please specify):

Which of the following best describes your current position?

- ○ Graduate Student
- ○ Postdoc
- ○ Lecturer or Instructor
- ○ Assistant, Associate, or Full Professor
- ○ Staff
- ○ ________ Other (please specify):

In your current position, what is your primary responsibility?

- ○ Teaching
- ○ Research
- ○ Teaching and Research Equally
- ○ ________ Other (please describe briefly)

What is the focus of your current research, if applicable? (please select all that apply)

- ☐ I am not currently engaged in research
- ☐ Molecular/Cellular/Developmental Biology
- ☐ Physiology
- ☐ Ecology/Evolutionary Biology
- ☐ Discipline-Based Education Research
- ☐ ________ Other (please specify):

What is or was the focus of your graduate training? (please select all that apply)

- ☐ Molecular/Cellular/Developmental Biology
- ☐ Physiology
- ☐ Ecology/Evolutionary Biology
- ☐ Discipline-Based Education Research
- ☐ [ ] Other (please specify):

What is the primary focus of the majority of biology courses that you teach? (please select one)

- ○ Molecular/Cellular/Developmental Biology
- ○ Physiology
- ○ Ecology/Evolutionary Biology
- ○ General Biology
- ○ [ ] Other (please specify):

In an average academic year when you are teaching, how many of your courses are at each of the following academic levels?

- [0] Non-Majors Lower-Level (100-200 level)
- [0] Majors Lower-Level (100-200 level)
- [0] Upper-Level (300-400 level)
- [0] Graduate-Level (500+ level)

How familiar are you with the *Vision and Change* report issued by the AAAS in 2011?

- ○ Extremely Familiar
- ○ Very Familiar
- ○ Somewhat Familiar
- ○ Slightly Familiar
- ○ Not At All Familiar

Have you previously provided feedback on the BioSkills Guide?

- ○ Yes
- ○ No

Please indicate your interest in any of the following forms of follow-up communication (please select all the apply)

- ☐ I would like to be sent a letter documenting my participation in this biology education activity for my records.
- ☐ I would like to be sent a copy of the final version of the BioSkills Guide once it is ready.

☐ It would be OK if you contacted me in the future to follow up on my answers to this survey.

If you checked any of the options in the preceding question, please enter your email address.

Optional: Please share any final comments you have about this survey or the BioSkills guide in general.

We are looking for more reviewers! If you have colleagues who would be interested in participating, we would be grateful if you shared this survey link with them:

bit.ly/BiologySkillsSurvey

Alternatively, you may enter their contact info (name and/or email address) and we will send them an invitation.

Powered by Qualtrics

**Supplemental Material 7. BioSkills validation phase questionnaire.**

Questionnaire used during national validation survey. Questionnaire for pilot validation was identical, except for wording of one learning outcome.

### BioSkills: Core Competency Learning Outcomes

**Welcome Page**

Based on the recommendations of "Vision and Change in Undergraduate Biology Education: A Call to Action", this NSF-supported project is intended to use the perspectives and priorities of a wide range of biology educators to unpack six "core competencies" (listed below) into measurable learning outcomes. Once completed, these learning outcomes will be made available to the community as a resource for planning and assessment of skills training in undergraduate biology. To date, we have used feedback from over 200 biology educators to develop and iteratively revise this set of learning outcomes, which we're collectively calling the "BioSkills Guide". To determine if the BioSkills Guide has broad support, we are asking a range of biology educators to rate the importance of the outcomes for a graduating general biology major.

**Vision and Change Core Competencies**

- **Process of Science**
- **Quantitative Reasoning**
- **Modeling & Simulation**
- **Interdisciplinary Nature of Science**
- **Communication & Collaboration**
- **Science & Society**

Thank you in advance for being a part of this work. The survey is expected to take ~15 minutes, and you can leave and return to the survey as needed until February 11 (your progress will save in your browser). Within the survey, you will find multiple links to download a copy of the BioSkills Guide for your personal use. If you have any questions about the project or this survey, please do not hesitate to contact me, Dr. Alexa Clemmons, or Dr. Alison Crowe.

**Contact information:**

Alexa Clemmons, Ph.D. (project lead)
Postdoctoral Research Associate
Biology Education Research Group
Department of Biology
University of Washington


Alison Crowe, Ph.D.
Principal Lecturer
Biology Education Research Group
Department of Biology
University of Washington



Have you ever served as the instructor of record for a college-level life sciences course?

- ○ Yes
- ○ No

**Screened Out**

Thank you for your interest in the BioSkills Guide! At this time we are soliciting feedback from college biology instructors, but in the future we plan to widen our scope.

If you would be interested in providing feedback then, please enter your email address. If you enter your email address we will also notify you when the BioSkills Guide is ready for distribution.

**Instructions**

#### Instructions

**You will be asked to review two or three randomly assigned core competencies,** although we would love your feedback on additional competencies if you have the time**.** You do **not** need to have experience teaching the particular core competencies you are rating.

In the current draft of the BioSkills Guide, each core competency contains multiple **program**-level learning outcomes. In addition, each program-level learning outcome contains multiple **course**-level learning outcomes. Throughout the survey, you will switch between evaluating program- and course-level outcomes. We will use the figure below to remind you of this structure and cue when the survey is switching between levels.

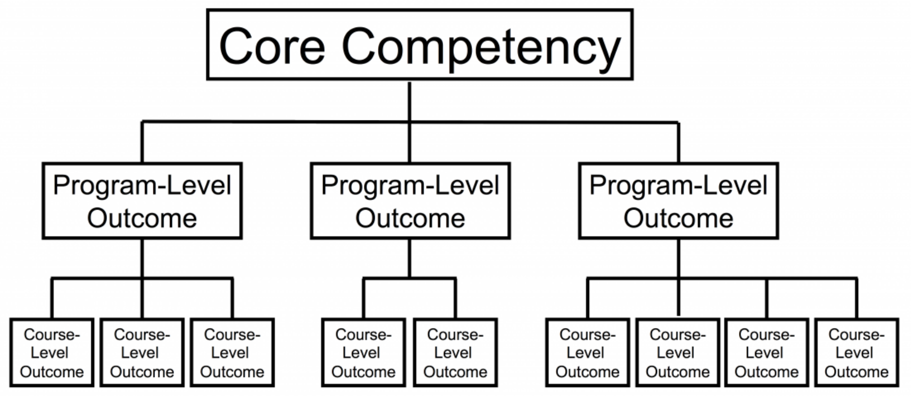


**Important note:** We intend for the BioSkills Guide to contain the learning outcomes that we, as a community, think all graduating general biology majors should achieve. **Therefore, please rate the outcomes based on whether they are important and reasonable to accomplish <u>over the course of a four-year general biology program</u>**, not introductory courses alone (1-2 years only) or in a graduate program (5+ years). Additionally, please evaluate the outcomes independently, not relative to one another (i.e. you are rating not ranking the outcomes).

#### [***Following the instructions, blocks of questions (each block corresponding to one of the six core competencies) were randomly assigned. All respondents were given 3 blocks of questions, with the option to complete additional.***]

**Process of Science**

In this portion of the survey, we would like you to rate the importance of learning outcomes for one particular core competency: **Process of Science.** Please rate the outcomes based on whether they are important to accomplish over the course of a **four-year general biology program**. Additionally, please evaluate the outcomes independently, not relative to one another.

If you would like to see the entire BioSkills Guide for context, click here. Please do not share this draft with others.

#### Process of Science

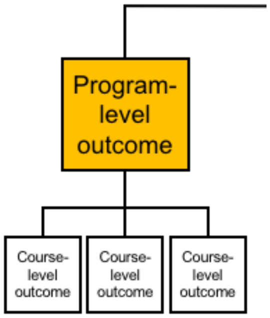


How important or unimportant is it for graduating general biology majors to achieve the following **program-level** outcome?

| | Very Unimportant | Unimportant | Neither Important Nor Unimportant | Important | Very Important |
|---|---|---|---|---|---|
| Scientific Thinking: Explain how science generates knowledge of the natural world. | O | O | O | O | O |

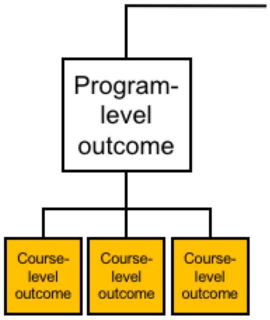


Each of the following course-level outcomes are classified under the **Scientific Thinking** program-level outcome.

How important or unimportant is it for graduating general biology majors to achieve the following **course-level** outcomes?

| | Very Unimportant | Unimportant | Neither Important Nor Unimportant | Important | Very Important |
|---|---|---|---|---|---|
| Explain how scientists use inference and evidence-based reasoning to generate knowledge. | O | O | O | O | O |
| Describe the iterative nature of science and how new evidence can lead to the revision of scientific knowledge. | O | O | O | O | O |

○ Click here if you would like to comment on the content or wording of the above outcomes.

Please share any feedback you have about the content or wording of these outcomes.

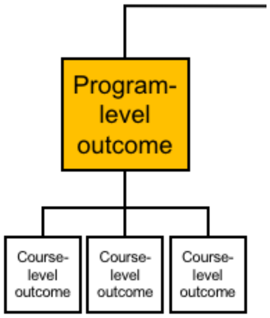


How important or unimportant is it for graduating general biology majors to achieve the following **program-level** outcome?

| | Very Unimportant | Unimportant | Neither Important Nor Unimportant | Important | Very Important |
|---|---|---|---|---|---|
| Information Literacy: Locate, interpret, and evaluate scientific information. | ○ | ○ | ○ | ○ | ○ |

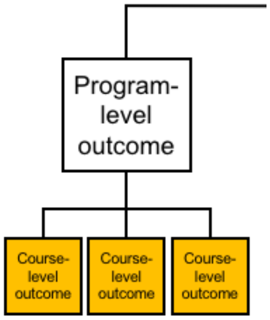


Each of the following course-level outcomes are classified under the **Information Literacy** program-level outcome.

How important or unimportant is it for graduating general biology majors to achieve the following **course-level** outcomes?

| | Very Unimportant | Unimportant | Neither Important Nor Unimportant | Important | Very Important |
|---|---|---|---|---|---|

| | Very Unimportant | Unimportant | Neither Important Nor Unimportant | Important | Very Important |
|---|---|---|---|---|---|
| Find and evaluate the credibility of a variety of sources of scientific information, including popular science media and scientific journals. | O | O | O | O | O |
| Interpret, summarize, and evaluate evidence in primary literature. | O | O | O | O | O |
| Evaluate claims in scientific papers, popular science media, and other sources using evidence-based reasoning. | O | O | O | O | O |

O Click here if you would like to comment on the content or wording of the above outcomes.

Please share any feedback you have about the content or wording of these outcomes.

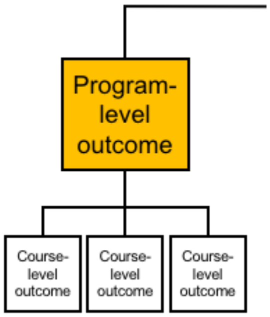


How important or unimportant is it for graduating general biology majors to achieve the following **program-level** outcome?

| | Very Unimportant | Unimportant | Neither Important Nor Unimportant | Important | Very Important |
|---|---|---|---|---|---|
| Question Formulation: Pose testable questions and hypotheses to address gaps in knowledge. | O | O | O | O | O |

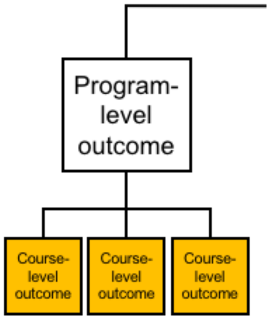


Each of the following course-level outcomes are classified under the **Question Formulation** program-level outcome.

How important or unimportant is it for graduating general biology majors to achieve the following **course-level** outcomes?

| | Very Unimportant | Unimportant | Neither Important Nor Unimportant | Important | Very Important |
|---|---|---|---|---|---|
| Recognize gaps in our current understanding of a biological system or process and identify what specific information is missing. | ○ | ○ | ○ | ○ | ○ |
| Develop research questions based on your own or others' observations. | ○ | ○ | ○ | ○ | ○ |
| Formulate testable hypotheses and state their predictions. | ○ | ○ | ○ | ○ | ○ |

○ Click here if you would like to comment on the content or wording of the above outcomes.

Please share any feedback you have about the content or wording of these outcomes.

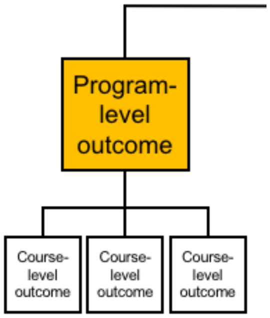


How important or unimportant is it for graduating general biology majors to achieve the following **program-level** outcome?

| | Very Unimportant | Unimportant | Neither Important Nor Unimportant | Important | Very Important |
|---|---|---|---|---|---|
| Study Design: Plan, evaluate, and implement scientific investigations. | ○ | ○ | ○ | ○ | ○ |

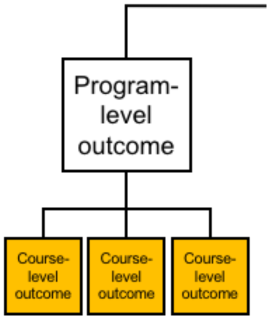


Each of the following course-level outcomes are classified under the **Study Design** program-level outcome.

How important or unimportant is it for graduating general biology majors to achieve the following **course-level** outcomes?

| | Very Unimportant | Unimportant | Neither Important Nor Unimportant | Important | Very Important |
|---|---|---|---|---|---|
| Compare the strengths and limitations of various study designs. | ○ | ○ | ○ | ○ | ○ |
| Design controlled experiments, including plans for analyzing the data. | ○ | ○ | ○ | ○ | ○ |
| Execute protocols and accurately record measurements and observations. | ○ | ○ | ○ | ○ | ○ |
| Identify methodological problems and suggest how to troubleshoot them. | ○ | ○ | ○ | ○ | ○ |
| Evaluate and suggest best practices for responsible research conduct (e.g., lab safety, record keeping, proper citation of sources). | ○ | ○ | ○ | ○ | ○ |

○ Click here if you would like to comment on the content or wording of the above outcomes.

Please share any feedback you have about the content or wording of these outcomes.

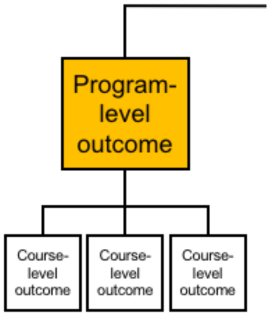


How important or unimportant is it for graduating general biology majors to achieve the following **program-level** outcome?

| | Very Unimportant | Unimportant | Neither Important Nor Unimportant | Important | Very Important |
|---|---|---|---|---|---|
| Data Interpretation & Evaluation: Interpret, evaluate, and draw conclusions from data in order to make evidence-based arguments about the natural world. | O | O | O | O | O |

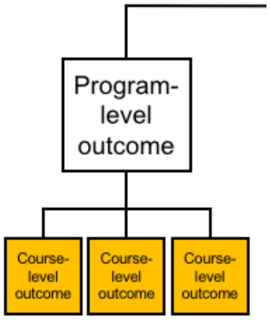


Each of the following course-level outcomes are classified under the **Data Interpretation & Evaluation** program-level outcome.

How important or unimportant is it for graduating general biology majors to achieve the following **course-level** outcomes?

| | Very Unimportant | Unimportant | Neither Important Nor Unimportant | Important | Very Important |
|---|---|---|---|---|---|
| Analyze data, summarize resulting patterns, and draw appropriate conclusions. | O | O | O | O | O |
| Describe sources of error and uncertainty in data. | O | O | O | O | O |
| Make evidence-based arguments using your own and others' findings. | O | O | O | O | O |

| | Very Unimportant | Unimportant | Neither Important Nor Unimportant | Important | Very Important |
| --- | --- | --- | --- | --- | --- |
| Relate conclusions to original hypothesis, consider alternative hypotheses, and suggest future research directions based on findings. | O | O | O | O | O |

O Click here if you would like to comment on the content or wording of the above outcomes.

Please share any feedback you have about the content or wording of these outcomes.

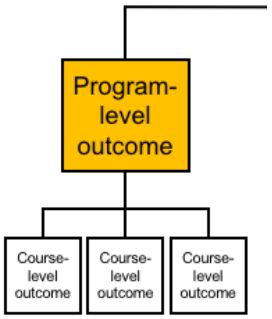


How important or unimportant is it for graduating general biology majors to achieve the following **program-level** outcome?

| | Very Unimportant | Unimportant | Neither Important Nor Unimportant | Important | Very Important |
| --- | --- | --- | --- | --- | --- |
| Doing Research: Apply science process skills to address a research question in a course-based or independent research experience. | O | O | O | O | O |

O Click here if you would like to comment on the content or wording of the above outcomes.

Please share any feedback you have about the content or wording of these outcomes.

#### Process of Science

You have now reviewed all of the program-level and course-level learning outcomes (displayed below) for this core competency.

| PROCESS OF SCIENCE | |
|---|---|
| **Program-Level Learning Outcomes** | **Course-Level Learning Outcomes** |
| **SCIENTIFIC THINKING** Explain how science generates knowledge of the natural world. | Explain how scientists use inference and evidence-based reasoning to generate knowledge. |
| | Describe the iterative nature of science and how new evidence can lead to the revision of scientific knowledge. |
| **INFORMATION LITERACY** Locate, interpret, and evaluate scientific information. | Find and evaluate the credibility of a variety of sources of scientific information, including popular science media and scientific journals. |
| | Interpret, summarize, and evaluate evidence in primary literature. |
| | Evaluate claims in scientific papers, popular science media, and other sources using evidence-based reasoning. |
| **QUESTION FORMULATION** Pose testable questions and hypotheses to address gaps in knowledge. | Recognize gaps in our current understanding of a biological system or process and identify what specific information is missing. |
| | Develop research questions based on your own or others' observations. |
| | Formulate testable hypotheses and state their predictions. |
| **STUDY DESIGN** Plan, evaluate, and implement scientific investigations. | Compare the strengths and limitations of various study designs. |
| | Design controlled experiments, including plans for analyzing the data. |
| | Execute protocols and accurately record measurements and observations. |
| | Identify methodological problems and suggest how to troubleshoot them. |
| | Evaluate and suggest best practices for responsible research conduct (e.g., lab safety, record keeping, proper citation of sources). |
| **DATA INTERPRETATION & EVALUATION** Interpret, evaluate, and draw conclusions from data in order to make evidence-based arguments about the natural world. | Analyze data, summarize resulting patterns, and draw appropriate conclusions. |
| | Describe sources of error and uncertainty in data. |
| | Make evidence-based arguments using your own and others' findings. |
| | Relate conclusions to original hypothesis, consider alternative hypotheses, and suggest future research directions based on findings. |
| **DOING RESEARCH** Apply science process skills to address a research question in a course-based or independent research experience. | |

○ Optional: Given this review, please click here if you believe there are essential learning outcomes missing from the Process of Science core competency.

Please share any essential learning outcomes you believe are missing from this core competency.

Optional: Please share any other feedback on the **Process of Science** core competency.

**Quantitative Reasoning**

In this portion of the survey, we would like you to rate the importance of learning outcomes for one particular core competency: **Quantitative Reasoning.** Please rate the outcomes based on whether they are important to accomplish over the course of a **four-year general biology program**. Additionally, please evaluate the outcomes independently, not relative to one another.

If you would like to see the entire BioSkills Guide for context, click here. Please do not share this draft with others.

#### Quantitative Reasoning

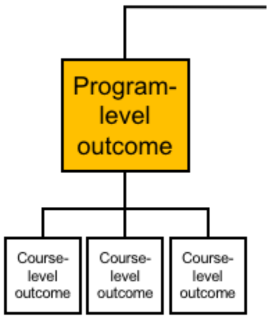


How important or unimportant is it for graduating general biology majors to achieve the following **program-level** outcome?

| | Very Unimportant | Unimportant | Neither Important Nor Unimportant | Important | Very Important |
|---|---|---|---|---|---|
| Numeracy: Use basic mathematics (e.g., algebra, probability, unit conversions) in biological contexts. | O | O | O | O | O |

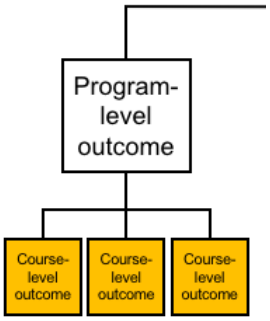


Each of the following course-level outcomes are classified under the **Numeracy** program-level outcome.

How important or unimportant is it for graduating general biology majors to achieve the following **course-level** outcomes?

| | Very Unimportant | Unimportant | Neither Important Nor Unimportant | Important | Very Important |
|---|---|---|---|---|---|
| Perform basic calculations (e.g., percentages, frequencies, rates, means). | O | O | O | O | O |
| Select and apply appropriate equations (e.g., Hardy-Weinberg, Nernst, Gibbs free energy) to solve problems. | O | O | O | O | O |

| | Very Unimportant | Unimportant | Neither Important Nor Unimportant | Important | Very Important |
|---|---|---|---|---|---|
| Interpret and manipulate mathematical relationships (e.g., scale, ratios, units) to make quantitative comparisons. | O | O | O | O | O |
| Use probability and understanding of biological variability to reason about biological processes and statistical analyses. | O | O | O | O | O |
| Use rough estimates informed by biological knowledge to check quantitative work. | O | O | O | O | O |
| Describe how quantitative reasoning helps biologists understand the natural world. | O | O | O | O | O |

O Click here if you would like to comment on the content or wording of the above outcomes.

Please share any feedback you have about the content or wording of these outcomes.

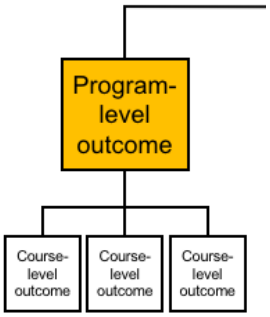


How important or unimportant is it for graduating general biology majors to achieve the following **program-level** outcome?

| | Very Unimportant | Unimportant | Neither Important Nor Unimportant | Important | Very Important |
|---|---|---|---|---|---|
| Quantitative & Computational Data Analysis: Apply the tools of graphing, statistics, and data science to analyze biological data. | O | O | O | O | O |

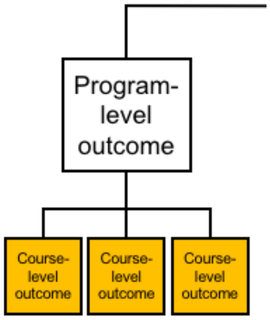


Each of the following course-level outcomes are classified under the **Quantitative & Computational Data Analysis** program-level outcome.

How important or unimportant is it for graduating general biology majors to achieve the following **course-level** outcomes?

| | Very Unimportant | Unimportant | Neither Important Nor Unimportant | Important | Very Important |
|---|---|---|---|---|---|
| Record, organize, and annotate simple data sets. | ○ | ○ | ○ | ○ | ○ |
| Create and interpret informative graphs and other data visualizations. | ○ | ○ | ○ | ○ | ○ |
| Select, carry out, and interpret statistical analyses. | ○ | ○ | ○ | ○ | ○ |
| Describe how biologists answer research questions using databases, large data sets, and data science tools. | ○ | ○ | ○ | ○ | ○ |
| Interpret the biological meaning of quantitative results. | ○ | ○ | ○ | ○ | ○ |

○ Click here if you would like to comment on the content or wording of the above outcomes.

Please share any feedback you have about the content or wording of these outcomes.

##### Quantitative Reasoning

You have now reviewed all of the program-level and course-level learning outcomes (displayed below) for this core competency.

| QUANTITATIVE REASONING | |
|---|---|
| Program-Level Learning Outcomes | Course-Level Learning Outcomes |
| **NUMERACY**<br>Use basic mathematics (e.g., algebra, probability, unit conversions) in biological contexts. | Perform basic calculations (e.g., percentages, frequencies, rates, means). |
| | Select and apply appropriate equations (e.g., Hardy-Weinberg, Nernst, Gibbs free energy) to solve problems. |
| | Interpret and manipulate mathematical relationships (e.g., scale, ratios, units) to make quantitative comparisons. |
| | Use probability and understanding of biological variability to reason about biological processes and statistical analyses. |
| | Use rough estimates informed by biological knowledge to check quantitative work. |
| | Describe how quantitative reasoning helps biologists understand the natural world. |
| **QUANTITATIVE & COMPUTATIONAL DATA ANALYSIS**<br>Apply the tools of graphing, statistics, and data science to analyze biological data. | Record, organize, and annotate simple data sets. |
| | Create and interpret informative graphs and other data visualizations. |
| | Select, carry out, and interpret statistical analyses. |
| | Describe how biologists answer research questions using databases, large data sets, and data science tools. |
| | Interpret the biological meaning of quantitative results. |

○ Optional: Given this review, please click here if you believe there are essential learning outcomes missing from the Quantitative Reasoning core competency.

Please share any essential learning outcomes you believe are missing from this core competency.

Optional: Please share any other feedback on the **Quantitative Reasoning** core competency.

**Modeling & Simulation**

In this portion of the survey, we would like you to rate the importance of learning outcomes for one particular core competency: **Modeling & Simulation.** Please rate the outcomes based on whether they are important to accomplish over the course of a **four-year general biology program**. Additionally, please evaluate the outcomes independently, not relative to one another.

If you would like to see the entire BioSkills Guide for context, click here. Please do not share this draft with others.

#### Modeling & Simulation

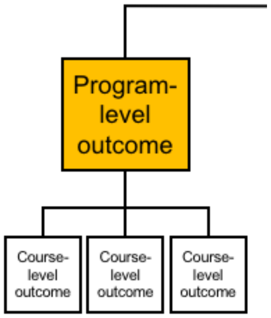

How important or unimportant is it for graduating general biology majors to achieve the following **program-level** outcome?

| | Very Unimportant | Unimportant | Neither Important Nor Unimportant | Important | Very Important |
|---|---|---|---|---|---|
| Purpose of Models: Recognize the important roles that scientific models, of many different types (conceptual, mathematical, physical, etc.), play in predicting and communicating biological phenomena. | O | O | O | O | O |

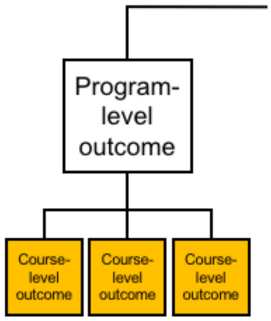


Each of the following course-level outcomes are classified under the **Purpose of Models** program-level outcome.

How important or unimportant is it for graduating general biology majors to achieve the following **course-level** outcomes?

| | Very Unimportant | Unimportant | Neither Important Nor Unimportant | Important | Very Important |
|---|---|---|---|---|---|
| Describe why biologists use simplified representations (models) when solving problems and communicating ideas. | O | O | O | O | O |
| Given two models of the same biological process or system, compare their strengths, limitations, and assumptions. | O | O | O | O | O |

O Click here if you would like to comment on the content or wording of the above outcomes.

Please share any feedback you have about the content or wording of these outcomes.

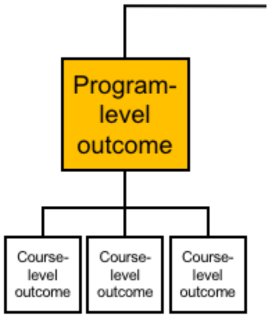


How important or unimportant is it for graduating general biology majors to achieve the following **program-level** outcome?

| | Very Unimportant | Unimportant | Neither Important Nor Unimportant | Important | Very Important |
|---|---|---|---|---|---|
| Model Application: Make inferences and solve problems using models and simulations. | O | O | O | O | O |

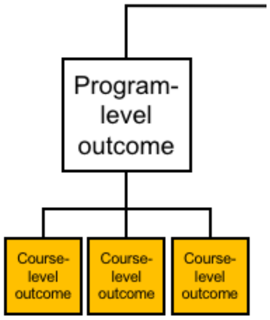


Each of the following course-level outcomes are classified under the **Model Application** program-level outcome.

How important or unimportant is it for graduating general biology majors to achieve the following **course-level** outcomes?

| | Very Unimportant | Unimportant | Neither Important Nor Unimportant | Important | Very Important |
|---|---|---|---|---|---|
| Summarize relationships and trends that can be inferred from a given model or simulation. | O | O | O | O | O |
| Use models and simulations to make predictions and refine hypotheses. | O | O | O | O | O |

O Click here if you would like to comment on the content or wording of the above outcomes.

Please share any feedback you have about the content or wording of these outcomes.

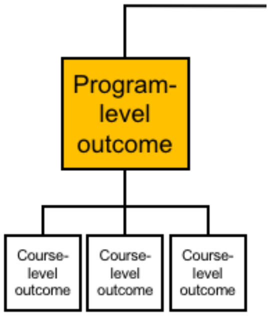


How important or unimportant is it for graduating general biology majors to achieve the following **program-level** outcome?

| | Very Unimportant | Unimportant | Neither Important Nor Unimportant | Important | Very Important |
|---|---|---|---|---|---|
| Modeling: Build and evaluate models of biological systems. | | | | | |

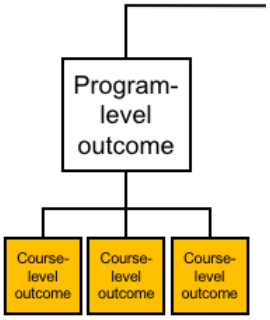


Each of the following course-level outcomes are classified under the **Modeling** program-level outcome.

How important or unimportant is it for graduating general biology majors to achieve the following **course-level** outcomes?

| | Very Unimportant | Unimportant | Neither Important Nor Unimportant | Important | Very Important |
|---|---|---|---|---|---|
| Build and revise conceptual models (e.g., diagrams, concept maps, flow charts) to propose how a biological system or process works. | | | | | |
| Identify important components of a system and describe how they influence each other (e.g., positively or negatively). | | | | | |

| | Very Unimportant | Unimportant | Neither Important Nor Unimportant | Important | Very Important |
|---|---|---|---|---|---|
| Evaluate conceptual, mathematical, or computational models by comparing their predictions with empirical data. | ○ | ○ | ○ | ○ | ○ |

○ Click here if you would like to comment on the content or wording of the above outcomes.

Please share any feedback you have about the content or wording of these outcomes.

#### Modeling & Simulation

You have now reviewed all of the program-level and course-level learning outcomes (displayed below) for this core competency.

| MODELING & SIMULATION | |
|---|---|
| **Program-Level Learning Outcomes** | **Course-Level Learning Outcomes** |
| **PURPOSE OF MODELS**<br>Recognize the important roles that scientific models, of many different types (conceptual, mathematical, physical, etc.), play in predicting and communicating biological phenomena. | Describe why biologists use simplified representations (models) when solving problems and communicating ideas. |
| | Given two models of the same biological process or system, compare their strengths, limitations, and assumptions. |
| **MODEL APPLICATION**<br>Make inferences and solve problems using models and simulations. | Summarize relationships and trends that can be inferred from a given model or simulation. |
| | Use models and simulations to make predictions and refine hypotheses. |
| **MODELING**<br>Build and evaluate models of biological systems. | Build and revise conceptual models (e.g., diagrams, concept maps, flow charts) to propose how a biological system or process works. |
| | Identify important components of a system and describe how they influence each other (e.g., positively or negatively). |
| | Evaluate conceptual, mathematical, or computational models by comparing their predictions with empirical data. |

○ Optional: Given this review, please click here if you believe there are essential learning outcomes missing from the Modeling & Simulation core competency.

Please share any essential learning outcomes you believe are missing from this core competency.

Optional: Please share any other feedback on the **Modeling & Simulation** core competency.

**Option to Continue**

Thank you for all of your feedback so far! We know that your time is valuable. We would love your feedback on additional outcomes, if you have the time.

Would you like to evaluate another set of outcomes?

### [***This question was shown after first 3 randomly assigned blocks of questions, and then subsequently after each additional block of questions until all 6 blocks were complete.***]

- Yes
- No

**Interdisciplinary Nature of Science**

In this portion of the survey, we would like you to rate the importance of learning outcomes for one particular core competency: **Interdisciplinary Nature of Science.** Please rate the outcomes based on whether they are important to accomplish over the course of a **four-year general biology program**. Additionally, please evaluate the outcomes independently, not relative to one another.

If you would like to see the entire BioSkills Guide for context, click here. Please do not share this draft with others.

#### Interdisciplinary Nature of Science

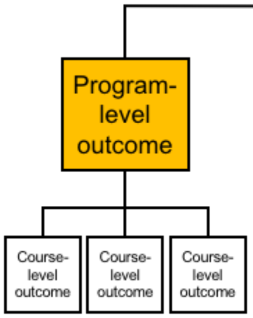


How important or unimportant is it for graduating general biology majors to achieve the following **program-level** outcome?

| | Very Unimportant | Unimportant | Neither Important Nor Unimportant | Important | Very Important |
|---|---|---|---|---|---|
| Connecting Scientific Knowledge: Integrate concepts across other STEM disciplines (e.g., chemistry, physics) and multiple fields of biology (e.g., cell biology, ecology). | O | O | O | O | O |

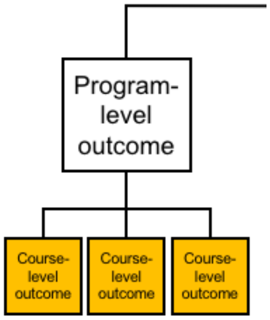


Each of the following course-level outcomes are classified under the **Connecting Scientific Knowledge** program-level outcome.

How important or unimportant is it for graduating general biology majors to achieve the following **course-level** outcomes?

| | Very Unimportant | Unimportant | Neither Important Nor Unimportant | Important | Very Important |
|---|---|---|---|---|---|
| Given a biological problem, identify relevant concepts from other STEM disciplines or fields of biology. | O | O | O | O | O |
| Build models or explanations of simple biological processes that include concepts from other STEM disciplines or multiple fields of biology. | O | O | O | O | O |

O Click here if you would like to comment on the content or wording of the above outcomes.

Please share any feedback you have about the content or wording of these outcomes.

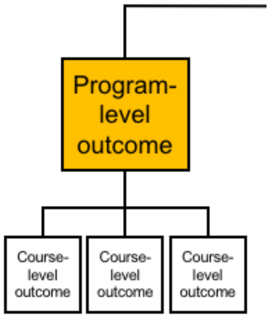


How important or unimportant is it for graduating general biology majors to achieve the following **program-level** outcome?

| | Very Unimportant | Unimportant | Neither Important Nor Unimportant | Important | Very Important |
|---|---|---|---|---|---|
| Interdisciplinary Problem Solving: Consider interdisciplinary solutions to real-world problems. | O | O | O | O | O |

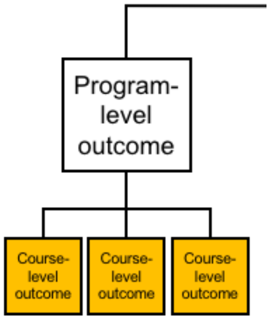


Each of the following course-level outcomes are classified under the **Interdisciplinary Problem Solving** program-level outcome.

How important or unimportant is it for graduating general biology majors to achieve the following **course-level** outcomes?

| | Very Unimportant | Unimportant | Neither Important Nor Unimportant | Important | Very Important |
|---|---|---|---|---|---|
| Describe examples of real-world problems that are too complex to be solved by applying biological approaches alone. | O | O | O | O | O |
| Suggest how collaborators in STEM and non-STEM disciplines could contribute to solutions of real-world problems. | O | O | O | O | O |
| Be able to explain biological concepts, data, and methods, including their limitations, using language understandable by collaborators in other disciplines. | O | O | O | O | O |

O Click here if you would like to comment on the content or wording of the above outcomes.

Please share any feedback you have about the content or wording of these outcomes.

**Interdisciplinary Nature of Science**

You have now reviewed all of the program-level and course-level learning outcomes (displayed below) for this core competency.

| INTERDISCIPLINARY NATURE OF SCIENCE | |
|---|---|
| Program-Level Learning Outcomes | Course-Level Learning Outcomes |
| **CONNECTING SCIENTIFIC KNOWLEDGE** Integrate concepts across other STEM disciplines (e.g., chemistry, physics) and multiple fields of biology (e.g., cell biology, ecology). | Given a biological problem, identify relevant concepts from other STEM disciplines or fields of biology. |
| | Build models or explanations of simple biological processes that include concepts from other STEM disciplines or multiple fields of biology. |
| **INTERDISCIPLINARY PROBLEM SOLVING** Consider interdisciplinary solutions to real-world problems. | Describe examples of real-world problems that are too complex to be solved by applying biological approaches alone. |
| | Suggest how collaborators in STEM and non-STEM disciplines could contribute to solutions of real-world problems. |
| | Be able to explain biological concepts, data, and methods, including their limitations, using language understandable by collaborators in other disciplines. |

○ Optional: Given this review, please click here if you believe there are essential learning outcomes missing from the Interdisciplinary Nature of Science core competency.

Please share any essential learning outcomes you believe are missing from this core competency.

Optional: Please share any other feedback on the **Interdisciplinary Nature of Science** core competency.

**Communication & Collaboration**

In this portion of the survey, we would like you to rate the importance of learning outcomes for one particular core competency: **Communication & Collaboration.** Please rate the outcomes based on whether they are important to accomplish over the course of a **four-year general biology program**. Additionally, please evaluate the outcomes independently, not relative to one another.

If you would like to see the entire BioSkills Guide for context, click here. Please do not share this draft with others.

#### Communication & Collaboration

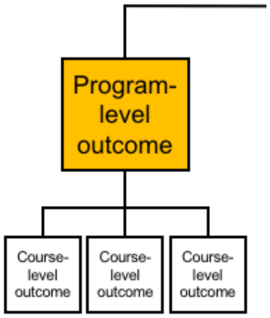


How important or unimportant is it for graduating general biology majors to achieve the following **program-level** outcome?

| | Very Unimportant | Unimportant | Neither Important Nor Unimportant | Important | Very Important |
|---|---|---|---|---|---|
| Communication: Share ideas, data, and findings with others clearly and accurately. | O | O | O | O | O |

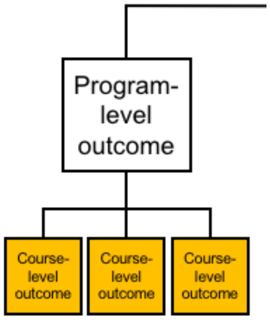


Each of the following course-level outcomes are classified under the **Communication** program-level outcome.

How important or unimportant is it for graduating general biology majors to achieve the following **course-level** outcomes?

| | Very Unimportant | Unimportant | Neither Important Nor Unimportant | Important | Very Important |
|---|---|---|---|---|---|
| Use appropriate language and style to communicate science effectively to targeted audiences (e.g., general public, biology experts, collaborators in other disciplines). | O | O | O | O | O |
| Use a variety of modes to communicate science (e.g., oral, written, visual). | O | O | O | O | O |

O Click here if you would like to comment on the content or wording of the above outcomes.

Please share any feedback you have about the content or wording of these outcomes.

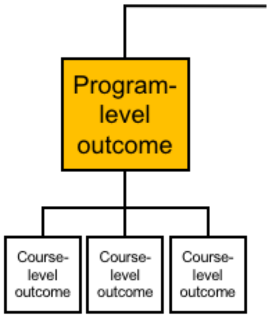


How important or unimportant is it for graduating general biology majors to achieve the following **program-level** outcome?

| | Very Unimportant | Unimportant | Neither Important Nor Unimportant | Important | Very Important |
|---|---|---|---|---|---|
| Collaboration: Work productively in teams with people who have diverse backgrounds, skill sets, and perspectives. | O | O | O | O | O |

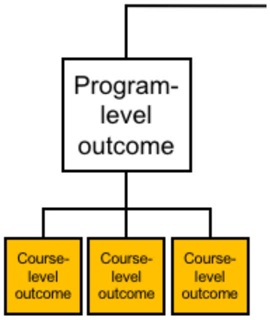


Each of the following course-level outcomes are classified under the **Collaboration** program-level outcome.

How important or unimportant is it for graduating general biology majors to achieve the following **course-level** outcomes?

| | Very Unimportant | Unimportant | Neither Important Nor Unimportant | Important | Very Important |
|---|---|---|---|---|---|
| Work with teammates to establish and periodically update group plans and expectations (e.g., team goals, project timeline, rules for group interactions, individual and collaborative tasks). | O | O | O | O | O |

| | Very Unimportant | Unimportant | Neither Important Nor Unimportant | Important | Very Important |
|---|---|---|---|---|---|
| Elicit, listen to, and incorporate ideas from teammates with different perspectives and backgrounds. | ○ | ○ | ○ | ○ | ○ |
| Work effectively with teammates to complete projects. | ○ | ○ | ○ | ○ | ○ |

○ Click here if you would like to comment on the content or wording of the above outcomes.

Please share any feedback you have about the content or wording of these outcomes.

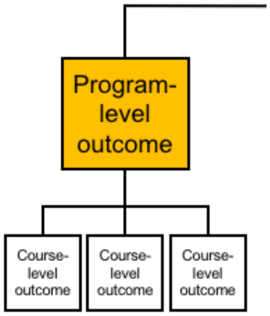


How important or unimportant is it for graduating general biology majors to achieve the following **program-level** outcome?

| | Very Unimportant | Unimportant | Neither Important Nor Unimportant | Important | Very Important |
|---|---|---|---|---|---|
| Collegial Review: Provide and respond to constructive feedback in order to improve individual and team work. | ○ | ○ | ○ | ○ | ○ |

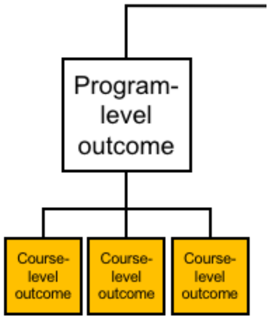

Each of the following course-level outcomes are classified under the **Collegial Review** program-level outcome.

How important or unimportant is it for graduating general biology majors to achieve the following **course-level** outcomes?

| | Very Unimportant | Unimportant | Neither Important Nor Unimportant | Important | Very Important |
|---|---|---|---|---|---|
| Evaluate feedback from others and revise work or behavior appropriately. | ○ | ○ | ○ | ○ | ○ |
| Critique others' work and ideas constructively and respectfully. | ○ | ○ | ○ | ○ | ○ |

○ Click here if you would like to comment on the content or wording of the above outcomes.

Please share any feedback you have about the content or wording of these outcomes.

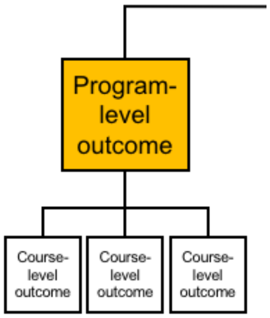


How important or unimportant is it for graduating general biology majors to achieve the following **program-level** outcome?

| | Very Unimportant | Unimportant | Neither Important Nor Unimportant | Important | Very Important |
|---|---|---|---|---|---|
| Metacognition: Reflect on your own learning, performance, and achievements. | ○ | ○ | ○ | ○ | ○ |

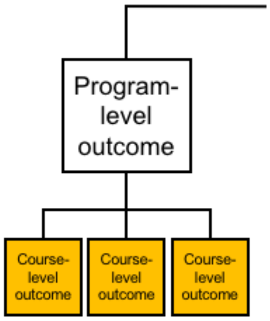


Each of the following course-level outcomes are classified under the **Metacognition** program-level outcome.

How important or unimportant is it for graduating general biology majors to achieve the following **course-level** outcomes?

| | Very Unimportant | Unimportant | Neither Important Nor Unimportant | Important | Very Important |
|---|---|---|---|---|---|
| Evaluate your own understanding and skill level. | O | O | O | O | O |
| Assess personal progress and contributions to your team and generate a plan to change your behavior as needed. | O | O | O | O | O |

O Click here if you would like to comment on the content or wording of the above outcomes.

Please share any feedback you have about the content or wording of these outcomes.

#### Communication & Collaboration

You have now reviewed all of the program-level and course-level learning outcomes (displayed below) for this core competency.

| COMMUNICATION & COLLABORATION | |
|---|---|
| **Program-Level Learning Outcomes** | **Course-Level Learning Outcomes** |
| **COMMUNICATION** Share ideas, data, and findings with others clearly and accurately. | Use appropriate language and style to communicate science effectively to targeted audiences (e.g., general public, biology experts, collaborators in other disciplines). |
| | Use a variety of modes to communicate science (e.g., oral, written, visual). |
| **COLLABORATION** Work productively in teams with people who have diverse backgrounds, skill sets, and perspectives. | Work with teammates to establish and periodically update group plans and expectations (e.g., team goals, project timeline, rules for group interactions, individual and collaborative tasks). |
| | Elicit, listen to, and incorporate ideas from teammates with different perspectives and backgrounds. |
| | Work effectively with teammates to complete projects. |
| **COLLEGIAL REVIEW** Provide and respond to constructive feedback in order to improve individual and team work. | Evaluate feedback from others and revise work or behavior appropriately. |
| | Critique others' work and ideas constructively and respectfully. |
| **METACOGNITION** Reflect on your own learning, performance, and achievements. | Evaluate your own understanding and skill level. |
| | Assess personal progress and contributions to your team and generate a plan to change your behavior as needed. |

○ Optional: Given this review, please click here if you believe there are essential learning outcomes missing from the Communication & Collaboration core competency.

Please share any essential learning outcomes you believe are missing from this core competency.

Optional: Please share any other feedback on the **Communication & Collaboration** core competency.

**Science & Society**

In this portion of the survey, we would like you to rate the importance of learning outcomes for one particular core competency: **Science & Society.** Please rate the outcomes based on whether they are important to accomplish over the course of a **four-year general biology program**. Additionally, please evaluate the outcomes independently, not relative to one another.

If you would like to see the entire BioSkills Guide for context, click here. Please do not share this draft with others.

#### Science & Society

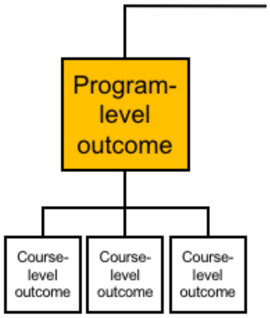


How important or unimportant is it for graduating general biology majors to achieve the following **program-level** outcome?

| | Very Unimportant | Unimportant | Neither Important Nor Unimportant | Important | Very Important |
|---|---|---|---|---|---|
| Ethics: Demonstrate the ability to critically analyze ethical issues in the conduct of science. | O | O | O | O | O |

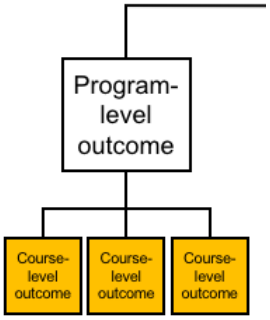


Each of the following course-level outcomes are classified under the **Ethics** program-level outcome.

How important or unimportant is it for graduating general biology majors to achieve the following **course-level** outcomes?

| | Very Unimportant | Unimportant | Neither Important Nor Unimportant | Important | Very Important |
|---|---|---|---|---|---|
| Identify and evaluate ethical considerations (e.g., use of animal or human subjects, conflicts of interest, confirmation bias) in a given research study. | O | O | O | O | O |
| Critique how ethical controversies in biological research have been and can continue to be addressed by the scientific community. | O | O | O | O | O |

O Click here if you would like to comment on the content or wording of the above outcomes.

Please share any feedback you have about the content or wording of these outcomes.

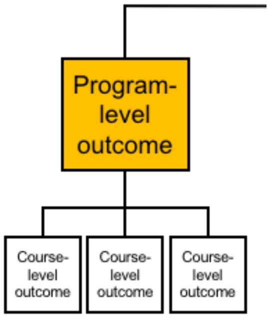


How important or unimportant is it for graduating general biology majors to achieve the following **program-level** outcome?

| | Very Unimportant | Unimportant | Neither Important Nor Unimportant | Important | Very Important |
|---|---|---|---|---|---|
| Societal Influences: Consider the potential impacts of outside influences (historical, cultural, political, technological) on how science is practiced. | ○ | ○ | ○ | ○ | ○ |

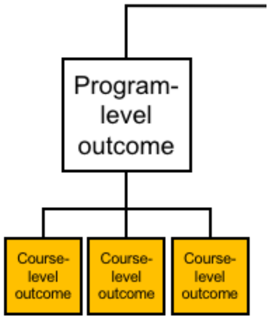


Each of the following course-level outcomes are classified under the **Societal Influences** program-level outcome.

How important or unimportant is it for graduating general biology majors to achieve the following **course-level** outcomes?

| | Very Unimportant | Unimportant | Neither Important Nor Unimportant | Important | Very Important |
|---|---|---|---|---|---|
| Describe examples of how scientists' backgrounds and biases can influence science and how science is enhanced through diversity. | ○ | ○ | ○ | ○ | ○ |

| | Very Unimportant | Unimportant | Neither Important Nor Unimportant | Important | Very Important |
|---|---|---|---|---|---|
| Identify and describe how systemic factors (e.g., socioeconomic, political) affect how and by whom science is conducted. | O | O | O | O | O |

O Click here if you would like to comment on the content or wording of the above outcomes.

Please share any feedback you have about the content or wording of these outcomes.

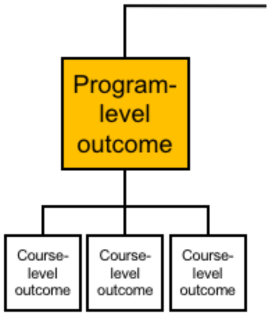


How important or unimportant is it for graduating general biology majors to achieve the following **program-level** outcome?

| | Very Unimportant | Unimportant | Neither Important Nor Unimportant | Important | Very Important |
|---|---|---|---|---|---|
| Science's Impact on Society: Apply scientific reasoning in daily life and recognize the impacts of science on a local and global scale. | O | O | O | O | O |

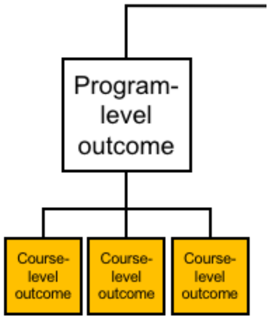


Each of the following course-level outcomes are classified under the **Science's Impact on Society** program-level outcome.

How important or unimportant is it for graduating general biology majors to achieve the following **course-level** outcomes?

| | Very Unimportant | Unimportant | Neither Important Nor Unimportant | Important | Very Important |
|---|---|---|---|---|---|
| Apply evidence-based reasoning and biological knowledge in daily life (e.g., consuming popular media, deciding how to vote). | O | O | O | O | O |
| Use examples to describe the relevance of science in everyday experiences. | O | O | O | O | O |
| Identify and describe the broader societal impacts of biological research on different stakeholders. | O | O | O | O | O |
| Describe the roles scientists have in facilitating public understanding of science. | O | O | O | O | O |

O Click here if you would like to comment on the content or wording of the above outcomes.

Please share any feedback you have about the content or wording of these outcomes.

#### Science & Society

You have now reviewed all of the program-level and course-level learning outcomes (displayed below) for this core competency.

**SCIENCE & SOCIETY**

| Program-Level Learning Outcomes | Course-Level Learning Outcomes |
|---|---|
| **ETHICS** Demonstrate the ability to critically analyze ethical issues in the conduct of science. | Identify and evaluate ethical considerations (e.g., use of animal or human subjects, conflicts of interest, confirmation bias) in a given research study. |
| | Critique how ethical controversies in biological research have been and can continue to be addressed by the scientific community. |
| **SOCIETAL INFLUENCES** Consider the potential impacts of outside influences (historical, cultural, political, technological) on how science is practiced. | Describe examples of how scientists' backgrounds and biases can influence science and how science is enhanced through diversity. |
| | Identify and describe how systemic factors (e.g., socioeconomic, political) affect how and by whom science is conducted. |
| **SCIENCE'S IMPACT ON SOCIETY** Apply scientific reasoning in daily life and recognize the impacts of science on a local and global scale. | Apply evidence-based reasoning and biological knowledge in daily life (e.g., consuming popular media, deciding how to vote). |
| | Use examples to describe the relevance of science in everyday experiences. |
| | Identify and describe the broader societal impacts of biological research on different stakeholders. |
| | Describe the roles scientists have in facilitating public understanding of science. |

- ○ Optional: Given this review, please click here if you believe there are essential learning outcomes missing from the Science & Society core competency.

Please share any essential learning outcomes you believe are missing from this core competency.

[ ]

Optional: Please share any other feedback on the **Science & Society** core competency.

[ ]

**Demographics**

#### Demographic Questions

We ask that you complete the following demographic questions so that we can determine if we have surveyed a representative population. We will not link specific responses with any individual identifying information when sharing the results of this survey.

What is the name of your current institution? (This will be used to gather additional institutional demographic information.)

[ ]

Which of the following best describes your institution type?

- ○ Associate's Degree-Granting
- ○ Bachelor's Degree-Granting
- ○ Master's Degree-Granting
- ○ Doctoral Degree-Granting
- ○ [ ] Other (please specify):

Which of the following best describes your current position?

- ○ Graduate Student
- ○ Postdoc
- ○ Lecturer or Instructor
- ○ Assistant, Associate, or Full Professor
- ○ Staff
- ○ [ ] Other (please specify):

In your current position, what is your primary responsibility?

- ○ Teaching
- ○ Research
- ○ Teaching and Research Equally
- ○ [ ] Other (please describe briefly)

What is the focus of your <u>current</u> research, if applicable? (please select all that apply)

- ☐ I am not currently engaged in research
- ☐ Molecular/Cellular/Developmental Biology
- ☐ Physiology
- ☐ Ecology/Evolutionary Biology
- ☐ Discipline-Based Education Research
- ☐ [ ] Other (please specify):

What is or was the focus of your graduate training? (please select all that apply)

- ☐ Molecular/Cellular/Developmental Biology
- ☐ Physiology
- ☐ Ecology/Evolutionary Biology
- ☐ Discipline-Based Education Research
- ☐ [ ] Other (please specify):

What is the <u>primary</u> focus of the majority of biology courses that you teach? (please select one)

- ○ Molecular/Cellular/Developmental Biology
- ○ Physiology
- ○ Ecology/Evolutionary Biology
- ○ General Biology
- ○ [ ] Other (please specify):

In an average academic year when you are teaching, how many of your courses are at each of the following academic levels?

- [0] Non-Majors Lower-Level (100-200 level)
- [0] Majors Lower-Level (100-200 level)
- [0] Upper-Level (300-400 level)
- [0] Graduate-Level (500+ level)

How familiar are you with the *Vision and Change* report issued by the AAAS in 2011?

- ○ Extremely Familiar
- ○ Very Familiar
- ○ Somewhat Familiar
- ○ Slightly Familiar
- ○ Not At All Familiar

What is your gender?

- ○ Female
- ○ Male

○ [ ] Other identity (please specify)

Have you previously provided feedback on the BioSkills Guide?

○ Yes

○ No

Would you like to be sent a copy of the final version of the BioSkills Guide once it is ready?

○ Yes

○ No

If you answered yes to the preceding question, please enter your email address.

[ ]

Optional: Please share any final comments you have about this survey or the BioSkills guide in general.

[ ]

Optional: We are looking for more participants! If you have colleagues who would be interested in participating, we would be grateful if you shared this survey link with them:

https://uwbiology.co1.qualtrics.com/jfe/form/SV_d0wvusxksI1cxtb

Alternatively, you can enter their name and email address below, and we will send them an invitation.

[ ]

Powered by Qualtrics
